## Supplementary information for "Universal probabilistic programming offers a powerful approach to statistical phylogenetics"

#### 1 Probabilistic programming: an introduction

In this section, we give a brief introduction to (universal) probabilistic programming languages (PPLs), focusing on the key constructs that are available in most PPLs. First, we give a short overview of different PPLs, followed by an introduction to three essential concepts in probabilistic programming: (i) sampling, (ii) conditioning, and (iii) inference. These concepts are illustrated here in the WebPPL language[a], which is based on a functional subset[b] of the JavaScript language.

##### 1.1 Overview

A central objective of probabilistic programming is to separate the *model* from the *inference* algorithm, such that a user can construct and use a probabilistic model without the need to implement the inference algorithm explicitly. Instead, it is the task of the runtime system of the PPL to automatically perform the inference, potentially based on some method preferences specified by the user.

Programming and modeling languages that separate the model specification from the inference algorithm have been around for several decades. One of the first of these languages is BUGS (Bayesian inference Using Gibbs Sampling)[1]. BUGS allows users to describe probabilistic graphical models[2]—in particular Bayesian networks—in a declarative way. The model parameters of interest are then estimated by automatically applying Bayesian inference using Gibbs sampling (and some other methods). More recent languages that separate modeling and inference of graphical models include Infer.NET[3].

Although the above-mentioned languages and environments have shown great success in their application areas, they have certain model restrictions. In particular, they are limited to models where the dependencies between random variables can be expressed as a Bayesian network, that is, a finite directed acyclic graph, potentially with if-then-else conditions over variables. In some domains, this is not sufficient to describe the models of interest. Rather recently, the concept of *probabilistic programming languages*[4] has gained significant attention as a promising solution, in particular within the machine learning and programming language communities. The key idea of this new paradigm is to extend Turing-complete programming languages with probabilistic operations that include, for example, the drawing of (random) samples from a given probability distribution, the conditioning of random variables on observed outcomes, and the marginalization of random variables[5,6].

Such languages are sometimes referred to as *universal probabilistic programming languages* to clearly differentiate them from languages based on Bayesian networks, which have sometimes in recent years also been included in the probabilistic programming family. Here, we will use the terms “probabilistic programming” and “probabilistic programming language (PPL)” exclusively for universal languages.

Turing-completeness is an important concept in computer science, describing how expressive a programming language is. The famous Church-Turing thesis conjectures that any function, whose value can be computed by an algorithm, can be computed by a Turing-complete programming language. For instance, PPLs make it possible to use recursion (or loops) dependent on a stochastic expression when defining probabilistic models. This means that the graphical network describing the model is *dynamic* and can change during inference due to random sampling and observed data. In a PPL, the probabilistic model *is a program*, where the inference algorithm is not part of that program (the model). Hence, an alternative and potentially more intuitive name for a probabilistic program may be a *programmatic model*: a model that is implemented as a program.

One of the earliest PPLs is Church[7], which extends a functional subset of the Scheme programming language. Other PPLs (both universal and non-universal) include Figaro[8] (a PPL embedded in Scala), WebPPL[9] (a recent PPL embedded into JavaScript), Anglican[10] (a general-purpose PPL embedded into Clojure that runs on the Java virtual machine), Venture[11] (a PPL with syntax similar to JavaScript), Edward[12] (a Python library for probabilistic modelling), Pyro[13] (a PPL built on top of

*
†F.R., J.K., and V.S. contributed equally to this work.
[a]`http://webppl.org`
[b]Functional programming (FP) is a programming paradigm, in which code is structured in units called *functions* that have no side effects; i.e. they only operate on a given input and produce an output but do not manipulate external objects.

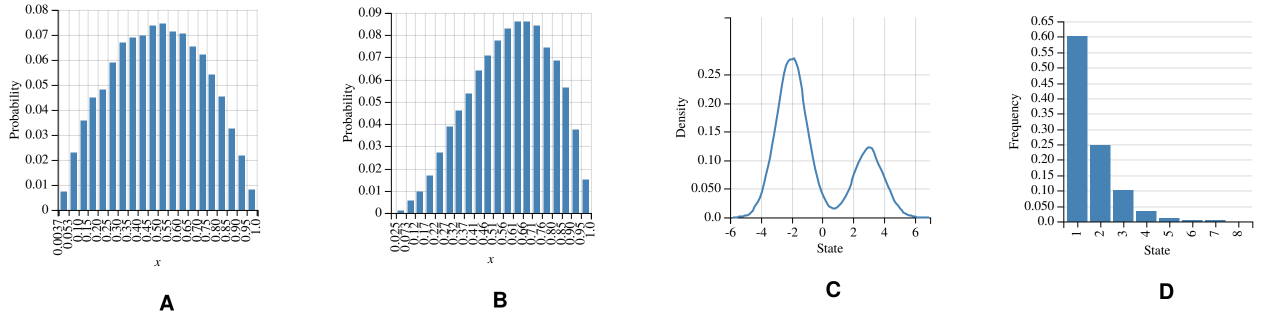


**Fig. 1 A** Prior distribution of the bias for the coin flip example. **B** Posterior distribution of the bias after observing heads once. **C** Mixture model of two Gaussian models mixed using stochastic branching. **D** Resulting geometric distribution with $p = 0.6$. All plots are generated using the WebPPL environment.

PyTorch), Birch [14] (a PPL that compiles into C++), and Stan [15] (a platform for statistical modelling and computation). Note that this list is far from complete, and there exist many more experimental PPLs.

In this paper, WebPPL and Birch have been used for the reason of simplicity and efficiency, respectively. Some of the authors of this paper are currently developing a new domain-specific probabilistic programming language on top of the Miking [16] platform. This language, called TreePPL [c], is designed specifically for the domain of statistical phylogenetics.

In the rest of this section, we describe the key concepts of probabilistic programming. The examples are given in WebPPL, but could easily be translated into any of the other universal PPLs. The WebPPL code can be run in a web browser through the WebPPL project web page [d]. For a more comprehensive introduction to probabilistic programming, see for example the introductory text by van de Meent et al. [17].

#### 1.2 Sampling

The first key construct in probabilistic programs is *sample*, meaning that a value is drawn from a given probability distribution. Consider the following WebPPL code:

```
sample(Bernoulli({p: 0.5}))
```

The program models a simple coin flip scenario, where we sample from the Bernoulli distribution with probability 0.5, that is, a fair coin. When executed, the program returns either `true` or `false`, with probabilities corresponding to the sampled distribution.

Suppose we instead introduce another random variable $x$ that models the probability of getting heads on the toss of the coin. Mathematically, such a model can be defined as follows:

$$x \sim \text{Beta}(\alpha, \beta)$$
$$y \sim \text{Bernoulli}(x)$$

where the beta distribution is used as a prior probability distribution for the value of $x$. The same model can be written as a PPL program (assuming we set $\alpha = \beta = 2$)

```
var x = sample(Beta({a: 2, b: 2}))
var y = sample(Bernoulli({p: x}))
```

Note that the `sample` construct is conceptually used to denote random variables, in this case the two variables `x` and `y`. If we run the program many times and plot the values of `x`, we get an approximation of the probability density function (PDF) for $x$ in our mathematical model, that is, an approximation of the $\text{Beta}(2, 2)$ distribution. Because the expected value of $x$ is 0.5, `y` in the program still models an unbiased coin.

#### 1.3 Conditioning and observations

Probabilistic programs are based on Bayesian statistics, and typically are intended to compute the posterior distribution, given a prior distribution and some observations. In the coin flip example, suppose we observe heads (encoded as `true`) after flipping the coin once. We want to infer $p(x|y)$, the posterior distribution of $x$, conditioned on the new observation $y =$ `true`. As in the previous example, we assume that the prior distribution of $x$ is $\text{Beta}(2, 2)$. This model can be defined as follows.

```
var coinFlip = function() {
    var x = sample(Beta({a: 2, b: 2}))
    observe(Bernoulli({p: x}), true)
    return x
}
```

[c] https://treeppl.org/
[d] http://webppl.org

Note how the second `sample` construct is replaced with an `observe` construct. The program returns `x`, which is a (weighted) sample from the posterior distribution of $x$, computed by updating the prior distribution of $x$ (defined explicitly in the model) with the observation that the coin flip resulted in `true` (conditioning on the observation). Supplementary Figure 1A shows the prior distribution of the bias $x$, which corresponds to the Beta(2, 2) distribution. Note how the posterior distribution in Supplementary Figure 1B has moved closer to 1 (bias towards heads), compared to the prior distribution.

The `observe` statement is a way of weighting a sample according to some distribution. It is basically equivalent to sampling from a distribution, followed by conditioning, using the `condition` statement covered in the main text[e]. In WebPPL, there is also a construct called `factor`, which performs explicit weighting (also called scoring) of samples. A score or weight for a specific value can be computed using a distribution's PDF. In WebPPL, there is a `score` method that gives back the score of a value for a specific distribution. Thus, the `observe` statement in the previous example could be replaced with the following equivalent encoding using `factor`:

```
factor(Bernoulli({p: x}).score(true))
```

The `factor` construct is often used explicitly in the phylogenetic models described in this paper, especially in WebPPL.

#### 1.4 Recursive models and stochastic branching

In the previous examples, the models were very simple. However, the power of universal probabilistic programming is that a model can be *any* program, which may include variable declarations and standard control flow operators. Consider the following program that includes an `if` statement:

```
var mixture = function(){
  if (sample(Bernoulli({p: 0.7}))) {
    return sample(Gaussian({mu: -2, sigma: 1}))
  } else {
    return sample(Gaussian({mu: 3, sigma: 1}))
  }
}
```

The model illustrates the use of *stochastic branching*, meaning that the paths taken in a program depend on the outcome of sampling. In the example, the guard of the `if` statement samples from the Bernoulli distribution. Depending on whether the `true` or `false` branch is taken, sampling of the resulting value is done with different Gaussian distributions (different $\mu$ values). The plot of the model is shown in Supplementary Figure 1C. As can be seen in the figure, the `true` branch has larger weight, because of the probability of 0.7 of it being chosen.

Stochastic branching can be combined with recursion: this is a key building block for phylogenetic models. Consider the following model, which describes a model of the geometric distribution:

```
var geometric = function(p) {
    if (sample(Bernoulli({p: p}))) {
        return 1
    } else {
        return geometric(p) + 1
    }
}
```

Note that there is no requirement of a deterministic termination of the recursion: the termination of the recursion depends on the stochastic branch, which each time depends on a different latent random variable. Supplementary Figure 1D shows the plot of `geometric(0.6)`. The simple recursion above generates a linear sequence of random length. We use similar recursions in our scripts to model the processes that generate bifurcating phylogenetic trees. We do that by including two recursive calls within the same function, one for each descendant of a speciation event.

#### 1.5 Inference

The focus of this tutorial text has so far been on the model (the probabilistic program), and not on the inference algorithms. As discussed previously, in probabilistic programming, the choice of inference algorithm is intentionally separated from the model. For instance, using the `Infer` method of WebPPL, a user can apply the Sequential Monte Carlo (SMC) method to perform the inference of the coin example

```
Infer({model: coinFlip, method: 'SMC', particles: 20000})
```

or, alternatively, a Markov chain Monte Carlo (MCMC) method can be used:

```
Infer({model: coinFlip, method: 'MCMC', samples: 20000, burn: 5000})
```

The user also needs to specify the granularity of the approximation, using the number of particles or samples for SMC or MCMC, respectively.

In general, a key strength of the probabilistic programming paradigm is its expressive power, which is clearly shown in this paper within the domain of phylogenetics. One of the main research challenges within the PPL community is how to develop

[e]According to the WebPPL documentation, for efficiency, the `observe` statement should be used instead of the combination of sampling and conditioning, especially for continuous distributions.

inference algorithms and compilers that scale to very large and complex models. Although this is an active area of research, our study hopefully demonstrates that already state-of-the-art PPL systems make it possible to perform effective inference on non-trivial phylogenetic models.

### 2 Tools for phylogenetic probabilistic programming

The paper is accompanied by a code repository containing all the sources, tools and data used for the study, including documentation. Specifically, the resources in the repository are designed to facilitate the use of two existing probabilistic programming languages—WebPPL and Birch—for phylogenetic inference. The code repository is available at:

`https://github.com/phyppl/probabilistic-programming`

The reader is referred to the `README.md` file in the repository, in which we describe how to install the tools and how to use them to rerun our analyses or to experiment with probabilistic programming for phylogenetic problems.

#### 2.1 WebPPL for statistical phylogenetics

WebPPL is a universal probabilistic programming language based on JavaScript. We have written two packages, `phywppl` and `phyjs` that enable the reader to run phylogenetic simulations in WebPPL. The verification of the WebPPL programs relies on an auxiliary R package, `rppl`. Please refer to the aforementioned online documentation for further explanation.

#### 2.2 Birch for statistical phylogenetics

Birch is a universal probabilistic programming language compiling into C++. The models presented in this paper are run like regular Birch packages. Refer to the aforementioned online documentation for further explanation.

#### 2.3 Reading in phylogenetic trees

The WebPPL and Birch scripts we provide either simulate the diversification process along an observed reconstructed tree or computes the likelihood using analytical equations for such a tree. To facilitate the import of the observed tree data, we use a new JSON format for phylogenetic trees named PhyJSON [18]. Supported by the resources we provide in the repository, both WebPPL and Birch have mechanisms for reading in phylogenetic trees stored in the PhyJSON format. We also provide a stand-alone tool, `nexus2phyjson`, which can be used to convert trees in Nexus tree files to PhyJSON format [18].

For convenience, we include several phylogenetic trees in the `phyjs` package for purposes such as testing and verification (Table 1). An up-to-date account of the included test trees is provided in the `webppl/phyjs/README.md` file.

### 3 Diversification models

#### 3.1 Basic notation and terminology

All of the diversification models considered in this study can be generically described as follows (see Table 2 for a summary of the notation). The process starts at some time $t_0 > 0$ in the past, where $t = 0$ represents the present time. Evolutionary lineages split (speciate) at a per-lineage rate $\lambda$ and go extinct at per-lineage rate $\mu$. The rates $\lambda$ and $\mu$ are either constant, time-dependent or lineage-dependent, depending on the specific model. Each speciation event produces two lineages that further evolve independently of each other. The process is stopped when reaching the present ($t = 0$), at which point lineages still surviving are sampled (included in the observed tree) with probability $\rho \leq 1$.

The diversification process generates both trees with lineages that survive until the present, and trees that go completely extinct. Many surviving trees include side branches or whole subtrees that went extinct along the way. If the extinct parts and the branches leading to unsampled taxa are pruned away, we get what is called the "reconstructed tree". In other words, the reconstructed tree is the subtree spanned by only those surviving lineages that have been sampled.

In diversification analyses, the focus is typically on reconstructed trees. For simplicity, we assume here that the reconstructed tree is known without error, but we note that it is straightforward to extend our probabilistic programming approach to accommodate

**Table 1** Example trees provided in the `phyjs` package.

| Tree | Leaves | Age (Ma) | Description | Reference |
|---|---|---|---|---|
| `phyjs.bisse_32` | 32 | 13.0 | Example tree from Mesquite software | [19] |
| `phyjs.cetacean_87` | 87 | 35.9 | Cetacean tree from BAMMtools package | [20] |
| `phyjs.primates_233` | 233 | 65.1 | Primate tree from Diversitree package | [21] |

**Table 2** Summary of notation.

| Symbol | Interpretation |
|---|---|
| $\lambda$ | speciation (birth) rate |
| $\mu$ | extinction (death) rate |
| $\epsilon$ | turnover, $\mu/\lambda$ |
| $\rho$ | probability of sampling a leaf |
| $\lambda(t)$ | function characterizing time dependence of $\lambda$ in some models |
| $\lambda_0$ | initial $\lambda$, when $\lambda$ varies over time |
| $z$ | rate of exponential increase or decrease in $\lambda$ |
| $\lambda^i$ | speciation rate of process or branch $i$ |
| $\lambda^i(t)$ | time-dependent speciation rate of process $i$ |
| $\mu^i$ | extinction rate of process $i$ |
| $z^i$ | exponential rate of increase or decrease in $\lambda$ for process $i$ |
| $\alpha$ | long-term trend in $\lambda$ inheritance at speciation in ClaDS models |
| $\sigma$ | standard deviation (log scale) in $\lambda$ inheritance at speciation in ClaDS models |
| $\eta$ | rate of switching of diversification process in the BAMM and LSBDS models |
| $t_{\text{MRCA}}$ | age of most recent common ancestor |
| $\psi$ | reconstructed tree (extinct and unsampled side branches pruned away) |
| $n$ | number of leaves in the reconstructed tree |
| $V$ | the set of nodes (vertices) in the reconstructed tree |
| $a(i)$ | index of immediate ancestor of node $i$ |
| $l(i)$ | index of left descendant of node $i$ in oriented tree |
| $r(i)$ | index of right descendant of node $i$ in oriented tree |
| $c$ | number of cherries (terminal bifurcations) in a tree |
| $P(\cdot)$ | probability (density) |
| $L(\cdot)$ | likelihood |
| $S(t, \theta)$ | probability of process with parameters $\theta$ surviving from $t$ until the present |
| $Z$ | normalization constant of Bayes's theorem |

uncertainty about the tree by drawing the tree from an appropriate tree sample. Learning the parameters of the diversification process involves computing the likelihood of one or more reconstructed trees given different parameter values.

We denote a reconstructed tree $\psi = (V, t)$, where $V$ is a set of nodes (vertices) and $t$ is a corresponding vector of speciation ages. The tree has $n$ tips (terminal nodes or leaves) of degree one, $n - 1$ interior nodes of degree three, and the origin node of degree one. We index the nodes and their ages as follows:

- the origin has index 0;
- internal nodes have indices $\{1, 2, \ldots, n - 1\}$, ordered in decreasing age;
- tips have indices $\{n, n + 1, \ldots, 2n - 1\}$ (in any order)

The node $V_1$ corresponds to the first split between extant (surviving) lineages; it is referred to as *the most recent common ancestor* (MRCA) or *the root* of the reconstructed tree. The age of a node $i$ is $t_i$; leaves have age 0. A subtree with root at node $i$ and origin at time $t \geq t_i$ is denoted $\psi_i(t)$.

We will often find it convenient to distinguish between the two descendants of a node; without loss of generality, refer to them as the left and right descendant, respectively. A tree without leaf labels where nodes have been oriented in this way is an "oriented tree"[22].

We define three mapping functions for indices in an oriented tree:

- $a(i)$ is the index of the immediate ancestor of node $i$
- $l(i)$ is the index of the left descendant of node $i$
- $r(i)$ is the index of the right descendant of node $i$

#### 3.2 Conversions between tree spaces

In phylogenetics, we are interested in computing the likelihood of a labelled reconstructed tree, that is, a tree with leaf labels but with no distinction between the two descendants of a given ancestor. However, it is often convenient to derive the probability density of oriented trees without leaf labels first, and then convert it to a density on labelled trees without orientation[22]. The conversion factor is easy to find if we consider what happens if we start with a density on an oriented tree, then label it and finally remove the orientation. There are $n!$ unique ways of labelling an oriented tree, each with probability $1/n!$. When we remove

**Table 3** Overview of phylogenetic diversification models considered in the paper.

| Model | Full name | Reference |
|---|---|---|
| CRB | Constant rate birth model | Yule [24], Nee [25] |
| CRBD | Constant rate birth-death model | Feller [26] |
| TDB | Time-dependent birth model | Kendall [27] |
| TDBD | Time-dependent birth-death model | Kendall [27] |
| BAMM | Bayesian analysis of macro-evolutionary mixtures | Rabosky [28] |
| LSBDS | Lineage-specific birth-death shift model | Höhna et al. [29] |
| ClaDS[0-2] | Cladogenetic diversification rate shift models | Maliet et al. [30] |

the orientation, there are $2^{n-1}$ labelled oriented trees that produce the same labelled tree without orientation, where $n-1$ is the number of interior nodes in the tree. Thus, the conversion factor is $2^{n-1}/n!$.

For completeness, we derive the conversion factor with the operations in the reverse order, first dropping the orientation and then applying the labels. When labels are missing, there are $2^{n-1-c}$ unique oriented trees for each tree without orientation, where $c$ is the number of "cherries". A cherry is a pair of leaves that are each other's closest relatives[23]; without labels the descendants of a cherry are identical and there is only one unique way in which they can be oriented. Labelling a tree without orientation is similarly affected by cherries, so that there are $n!\,2^{-c}$ unique label assignments. The conversion factor is thus $2^{n-1-c}/(n!\,2^{-c}) = 2^{n-1}/n!$.

In the literature on advanced diversification models, it is common practice to derive the density on unlabelled oriented trees and ignore the conversion to a density on labelled unoriented trees; in fact, the omission of this factor is rarely acknowledged. This contrasts with the derivation of the analytical likelihood for simple models, such as CRBD, where the conversion factor is almost always accommodated. Previous work on diversification models has focused on a single model and a single tree; in such cases, ignoring the conversion factor is not a problem. However, here we compare diversification models using Bayes factors, so the normalization constant needs to be computed consistently for all models, that is, based on the same outcome space.

For convenience, our simulations of diversification processes assume unlabeled and oriented trees. This makes the scripts simpler, and it facilitates comparison to previous descriptions of these models. Our simulations are weighted with the appropriate conversion factor to generate the density for labelled and unoriented trees. Thus, the normalization constants (marginal likelihoods) we compute are directly comparable to the likelihoods computed using the standard analytical equations established for the simple diversification models, such as CRBD[22], for labelled and unoriented trees.

#### 3.3 Conditioning on the age of the MRCA

A process that starts at some time $t_0$ in the remote past will produce a reconstructed tree that has a *stalk*, i.e., a branch leading to the MRCA. However, we usually do not have any information about the length of this stalk. For this reason, and others, it is often more convenient in practice to condition the process on the first split in the reconstructed tree, $t_1$[22]. This can easily be done by noting that $t_1$ can be considered the time of origin for both the left and the right subtrees originating from the first split, and that both of these lineages survived until the present by the very definition of the concept of MRCA. Thus, the probability of the reconstructed tree, conditioned on the age of the MRCA, is obtained by multiplying together the probabilities of the left and the right subtrees and by conditioning on their joint survival. Given an oriented tree $\psi$, now without the "stalk" from the origin to the most recent common ancestor, the likelihood is thus given by

$$L(\psi \mid \theta, t_1) = \frac{P(\psi_{l(1)}(t_1)|\theta, t_1)P(\psi_{r(1)}(t_1)|\theta, t_1)}{(S(t_1, \theta))^2}, \tag{1}$$

where $\theta$ is the vector of parameters of the model, and $S(t, \theta)$ is the probability of the process surviving (producing at least one sampled descendant) after time $t$.

#### 3.4 Diversification models

In the paper, we consider nine different diversification models (Table 3).

The CRB(D) and TDB(D) models are simple diversification models, which assume that the process is the same for the entire tree, even though it can change over time. The other models (the advanced models) accommodate lineage-specific variation in diversification rates. The BAMM and LSBDS assume that the diversification process changes in a major way at certain points in time. In fact, the process is completely reset. Thus, LSBDS and BAMM can be described as models of punctuated change in diversification. The ClaDS models instead assume gradual, heritable changes in speciation and extinction rates. Specifically, this is modeled as small step-wise changes associated with speciation events (also called cladogenetic events).

We provide a tabular summary of the parameters of each model (Table 4) to facilitate comparison across them. We describe each model in detail below.

**Table 4** Summary of diversification model parameters.

| Model | Parameters | Notes |
|---|---|---|
| CRB | $\lambda$ | $\lambda$ is speciation rate |
| CRBD | $\lambda, \epsilon$ | $\epsilon = \mu/\lambda$ is turnover rate |
| TDB | $\lambda(t), z$ | $z$ is exponential time-dependence parameter |
| TDBD | $\lambda(t), \epsilon, z$ | |
| LSBDS | $\eta, \{(\lambda^i, \mu^i)\}$ | $\eta$ is change rate, $i$ is index of process |
| BAMM | $\eta, \{(\lambda^i, \mu^i, z^i)\}$ | $z^i$ is time-dependence parameter of process $i$ |
| ClaDS0 | $\alpha, \sigma, \{\lambda^i\}$ | $\alpha$ is trend parameter, $\sigma$ is noise parameter in $\lambda$ inheritance at speciation; $i$ is branch index in complete tree |
| ClaDS1 | $\alpha, \sigma, \mu, \{\lambda^i\}$ | $\mu$ is extinction rate |
| ClaDS2 | $\alpha, \sigma, \epsilon, \{\lambda^i\}$ | $\epsilon$ is turnover rate |

###### 3.4.1 Constant rate birth-death models (CRBD and CRB)

The constant rate birth-death (CRBD) model is the simplest model considered here. Evolutionary lineages split at a constant per-lineage birth rate $\lambda$ and go extinct at a constant per-lineage death rate $\mu$. The parameter vector for this model is thus $\theta = (\lambda, \mu, \rho)$. As a special case, we consider the constant rate birth (CRB) model, also known as the Yule model, with $\mu = 0$[25].

The probability that a CRBD process starting at time $t$ survives until the present and is sampled is known analytically[22]; it is

$$S(t, \lambda, \mu) = \frac{r}{\lambda - (\lambda - r/\rho)\, e^{-rt}}, \tag{2}$$

where $r = \lambda - \mu$ is known as the "net diversification rate". The likelihood of a reconstructed tree conditioned on the time of the MRCA is also known analytically; it is given by:

$$L(\psi|\theta, t_1) = \frac{2^{n-1}}{n!}\, \lambda^{n-2}\, \rho^n\, \frac{\widehat{g}(t_1)^2 \prod_{i=2}^{n-1} \widehat{g}(t_i)}{\widehat{g}(0)^n\, S(t_1)^2}, \tag{3}$$

where

$$\widehat{g}(t) = \frac{e^{-rt}}{(\lambda - (\lambda - r/\rho)\, e^{-rt})^2}. \tag{4}$$

Even though the likelihood is known analytically, there are no conjugate priors for $\lambda$ and $\mu$ that would yield an analytical posterior. Thus, we end up with an intractable integral if we want to learn these parameters from one or more observed trees.

###### 3.4.2 Time-dependent birth-death models (TDBD and TDB)

In the time-dependent birth-death model (TDBD), the speciation rate is assumed to change continuously through time. More specifically, we consider the following time-dependence:

$$\lambda(t) = \lambda_0\, e^{z(t_1 - t)}. \tag{5}$$

Thus, $\lambda_0$ is the speciation rate prevailing at the time of the MRCA. Furthermore, if $z < 0$ (resp. $z > 0$), the speciation rate decreases (resp. increases) exponentially when going toward the present. An exponentially decreasing speciation rate can be seen as an approximate model for diversity dependence. The parameter vector for this model is $\theta = (\lambda_0, \mu_0, x, \rho)$.

The likelihood appears to be intractable for the present model with exponentially varying speciation rate and constant extinction rate. On the other hand, a simple solution is available for the slightly different model examined here, in which $\lambda$ and $\mu$ are both exponentially decreasing or increasing at the same rate $z$ (and thus the turnover rate $\lambda/\mu$ is constant):

$$\lambda(t) = \lambda_0\, e^{z(t_1 - t)},$$
$$\mu(t) = \mu_0\, e^{z(t_1 - t)}.$$

Under this model, the probability that a lineage starting at time $t$ survives until the present and is sampled is now (for a general method of deriving the likelihood for time-dependent birth-death models, see Morlon et al.[31]):

$$S(t, \lambda_0, \mu_0, z) = \frac{r_0}{\lambda_0 - (\lambda_0 - r_0/\rho)\, e^{-(r_0/z)(1 - e^{-zt})}}, \tag{6}$$

where $r_0 = \lambda_0 - \mu_0$. The likelihood of a reconstructed tree conditioned on the time of the MRCA has the same general form as for the CRBD:

$$L(\psi|\theta, t_1) = \frac{2^{n-1}}{n!}\, \rho^n\, \frac{\widehat{g}(t_1)^2 \prod_{i=2}^{n-1} \widehat{g}(t_i)\, \lambda(t_i)}{\widehat{g}(0)^n\, S(t_1)^2} \tag{7}$$

with $S$ such as just given and:

$$\widehat{g}(t) = \frac{e^{-(r_0/z)(1-e^{-zt})}}{\left(\lambda_0 - (\lambda_0 - r_0/\rho)\, e^{-(r_0/z)(1-e^{-zt})}\right)^2}. \tag{8}$$

As a special case, we consider the time-dependent birth (TDB) model, which is equivalent to TDBD except that there is no extinction, that is, $\mu = 0$. Note that the TDBD model collapses to CRBD when $z = 0$. Similarly, TDB becomes equivalent to CRB when $z = 0$.

##### 3.4.3 Bayesian analysis of macroevolutionary mixtures (BAMM)

The BAMM model was proposed by Rabosky[28]. The original formulation of the change process is statistically incoherent[32] but it is straightforward to fix this, and we follow the slight reinterpretation of the model suggested by Moore et al.[32]. In this version, BAMM is an episodic, Poisson-modulated, birth-death process with exponentially decaying speciation rate. To describe the process, consider a generic lineage $e$ at time $t$. At this time point, the lineage is associated with rate parameters with index $e(t)$. Specifically, the lineage carries with it a triplet of rates $(\lambda^e(t), \mu^e(t), z^e(t))$. Then:

- at rate $\mu^e(t)$, the lineage goes extinct;
- at rate $\lambda^e(t)$, the lineage splits into two lineages (say $f$ and $g$), in which case the two daughter lineages inherit the current rates, i.e.

$$(\lambda^f(t), \mu^f(t), z^f(t)) = (\lambda^e(t), \mu^e(t), z^e(t)),$$
$$(\lambda^g(t), \mu^g(t), z^g(t)) = (\lambda^e(t), \mu^e(t), z^e(t));$$

- $\lambda^e(t)$ increases or decays exponentially at rate $z$;
- at rate $\eta$, the triplet of rate parameters is redrawn from a pre-specified trivariate distribution $\Phi$, i.e.

$$(\lambda^e(t_-), \mu^e(t_-), z^e(t_-)) \sim \Phi \tag{9}$$

The process starts with a single lineage $a$ at some time $t_0$, with rate parameters $(\lambda^a(t_0), \mu^a(t_0), z^a(t_0)) \sim \Phi$. Specifically, we start the process at the time immediately before the first split in the tree (at the MRCA), and we assume that the process index at this point is $a(t_{\text{MRCA}}) = o$ (o for origin). The prior distributions used in this paper for $\Phi$ were chosen to harmonize with the priors used for other models, as specified in the section on priors below. The process stops when reaching the present ($t = 0$) and the surviving lineages are sampled with probability $\rho$.

The likelihood under the BAMM model as defined here does not have an analytical solution, nor does it seem to be amenable to any known numerical techniques for solving the complex differential equations involved[32]. A recent paper[33] analyzes a variant of the BAMM model, which differs from the variant considered here in that the rate of diversification model shifts is considered to be constant for a clade, regardless of the number of lineages the clade contains. This is a somewhat unusual type of model, as lineages are usually considered to evolve independently and not as parts of a larger clade. Nevertheless, under this assumption they succeed in deriving an analytical expression of the likelihood for the case of a single shift. They then extend this to multiple shifts, keeping the likelihood computations manageable by assuming that a clade that has shifted diversification rates once cannot shift again.

Describing the BAMM model as defined here (or in the clade-spanning variant) using a PPL is straightforward. Effective inference can then be performed using generic techniques available for PPLs, such as SMC or PMCMC (particle Markov chain Monte Carlo), as we show in the current paper. No artificial model constraints need to be introduced.

##### 3.4.4 LSBDS

The recently proposed LSBDS model[29] can be seen as a specialized version of the BAMM model, in which $z = 0$ at all times and for all lineages. In other words, there is no exponential decay of speciation rates; the speciation rate remains constant between rate shift events. As a result, $\Phi$ is now a bivariate distribution.

Under these conditions, it becomes possible to compute the likelihood of a reconstructed tree by approximating $\Phi$ as a product of two discrete distributions, with a finite (and small) number of possible values for $\lambda$ and $\mu$, and then relying on standard numerical techniques for solving the differential equations involved[34,29]. This is similar to the discretization approach frequently used in phylogenetics in order to efficiently approximate the likelihood when rates vary across sites according to a continuous gamma distribution[35].

Using this approach, the backward-in-time recursion for the extinction probability and for the conditional likelihood, which are both conditioned on the current value of $\lambda$ and $\mu$ for the lineage under consideration, entails a set of $LM$ coupled master equations, where $L$ and $M$ are the number of bins used for the $\lambda$ and $\mu$ distributions, respectively. In practice, this imposes a rather strict constraint on the number of discretization bins that can be used, as the computational complexity otherwise becomes unmanageable. We note that the empirical examples discussed in the LSBDS paper all use a fixed value for $\mu$ across the tree, thus effectively setting $L = 1$. The SMC techniques we use in the current paper do not suffer from such limitations, as they rely on sampling values from the $\lambda$ and $\mu$ distributions.

Another model that also eliminates the $z$ variable of the BAMM model was presented recently[36]. This model is based on restricting the number of possible diversification models to a finite number $k$ of different diversification model categories. An MCMC algorithm is then used to sample over different histories of shifts among these categories over the tree. To simplify the computation of likelihoods, the authors further assume that no shifts among model categories occur on extinct side branches. In our PPL and SMC context, we do not gain much by restricting the number of diversification rate models, and there is no need to simplify the treatment of extinct side branches. Thus, we do not consider this model further here.

###### 3.4.5 The cladogenetic diversification rate shift models (ClaDS)

The ClaDS models[30] assume that the speciation rate changes by a small random amount at each speciation event. The extinction rate is assumed to be either equal to 0 (ClaDS0), constant but positive (ClaDS1), or proportional to the speciation rate, such that the turnover rate ($\epsilon = \mu/\lambda$) is constant (ClaDS2). Thus, in all cases, $\mu = \mu(\lambda)$ can be seen as a (possibly constant, for ClaDS0 and ClaDS1) deterministic function of $\lambda$.

Consider a generic lineage $e$ at time $t$. This lineage carries with it a rate $\lambda^e$. Then:

- at rate $\mu(\lambda^e)$, the lineage goes extinct;
- at rate $\lambda^e$, the lineage splits into two lineages (say $f$ and $g$), in which case the two daughter lineages draw their respective speciation rates, $\lambda^f$ and $\lambda^g$ as follows:

$$\log \lambda^f \sim \mathcal{N}(\log(\alpha\lambda^e), \sigma^2).$$
$$\log \lambda^g \sim \mathcal{N}(\log(\alpha\lambda^e), \sigma^2).$$

The process starts with a single lineage $o$ at some time $t_0$, with speciation rate $\lambda^o$.

The $\alpha$ parameter introduces a deterministic long-term trend in the otherwise random variation of $\lambda$ through time, across the many speciation events typically occurring over the complete phylogeny. When $\alpha < 1$ (resp. $\alpha > 1$), the logarithm of the speciation rate decreases (resp. increases) on average at a speciation event.

The original ClaDS paper[30] focuses on the mean diversification rate multiplier $m = \alpha \exp(\sigma^2/2)$ rather than on $\alpha$. However, we prefer the $\alpha$ parameterization because it allows us to find a convenient conjugate prior that supports delayed sampling and thus makes SMC inference more efficient (see below).

There is some superficial resemblance between $log\alpha$ and the $z$ parameter in the TDBD, TDB and BAMM models, also suggesting that this would be a natural parameterization. However, the dynamics of the ClaDS models is quite different from the TDBD, TDB and BAMM models. This makes the interpretation of both the $\alpha$ and $m$ parameters more complicated than what is immediately obvious. When there is noise in the ClaDS models ($\sigma > 0$), there will be variation across lineages in diversification rate. This will result in lineage selection, ensuring that the average diversification rate across lineages goes up more than suggested by the values of $m$ or $\alpha$[37]. Even within lineages, there will be slight deviations from the behavior that might be expected from the $m$ (or $\alpha$) values. For instance, when there is noise, setting $m = 1$ will result in the expected diversification rate at the end of some specified time period being lower than the starting rate. This is because the process is more likely to reach the end of the time period without any further change if the diversification rate goes down than if it goes up.

We note that it would be straightforward to estimate posterior distributions on $m$ using our scripts even though they use the $\alpha$ parameterization. This can be achieved either by adding a line that computes $m$ from $\alpha$ and $\sigma$ and then returns it to the model scripts before running the analysis, or by converting the sampled $\alpha$ and $\sigma$ values to $m$ values after the analysis has completed.

The likelihood under the ClaDS1 and ClaDS2 models is not analytically available, but it can be numerically evaluated[30]. The evaluation method originally developed for the ClaDS models involved various numerical approximation techniques, including discretization of time and rate space, and expands over thousands of lines of code in the RPANDA R package. Recently, considerably more efficient inference techniques based on data augmentation have been developed for these models[37].

#### 4 Prior probability distributions

To facilitate the interpretation of the Bayes factor tests, we standardized prior probability distributions across diversification models as much as possible in our analyses. Before going into details, it may be helpful to explicitly declare the parameterizations we assume for the statistical distributions used. Thus, for the exponential distribution, we assume the rate parameterization, for the inverse gamma distribution we use the shape-scale parameterization, and finally, for the normal (Gaussian) distribution, the parameters are the mean and the variance of the distribution.

Across all models, we used an Exponential(1) prior for the speciation rate, and a Uniform(0, 1) prior for the turnover rate, both common priors in the diversification model literature. The specific implementations are listed for each model below. For the $\sigma$ parameter of the ClaDS models, Maliet et al.[30] used a prior with most probability mass close to 0 ($\sigma \sim \text{InvGamma}(1, \log(1.1))$). Upon examination of the empirical results published in the same paper (shown in their Figure 4a), we concluded that this choice is overly conservative. That is, the prior puts so much probability on low values of $\sigma$ that the posterior may underestimate the extent of lineage-specific variation in diversification rates. We also note that it is more natural to consider an inverse gamma prior for the variance rather than the standard deviation of the normal distribution, since this is a conjugate prior for the normal distribution. Therefore, we used a $\sigma^2 \sim \text{InvGamma}(1, 0.2)$ prior in our analyses.

The original ClaDS paper[30] used an improper prior for the $\alpha$ parameter. This is not suitable for our purposes, as we need to simulate from the prior in SMC. We instead assumed $\log \alpha \sim \mathcal{N}(0, \sigma^2)$. By making the variance of the $\log \alpha$ prior dependent on $\sigma^2$, we establish a conjugate normal-inverse-gamma prior. This results in a joint prior on $(\log \alpha, \sigma^2)$ that has its mode for $\alpha$ at 0, at which point there is neither acceleration nor deceleration of speciation rates. The posterior distribution of $\alpha$ values reported earlier for the bird trees under the Clads2 model[30] is also well covered by this joint prior. For $z$, we used the prior proposed in the original BAMM paper[28], namely $z \sim \mathcal{N}(0, 0.05^2)$.

Finally, for the LSBDS and BAMM models, we wanted a prior on $\eta$ that was scaled to time. In the LSBDS and BAMM papers[29,28,38], it has been common to instead specify a prior scaled to the total length of the tree. This allows one, for instance, to specify a prior with an expectation of one change in the diversification process over the reconstructed tree. However, if the changes we observe in diversification rates are the result of some evolutionary process, then it would seem more reasonable to assume that the expected number of changes is a function of evolutionary time rather than of an arbitrarily circumscribed reconstructed tree. To obtain this effect, while still maintaining some scaling to tree size, we chose a prior with one expected change in diversification rates over the time period from the most recent common ancestor to the present. For a small reconstructed tree, this would correspond to an expectation of slightly more than one change over the tree, while the expectation could be more than a few changes in a big tree. Specifically, we assumed $\eta \sim \text{Exponential}(t_{\text{MRCA}})$, where $t_{\text{MRCA}}$ is the age of the first split in the tree.

For completeness, all prior probability distributions are listed below for each of the examined models (see also Supplementary Figure 2).

##### 4.1 CRB

The CRB model has only one parameter, $\lambda$, for which we use the standard prior:

$$\lambda \sim \text{Exponential}(1).$$

##### 4.2 CRBD

The CRBD model has two parameters, $\lambda$ and $\mu$. For $\lambda$ we use the standard prior, and for $\mu$ the indirect prior induced by assuming a uniform prior on the turnover rate $\epsilon = \mu/\lambda$.

$$\begin{aligned} \lambda &\sim \text{Exponential}(1), \\ \epsilon &\sim \text{Uniform}(0, 1). \end{aligned}$$

##### 4.3 TDB

For the TDB model, we applied the standard priors as follows:

$$\begin{aligned} \lambda_0 &\sim \text{Exponential}(1), \\ z &\sim \mathcal{N}(0, 0.05^2), \end{aligned}$$

where $\lambda_0$ is the initial speciation rate, and $z$ is the time dependence parameter in $\lambda(t) = \lambda_0 e^{zt}$. In other words, the standard $\lambda$ prior applies to the initial speciation rate in this model.

##### 4.4 TDBD

The TDB priors are extended to the TDBD case as follows:

$$\begin{aligned} \lambda_0 &\sim \text{Exponential}(1), \\ z &\sim \mathcal{N}(0, 0.05^2), \\ \epsilon &\sim \text{Uniform}(0, 1), \end{aligned}$$

where $\lambda_0$ is the initial speciation rate, $z$ is the time dependence parameter in $\lambda(t) = \lambda_0 e^{zt}$, and $\epsilon$ is the turnover rate. Note that in our implementation we have kept the turnover rate constant (rather than the extinction rate), i.e., $\mu(t) = \epsilon\lambda(t)$.

##### 4.5 ClaDS0

For the ClaDS0 model, we applied the standard $\lambda$ prior to the initial speciation rate, in line with the TDB(D) models. The $\alpha$ and $\sigma$ priors are justified above. Specifically, the ClaDS0 priors we used are:

$$\begin{aligned} \lambda_0 &\sim \text{Exponential}(1), \\ \sigma^2 &\sim \text{InvGamma}(1, 0.2), \\ \log \alpha &\sim \mathcal{N}(0, \sigma^2), \end{aligned}$$

where $\lambda_0$ is the initial speciation rate, $\sigma^2$ represents the variance in the inherited speciation rate and $\alpha$ is the *speciation trend parameter*.

##### 4.6 ClaDS1

The prior distributions related to the speciation rates are the same as for ClaDS0. In addition, we assume

$$\epsilon \sim \text{Uniform}(0, 1),$$

where $\epsilon$ is the initial turnover rate. The extinction rate, $\mu = \epsilon\lambda_0$, remains constant in the whole tree.

##### 4.7 ClaDS2

The prior distributions related to the speciation rates are the same as for ClaDS0. In addition, we assume

$$\epsilon \sim \text{Uniform}(0, 1),$$

where $\epsilon$ is the turnover rate. Unlike ClaDS1, the extinction rate changes at each speciation such that the turnover remains constant over the whole tree.

##### 4.8 LSBDS

For the LSBDS model, we define the joint prior $\Phi$, such that the all $\lambda^i$ values, including the speciation rate for the MRCA ($\lambda^o$), are drawn from the standard $\lambda$ prior used for other models, and such that the $\mu^i$ values, including the value of the MRCA, are drawn independently from the distribution induced by drawing $\epsilon$ from the standard uniform distribution used for other models. Specifically,

$$\begin{aligned}
\eta &\sim \text{Exponential}(t_{\text{MRCA}}),\\
\lambda^i &\sim \text{Exponential}(1),\\
\epsilon^i &\sim \text{Uniform}(0, 1),
\end{aligned}$$

where $\eta$ is the rate of shifts in diversification processes, $t_{\text{MRCA}}$ is the time (age) of the most recent common ancestor, and $\lambda^i$ and $\epsilon^i$ are the speciation and the turnover rates of the $i$-th diversification process.

##### BAMM

The prior distributions for BAMM are the same as for LSBDS, with addition of

$$z^i \sim \mathcal{N}(0, 0.05^2),$$

where $z^i$ is the time dependence parameter for the speciation rate of the $i$-th diversification process. This is the same prior distribution used for the $z$ parameter of the TDB and TDBD models.

#### 5 PPL model descriptions

In this section, we describe the PPL model scripts we used in the paper. We focus on WebPPL, as we think these model scripts are the most accessible to biologists. We start by describing model scripts that make use of the analytical likelihood equations. We then present the complete description of the explicit simulation script for the CRBD model that is partly covered in the main paper. Finally, we provide a brief overview of the simulation scripts for the remaining models. We end the section with a brief discussion of how the Birch scripts are similar to and how they differ from the WebPPL scripts. For full details, we refer the interested reader to the code repository accompanying the paper.

##### 5.1 Scripts based on analytical likelihoods

As mentioned in Section 3, the likelihood of a reconstructed tree conditioned on the age of the MRCA and the parameters of the diversification process is known analytically for the simple diversification models (CRB, CRBD, TDB, TDBD). We can take advantage of this in probabilistic programs, facilitating efficient inference of model parameters, by scoring simulations according to the analytical likelihood. To simplify the implementation of such scripts, we provide the analytical likelihoods as deterministic functions in the `phyjs` library. Four functions are available. The function `exactCRBDLikelihoodComplete (tree, lambda, mu)` computes the likelihood of a reconstructed tree under the CRBD model for specific values of $\lambda$ and $\mu$, assuming complete sampling of the leaves (tips) in the tree, $\rho = 1$. The function `exactCRBDLikelihoodRandom (tree, lambda, mu, rho)` computes the same likelihood when the leaves are randomly sampled with probability $\rho < 1$. Finally, the functions `exactTDBDLikelihoodComplete (tree, lambda, mu, z)` and `exactTDBDLikelihoodRandom (tree, lambda, mu, z, rho)` compute the corresponding probabilities for the TDBD model. By setting `mu = 0`, the functions can be used to compute the likelihoods for the CRB and TDB models.

The following listing shows how to infer the posterior distribution of $\lambda$ and $\epsilon$ for the CRBD model using the analytical likelihood and the MCMC inference method:

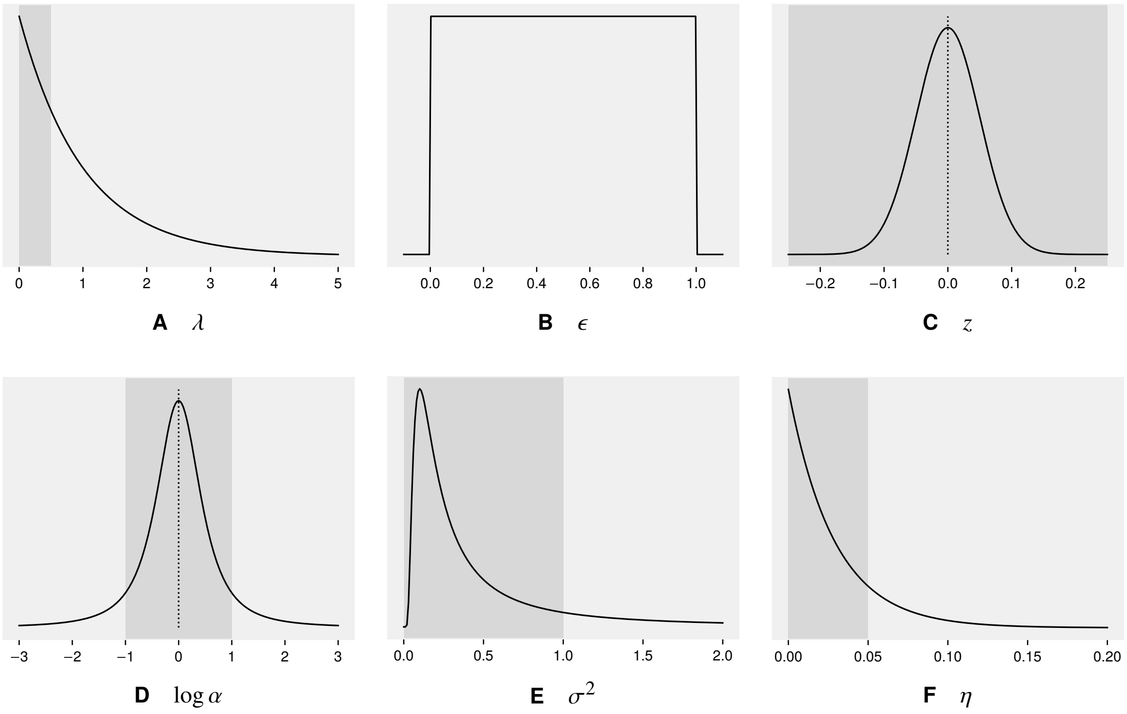


**Fig. 2** Prior distributions of model parameters. The shaded regions correspond to the region of parameter space illustrated in the posterior plots for the empirical analyses (Supplementary Figures 13–22). See also Supplementary Figure 23.

**Algorithm 1** CRBD model with analytical likelihood.

```
var tree = phyjs.bisse_32

var model = function() {
    var lambda = exponential({ a: 1 })
    var epsilon = uniform({ a:0.0, b: 1.0 })
    var mu = epsilon*lambda

    factor( exactCRBDLikelihoodComplete(tree, lambda, mu) )

    return [lambda, epsilon]
}

var dist = Infer({method: 'MCMC', samples: 100, lag: 10, burn: 1000, model: model})

dist
```

In the script, we first select one of the provided trees in the `phyjs` package. The model is then set up by specifying the priors on the model parameters, and computing the value of the extinction rate $\mu$, encoded as the variable `mu`. The simulation then scores the simulation according to the analytical likelihood of the sampled parameter values using the `factor` construct. In the final line of the `model` function, the values of the model parameters are returned.

For inferring the posterior distribution induced by the model function, the MCMC method is a good choice. For explanation of the inference settings, see the WebPPL documentation of the MCMC method[f]. The last line ensures that the estimated joint posterior distribution, encoded as `dist`, is printed.

The script can be run using the commands we provide in the code repository accompanying this paper[g], as explained in the documentation provided there. In the directory `webppl/phywppl/examples/` in the repository, we provide analytical scripts of this kind for the CRB, CRBD, TDB and TDBD models.

#### 5.2 Basic script for CRBD

Here, we give a complete WebPPL implementation of the CRBD model. The program describing the model is divided into two files to facilitate reuse of the code. The simulation part is specified in one file, and the analysis part in another. The simulation file contains code that simulates the CRBD process along a given tree for specified values of the model parameters. This file can be reused unaltered regardless of the particular analysis one wants to perform. The analysis file contains the specification of the priors, the data, and the inference method. This file needs to change from one analysis to another.

[f] `http://docs.webppl.org/en/master/inference/methods.html#mcmc`
[g] `https://github.com/phyppl/probabilistic-programming`

The analysis file (Algorithm 2) is structured in the same way as the script using the analytical likelihood for CRBD. However, instead of calling a function to compute the analytical likelihood, we call the simulation function for the CRBD model. This function weights the simulation appropriately for the given parameter values, conditioned on the observed reconstructed tree. The model function returns the model parameters, as before. However, instead of inferring the posterior distribution on those parameters, we now use SMC and focus on the normalization constant (the marginal likelihood or model evidence). The normalization constant estimate is available in the `normalizationConstant` property of the distribution object returned by the `Infer` function when the method is SMC. Similar example scripts are available in the `webppl/phywppl/examples/` directory for all models studied in the paper.

**Algorithm 2** Analysis script for CRBD simulation.

```
var tree = phyjs.read_phyjson("bisse_32.phyjson")

var model = function() {
   var lambda = exponential({ a: 1 })
   var epsilon = uniform({ a:0.0, b:1.0 })
   var mu = epsilon*lambda

   simCRBDNaive( tree, lambda, mu)

   return [lambda, epsilon]
}

var dist = Infer({method: 'SMC', particles: 10000, model: model})

dist.normalizationConstant
```

Let us now turn to the simulation script (Algorithm 3). The script presented here is a naive PPL implementation of the CRBD model in that it does not use the analytical likelihood. Instead, it explicitly simulates the speciation and extinction process conditioned on the reconstructed tree. The script is also naive in the sense that it does not include any modifications to support aligned SMC inference, which is important for improving inference efficiency. The advanced inference techniques we used in the paper, including alignment, are discussed in Section 6. The script forms a basic template that can be used to express all diversification models analyzed in our paper. It should also be straightforward to extend the script to a range of new diversification models that have not been explored previously.

**Algorithm 3** A complete WebPPL script for simulating CRBD.

```
var goesExtinct = function( startTime, lambda, mu )
{
   var t = exponential( {a: lambda + mu} );

   var currentTime = startTime - t;

   if ( currentTime < 0 ) {
         return false
   }

   var speciation = flip( lambda/(lambda+mu) )
   if ( !speciation )
      return true;

   return( crbdGoesExtinct( currentTime, lambda, mu )
         && crbdGoesExtinct( currentTime, lambda, mu ) );
}

var simBranch = function( startTime, stopTime, lambda, mu )
{
   var t = exponential ( {a: lambda} );

   var currentTime = startTime - t;

   if ( currentTime <= stopTime )
      return 0.0;

   factor( Math.log( 2.0 ) );
   condition ( crbdGoesExtinct( currentTime, lambda, mu ) )

   return simBranch( currentTime, stopTime, lambda, mu )
}

var simTree = function( tree, parent, lambda, mu )
{
   factor( - mu * ( parent.age - tree.age ) );

```

```
38     simBranch( parent.age, tree.age, lambda, mu );
39 
40     if ( tree.type == 'node')
41     {
42         factor( Math.log( lambda ) );
43 
44         simTree( tree.left, tree, lambda, mu )
45         simTree( tree.right, tree, lambda, mu )
46 
47     }
48 }
49 
50 var simCRBDNaive = function( tree, lambda, mu )
51 {
52     var numLeaves = phyjs.countLeaves( tree )
53     var corrFactor = ( numLeaves - 1 ) * Math.log( 2.0 ) - phyjs.lnFactorial( numLeaves )
54     factor( corrFactor )
55 
56     simTree( tree.left, tree, lambda, mu )
57     simTree( tree.right, tree, lambda, mu )
58 }
```

The main function in the script is `simCRBDNaive`, defined at the end of the script. It takes three parameters: the model parameters `lambda` and `mu`, and the `tree` on which to condition the simulation (note the actual implementation order). For simplicity, the process is simulated along an oriented and unlabelled tree (see Section 3.2). This assumption allows us to ignore the probability factor associated with rotation and labeling of the reconstructed tree during the main part of the simulation. To ensure that the simulation nevertheless carries the right weight, it is first endowed with the appropriate rotation and labeling probability (see Section 3.2) using two utility functions in the `phyjs` library and the `factor` construct in WebPPL. This is important for computing the correct normalizing constant, but does not affect inference otherwise, since this probability factor is the same for all simulations. Note that, for numerical stability, the particle weights in WebPPL are stored as logarithms.

Next, the function `simTree` is called on both children of the root node (the MRCA), initiating the recursion over the observed tree. Note that `simCRBDNaive` does not return anything. It is called only for the side-effect of weighting the sampled `lambda` and `mu` values by conditioning the simulation on the observed tree.

The function `simTree` is similar in structure to `simCRBDNaive`: it computes various weights and, if we have not reached a leaf, continues the recursion. Here, we present a naive implementation of |simTree|, where the execution of the probabilistic program is reweighted as soon as the information becomes available via calls to |factor|. A bit later, when we discuss advanced inference techniques (Section 6), we will show a version of the script, which only reweights the particle after the hidden side-branches have been processed. In the naive version, the first step is to factor the probability of no extinction on the branch from the parent to the node (line 36). This corresponds to observing zero extinction events from a Poisson distribution parameterized by the extinction rate |mu| and the branch length |parent.age - tree.age|, i.e. the weight can be obtained by plugging in 0 in the probability density function (pdf) of the Poisson distribution. Next, we simulate the hidden side-branches (line 38), the function will re-weight the computation accordingly. Finally, if we are at a node, we observe an immediate speciation event, i.e. 0 from an exponential distribution with parameter |lambda|, which has the weight of |Math.log(lambda)|.

The `simBranch` function recursively simulates speciation events along the branch. If there is a speciation event, the side branch it generates must go extinct, as it is not present in the observed reconstructed tree. We call such a speciation event a "hidden" speciation because it is not visible in the observed tree. To condition the simulation on the extinction of the side branch resulting from a hidden speciation, we require the call to the recursive simulation function `goesExtinct` to return true. The `goesExtinct` function is described in the main paper; it is defined at the top of the script presented here. It simulates an outcome of the birth-death process for given `lambda` and `mu` values, starting at a given time in the past and counting downwards until the present (time 0). If all lineages go extinct before reaching the present, the function returns `true`, otherwise it returns `false`. In connection with the call to `goesExtinct`, the `simBranch` function also needs to take a rotational factor into account. This arises because there are two indistinguishable simulations that correctly account for the tree we condition on: one in which the right descendant of the hidden speciation event goes extinct and the left descendant gives rise to the observed continuation of the lineage, and one in which the left descendant goes extinct and the right descendant gives rise to the observed continuation of the lineage. Thus, the correct probability score for the simulation is twice what would have resulted from a single call to `goesExtint`, and we therefore need to add $\log 2$ to the weight (recall that probability factors are represented on the log scale in WebPPL) before continuing the recursion.

The analysis and simulation scripts described above are simplified versions of the example script `crbd-naive.wppl` in the `webppl/phywppl/examples/` directory, and the similarly named model script in the `webppl/phywppl/models/` directory. The simulation script presented here differs in four details from the model script in the repository. First, the script in the repository accommodates the possibility of incomplete sampling of the leaves in the tree. Thus, there is an additional parameter $\rho$ in the model, encoded as the variable `rho` in the script. This variable appears as an argument to all simulation functions. The `goesExtinct` function needs to take the sampling probability into account, and is aptly renamed to `goesUndetected`.

Second, the definitions of the `simTree`, `simBranch` and `goesUndetected` functions are hidden inside the `simCRBDNaive` function. This allows us to use the same generic names for these functions in all diversification models; only the simulation function needs to have a unique name. Hopefully, this facilitates for readers to recognize how we extended the basic template to accommodate the other diversification models.

Third, the script in the repository employs guards against extreme values of the `lambda` variable, which can otherwise cause problems with numerical exceptions or stalled simulations. We solve these problems by assigning zero weight to the simulation if the `lambda` value is above or below certain threshold values. We verified that the discarded simulations have negligible impact on the inference for all the examined models using the chosen guard values.

Finally, unlike the simple script described here, the script in the repository corrects for survivorship bias as explained in the next section. Before moving on to this, we want to point out that the naive CRBD simulation is suitable mainly for exploratory analyses of small trees. For efficient inference in WebPPL on phylogenetic diversification models for larger trees, it is important to manually modify the scripts so that they support aligned SMC inference (see Section 6.1). The CRBD model is the only model for which we provide an unaligned ("naive") model script.

#### 5.3 Correcting for survivorship bias

As discussed above (Section 3.3), if we condition the simulation on the age of the MRCA, we implicitly condition on the survival of the two subtrees originating at this point in time. To do this in a probabilistic program, we need to divide the probability of a simulation by $S(t_{\text{MRCA}}, \theta)^2$, that is, the square of the probability that the process survives (and is sampled) if it starts at $t_{\text{MRCA}}$, and the model parameter values are $\theta$. If $S(t, \theta)$ is not available in closed form, this is potentially cumbersome because it involves a sum and integral over an infinite number of realizations of the process for each simulation. However, we can solve this by observing that the division by $S(t_{\text{MRCA}}, \theta)^2$, which we cannot evaluate in general, can be rewritten as follows:

$$p(\theta|\psi, \text{survival}) \propto \frac{p(\theta)p(\psi|\theta)}{S(t_{\text{MRCA}}, \theta)^2} = p(\theta)p(\psi|\theta) \sum_{M=1}^{\infty} M \; (1 - S(t_{\text{MRCA}}, \theta)^2)^{M-1} \; S(t_{\text{MRCA}}, \theta)^2. \tag{10}$$

This shows that we can correct for the survivorship bias by using the generative model encoded in the function `goesExtinct` (or `goesUndetected`) to simulate two evolutionary processes starting at $t_{\text{MRCA}}$. We repeat this until both simulations survive to the present time, and multiply the weight of the rest of the simulated diversification process along the observed tree by the number of repetitions required to achieve this.

In WebPPL, we use the following recursive function to compute the number of simulations required until both trees survive:

```
var M_goesExtinct = function( t, lambda, mu )
{
   if ( !goesExtinct( t, lambda, mu ) && !goesExtinct( t, lambda, mu ) )
      return 1
   else
      return 1 + M_goesExtinct( t, lambda, mu )
}
```

The following lines are then inserted at the end of the simulation function `simCRBDNaive` to correctly condition on the survival of the two subtrees defining the MRCA:

```
var M = M_goesExtinct( t, lambda, mu )
factor( Math.log( M ) )
```

The script in the repository is slightly more complex because we take incomplete sampling into account, and also implement a guard against an excessive number of repetitions.

#### 5.4 Scripts for other diversification models

Example analysis scripts for all models are provided in the directory `webppl/phywppl/examples/`, and generic model simulation scripts in the directory `webppl/phywppl/models/`. All simulation scripts we provide in the latter directory are set up to trigger aligned SMC inference in WebPPL. As mentioned above, the only exception is the CRBD model, where we provide both a naive, unaligned version (`phywppl/models/crbd-naive.wppl`) and an aligned version (`phywppl/models/crbd.wppl`) for instructional and testing purposes. We provide both scripts using analytical likelihoods and scripts using explicit simulation for all simple diversification models (CRB, CRBD, TDB, TDBD). The scripts for the CRB, TDB and TDBD models involve simple and straightforward modifications of the corresponding scripts for the CRBD model, described above.

The model scripts for all lineage-specific diversification models (ClaDS0, ClaDS1, ClaDS2, LSBDS, BAMM) follow the template described above for the CRBD model, including the modifications needed to trigger aligned SMC inference. Analogous component functions are used in the simulation scripts; they are even named the same except for the main simulation function, which is named after the corresponding diversification model.

In probabilistic programming, you have to be explicit about the model variables that you want to estimate. These are the variables that are returned from the model function. The focus in our study was on the normalization constant and the main model parameters. Therefore, our model simulation scripts do not have to return anything, as all relevant parameters are defined already in the analysis scripts in the `webppl/phywppl/examples/` folder. However, readers may well be interested in sampling the outcome of a diversification process along the tree. For instance, it may be desirable to analyze parameters such as the number and location of change events on different lineages, or the mean speciation rate for individual branches in the tree. To facilitate such analyses, we give an example model script for the ClaDS2 model returning the entire reconstructed tree, with descriptions of the outcome of the simulation process for each branch and node in the tree in extended Newick format. This script is found in the `webppl/phywpppl/models/` folder.

#### 5.5 Birch model scripts

Birch is an object oriented probabilistic programming language. It uses more concise syntax than WebPPL for the probabilistic constructs. For example, the assume statement in the form

```
x ~ Exponential(1);
```

is used to express that a random variable (`x` in the example above) is distributed according to a given probability distribution (an exponential distribution with rate 1). Execution of such a statement depends on whether the variable has a value or not. If it has, its behavior is equivalent with an observe statement; if not, the variable is associated with the given distribution. Birch uses delayed sampling, so a concrete value might not be sampled until needed. Birch also supports explicit sample and observe statements. To draw a value from an exponential distribution with rate $\lambda$, we would write

```
t <~ Exponential(λ);
```

To state that an outcome of a random variable distributed according to a Poisson distribution with rate $\lambda$ is 0, we would write

```
0 ~> Poisson(λ);
```

The factor statement in WebPPL corresponds to `yield FactorEvent(log_factor)`. To simplify the diversification model definitions, we have defined two helper functions for commonly used `yield` statements.

```
yield duple();
```

corresponds to

```
yield FactorEvent(log(2));
```

and is used to account for the rotational factor at hidden speciation events. Similarly,

```
yield impossible();
```

is the same as

```
yield FactorEvent(-inf);
```

and it is used when simulated side branches resulting from hidden speciation do not go extinct, that is, when they are incompatible with the observed tree. Note that `yield impossible()` statement also ceases the execution of the particle.

As we have mentioned above, Birch is an object-oriented language and the models take advantage of this. For instance, the CRBD model script defines a `CRBDModel` class, which is derived from a base class called `PhyModel`. Let us examine a somewhat simplified version of the `CRBDModel` class definition (Algorithm 4), to see how it compares to the WebPPL script.

**Algorithm 4** CRBDModel class definition in Birch (somewhat simplified)

```
class CRBDModel < PhyModel<PhyNode, PhyParameter> {
  λ_k:Real;
  λ_θ:Real;
  ε_min:Real;
  ε_max:Real;
  ρ:Real;

  fiber initial() -> Event {
    super.initial();
    θ.λ ~ Gamma(λ_k, λ_θ);
    θ.ε ~ Uniform(ε_min, ε_max);
  }

  fiber step() -> Event {
    count:Random<Integer>; // number of (hidden) speciation events
    count ~ Poisson(θ.λ * (node.t_beg - node.t_end));
    for i in 1..Integer(count) {
      t:Random<Real>;
      t ~ Uniform(node.t_end, node.t_beg);
      simulateUnobserved(t);
      yield duple();
    }

    0 ~> Poisson(θ.λ * θ.ε * (node.t_beg - node.t_end));

    if node.isSpeciation() {
      0.0 ~> Exponential(θ.λ);
    }
  }

  fiber simulateUnobserved(t_beg:Real) -> Event {
    Δ_d:Random<Real>; // waiting time until an extinction event
    Δ_d ~ Exponential(θ.λ * θ.ε);
    t_d:Real <- t_beg - Δ_d;
    if t_d < 0 {
```

```
      // Species survived to the present time
      yield impossible();
    }

    count:Random<Integer>; // number of speciation events
    count ~ Poisson(θ.λ * (t_beg - t_d));
    for i in 1..Integer(count) {
      t:Random<Real>;
      t ~ Uniform(t_d, t_beg);
      simulateUnobserved(t);
    }
  }
}
```

The `CRBDModel` class is derived from the class `PhyModel`, which is a templated class. The base class takes care of tasks that are common to all diversification models, such as walking over the tree. This is analogous to the recursive calls in the `simTree` function in the WebPPL script (Algorithm 3), which also walk over the branches in the tree. At the top of the class definition, the member variables and their types are declared. These are the parameters of the prior distributions for the model variables $\lambda$ and $\epsilon$. The parameters are assigned specific values when the class is instantiated in connection with running the program. The inference settings and input values for the analyses are in the `config/crbd.json` file and in each of the `input/<name of tree>.json` input files.

Instead of member functions, the class defines several member fibers. A `fiber` (also known as a coroutine) is similar to a function, but the execution might be paused (e.g., to resample the particles) and resumed. The `initial` fiber initializes the simulation by assuming `lambda` and `epsilon` to be distributed according to the appropriate priors. Note that these model variables are packaged inside an object called θ.

The `step` fiber corresponds to the `simBranch` function in the WebPPL script. In Birch, we use a different method for simulating the speciation and extinction events than in WebPPL. Rather than drawing the waiting times between hidden speciation events, we use the fact that the number of hidden events is described by a Poisson distribution, and the event positions are uniformly distributed over the branch length. This simulation method is faster than drawing each of the waiting times. In the line

```
0 ~> Poisson(θ.λ * θ.ε * node.branch_length);
```

we condition on the fact that there are 0 extinction events on the branch (recall that $\mu = \lambda\epsilon$). In WebPPL, we used a `factor` statement with the appropriate probability instead, which is an alternative way of accomplishing the same thing. Finally, in the line

```
0.0 ~> Exponential(θ.λ);
```

we condition on there being a speciation at the end of the branch (if it ends in an interior node). Equivalently, we could have factored in $\log \lambda$, as we did in WebPPL, with a `yield` statement.

The `simulateUnobserved` fiber corresponds to the `goesExtinct` function in WebPPL. However, here we first simulate the time until the branch goes extinct. If the branch does not go extinct, we set the weight to zero, effectively killing off the simulation. If it does go extinct, we simulate the hidden speciation events along the branch, and call `simulateUnobserved` recursively for each of those events.

The code described above is subject to change, as Birch is developing rapidly. However, this section illustrates the basic Birch features, and how they can be used to code diversification models efficiently. Hopefully, it also sheds additional light on general PPL concepts, as it gives alternative but equivalent ways of coding some model elements compared to the WebPPL scripts we have seen previously.

### 6 Inference

In this section, we provide additional details on the non-standard algorithms we used to allow efficient PPL inference on phylogenetic diversification models.

#### 6.1 Alignment

The encoding of the CRBD model given in Section 5.2 is rather natural—it is simply a description of the birth-death process, with a few calls to `factor` to correct for some probability effects that we do not model explicitly. Unfortunately, the default SMC algorithm implemented in WebPPL is quite inefficient for this naive implementation of the birth-death process. The algorithm *always* resamples particles (simulations are called *particles* in the SMC algorithm) at calls to `factor` and `condition`. Since, for every execution of the program, there is a different number of hidden speciation events on each branch in the observed tree, this will cause the SMC particles to get out of sync at resampling points. Particles that have few hidden speciation events may reach the end of the simulation long before particles that have many hidden speciation events. Thus, if we always resample at hidden speciation events, we will be comparing particles that can be at very different points in the simulation.

Intuitively, one might expect that it would be better to compare the particles only when they reach the same points in the probabilistic program. We call this *alignment* of the SMC resampling points. In the diversification models, we could, for instance, make sure that the resampling occurs only at the branching points in the observed tree. To explore this idea, we "tricked" the

SMC algorithm in WebPPL to align the resampling points by introducing a few modifications to the birth-death simulation in the `simBranch` and `simTree` functions, as illustrated in the code below (compare to the naive CRBD simulation presented above in Algorithm 3):

**Algorithm 5** A complete WebPPL script for simulating CRBD.

```
var simBranch = function( startTime, stopTime, lambda, mu )
{
   var t = exponential ( {a: lambda} );

   var currentTime = startTime - t;

   if ( currentTime <= stopTime )
      return 0.0;

   var sideDetection = crbdGoesUndetected( currentTime, lambda, mu )
   if ( sideDetection == false )
      return ( -Infinity )

   return simBranch( currentTime, stopTime, lambda, mu )
         + Math.log( 2.0 );
}

var simTree = function( tree, parent, lambda, mu )
{
   var lnProb1 = - mu * ( parent.age - tree.age );

   var lnProb2 = ( tree.type == 'node' ? Math.log( lambda ) : 0 );

   var lnProb3 = simBranch( parent.age, tree.age, lambda, mu );

   factor( lnProb1 + lnProb2 + lnProb3 )

   if ( tree.type == 'node')
   {
      simTree( tree.left, tree, lambda, mu )
      simTree( tree.right, tree, lambda, mu )
   }
}
```

Specifically, we need the WebPPL SMC implementation to skip the resampling induced at the calls to `factor` and `condition` within `simBranch` in the naive model script. We achieve this by replacing the `factor` and `condition` statements in the `simBranch` function by code that accumulates the weight and returns it to `simTree`. The accumulated weight is then passed as an argument to `factor` in `simTree`, after the entire branch has been processed, triggering resampling at this point. The `factor` statement is also passed the probability of no extinction on the branch (`lnProb1`), and the likelihood of a speciation at the end of the branch, if it is an interior branch in the observed tree (`lnProb2`). Note that, to improve efficiency, we immediately return `-Infinity` in `simBranch` if a call to `goesExtinct` returns false, since there is no need to continue the recursion if this occurs. By modifying the simulation script in this way, the SMC particles stay in sync. There are no triggers of resampling in the `simBranch` recursion, so resampling is always performed in `simTree`, in between processing branches of the observed tree.

Simulations on a few example trees of varying sizes confirm that this indeed improves SMC efficiency on diversification models considerably (Supplementary Figure 3). The larger the tree, the more important it is for SMC performance to align the resampling points in this way. Ideally, one should not have to manipulate model scripts in the way described above; alignment should be applied automatically when it improves SMC efficiency. This is an idea that we are exploring within the TreePPL project. The goal is to analyze the potential performance gains induced by resampling, and then apply it intelligently either in the compiler and/or the language runtime. We separately present a static analysis for automatic alignment of programs[39]. Note that alignment is not *guaranteed* to improve accuracy—in certain cases, it might actually degrade performance. However, for all models considered here, alignment is beneficial.

#### 6.2 Delayed sampling

Probabilistic computations involve not only simulation and observation, as represented by the `sample` and `observe` statements in a PPL, but also such computations as marginalization, enumeration, and conjugate updating.

Consider the following joint distribution between two variables $x$ and $\lambda$:

$$p(x, \lambda) = p(x \mid \lambda)p(\lambda), \tag{11}$$

where the two factors on the right are encoded in the probabilistic program as, for example:

$$\lambda \sim \text{Gamma}(1, 1),$$
$$x \sim \text{Poisson}(\lambda).$$

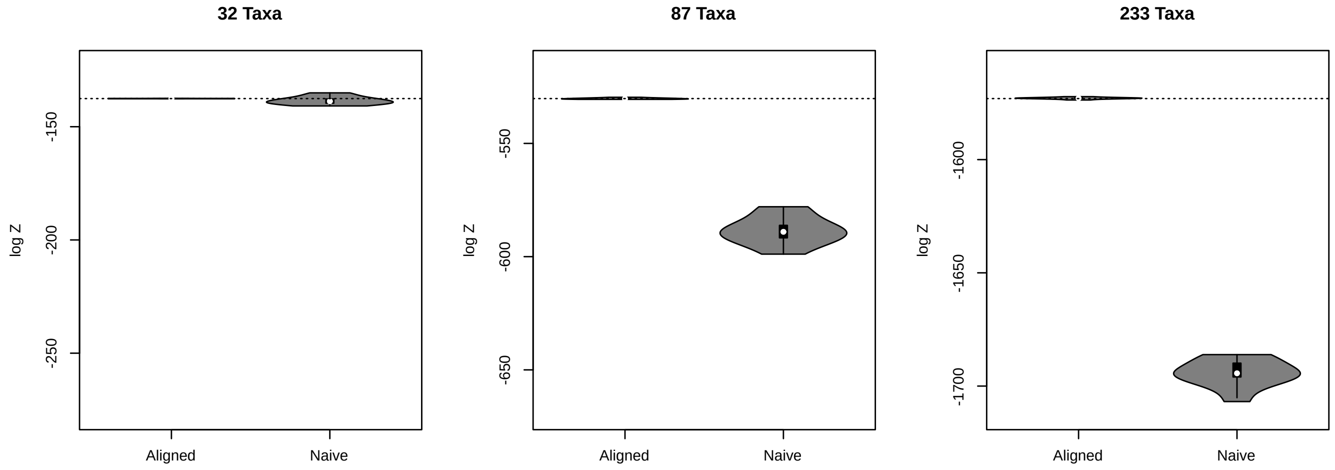


**Fig. 3** A comparison of the precision in the estimated normalization constant between naive and aligned CRBD. Left: 32-taxon tree (Bisse_32). Center: 87-taxon tree (Cetaceans_87). Right: 233-taxon tree (Primates_233). SMC inference with 10,000 particles in WebPPL. Dotted line: exact analytical solution. Parameters: $\lambda = 0.2$, $\epsilon = 0.5$, complete sampling of leaves assumed ($\rho = 1$).

We may wish to compute the marginal distribution of $x$:

$$p(x) = \int p(x \mid \lambda)p(\lambda)\,\mathrm{d}\lambda, \tag{12}$$

or, given a value of $x$, compute the posterior distribution over $\lambda$:

$$p(\lambda \mid x) = \frac{p(x \mid \lambda)p(\lambda)}{p(x)}. \tag{13}$$

Evaluations such as these can be performed analytically for random variables with a conjugate relationship (such as the gamma-Poisson relationship in the example above), or for discrete random variables where all possible outcomes can be enumerated. This can improve the performance of inference by, for example, reducing the variance in statistical estimators, such as that for the marginal likelihood.

Delayed sampling[6] is a particular heuristic that may be employed by a PPL to identify and leverage such situations to improve inference outcomes. It does so in a manner that produces correct results, even for programs with stochastic branches and unbounded recursion as may be encountered in Turing-complete programming languages. It is not necessary for the programmer to painstakingly code such computations by hand.

We have used delayed sampling extensively in this work, substantially reducing the variance in marginal likelihood estimates. In particular, for Poisson processes on trees, gamma prior distributions over rates are conjugate either to the Poisson-distributed number of events in a given time interval, or the exponentially-distributed time between events. These rate parameters are then automatically marginalized out by delayed sampling, substantially reducing variance in the marginal likelihood estimate for these models. The same approach to handling parameters is used in Kudlicka et al.[40] and Wigren et al.[41]. Delayed sampling is only available in Birch at this point.

#### 6.3 Alive particle filter

The resampling step in SMC amounts to drawing $N$ samples (with replacement) from the current set of $N$ particles with probabilities proportional to their weights. While simulating the evolution of unobserved side branches, if any species survives to the present day (and is sampled), the weight of the particle must be set to 0. This leads to sample impoverishment—there are fewer particles to choose from during resampling. In extreme cases, where all particles have zero weight, there are no particles to choose from at all, and the algorithm fails. This can be a serious problem for SMC inference on diversification models when the likelihood of extinction of side branches is low. For instance, this can occur if the net diversification rate ($\lambda - \mu$) is high.

The extended alive particle filter[40]—the development of which from the initial version of this algorithm[42] was inspired by phylogenetic diversification models—solves these two problems by replacing the particles with zero weights with new samples drawn from the particle set at the previous time step (again with probabilities proportional to the particle weights at that time) and repeating the propagation step (the simulation from the previous resampling point until the current resampling point). This replacing procedure is repeated until the weights of all particles are positive. Note that in order to estimate the marginal likelihood without bias, one needs to repeat this procedure for one additional particle. However, with a reasonable number of particles, this extra computational cost is negligible, and we therefore applied the alive particle filter to all analyses. The alive particle filter is only available in Birch at this point.

##### 6.4 Tree orientation

During the course of the study, we discovered that the orientation of the nodes in the observed tree can have a noticeable influence on the efficiency of SMC inference for some trees. The effect appears to be associated with highly imbalanced trees, which may be oriented such that left and right subtrees systematically have different properties. A depth-first SMC algorithm can apparently become misled by the imbalance between left and right descendants in such trees, so that early resampling events can select particles that do not do well towards the end of the simulation, decreasing the quality of the final estimate. We found that orienting all nodes such that the descendant branch with the shortest subtree length was always processed first solved this problem. Thus, all trees were reoriented in this way before final analyses in Birch.

#### 7 Verification

We performed a wide range of experiments to verify that the model scripts are correct. For all tests involving WebPPL or third-party software, the full set of experiments—including the source code, data, graphs and reports—can be found in the directory `verification` of the `phywppl` package. The verification experiments involving Birch were performed by changing the input files and/or models to fix the values of selected parameters. The results from these experiments are included in the above-mentioned directory, together with the results from the experiments involving WebPPL.

Here, we only present a summary of the experiments. They all use the 32-taxon example tree, which we provide as one of the builtin trees in the `phyjs` package. The tree has been previously used as an example in diversification model papers; it is originally from the Mesquite software[19] but does not appear to have been published separately. Only scrips adapted for aligned SMC inference were used in the verification experiments.

The experiments are based on several lines of attack. In the first round of tests, we used the fact that there are analytical solutions for the likelihood of the simple diversification models (CRB, CRBD, TDB and TDBD) under specific parameter values. Thus, we could verify that the normalization constant computed by SMC from our explicit simulation scripts for the same models (the scripts that simulate the process along the tree instead of calling the likelihood function) matched the corresponding analytical likelihoods for a wide range of specific parameter values. These tests are important because the explicit simulation scripts for the simple models served as templates for the scripts describing the more complex models.

The second round of tests were based on the observation that the more complex diversification models (ClaDS0, ClaDS1, ClaDS2, LSBDS and BAMM) all collapse to simpler models with analytically known likelihoods under specific parameter settings. This allows us to verify that the normalization constant computed from the scripts for the complex models matched the corresponding analytical likelihoods for select points in parameter space.

For other points in parameter space, we cannot verify the scripts for the more complex models against analytical likelihoods, but we can use other approaches to test their correctness. For instance, the WebPPL and Birch scripts for the complex models were implemented independently by different co-authors of this paper, and the inference algorithms in WebPPL and Birch were also different and based on independent implementations. In the third round of tests, we verified that the WebPPL and Birch scripts for the complex models gave the same normalization constant for a grid of parameter values despite these differences.

The fourth round of tests took advantage of the independent implementations available in third-party software for the ClaDS models[43] and for LSBDS[29]. We verified that our scripts for these models resulted in the same estimates of the likelihood as these implementations for a select set of parameter values, despite being based on entirely different code bases and computational strategies.

Third-party software also exists for BAMM[38] but it does not compute correct likelihoods for the model[32], so it cannot be used to verify our scripts. However, the BAMM model collapses to the LSBDS model when all $z^i = 0$. We therefore verified that our BAMM model script results in the same likelihood estimates as the LSBDS model script under select parameter values matching this constraint but lacking analytical solution.

##### 7.1 Simple models against analytical likelihoods

All simulation scripts for simple models (CRB, CRBD, TDB and TDBD) generated normalization constant estimates that matched the corresponding analytical likelihoods very closely. We observed some variance in the estimates for high $\lambda$ values, but these parameter values have low likelihood and are thus less important for inference (Supplementary Figure 4). All $\lambda \times \epsilon$ combinations for two values of $\rho$: $\rho = 0.5$ (incomplete sampling in the order of magnitude of the $\rho$-range for the bird trees) and $\rho = 1$ (complete sampling) have been checked.

##### 7.2 Complex models against analytical likelihoods

All model scripts for advanced diversification models (ClaDS0, ClaDS1, ClaDS2, LSBDS and BAMM) generated normalization constant estimates that matched analytical likelihoods under parameter settings for which closed solutions exist (Supplementary Figure 5, 6). Again, we checked all $\lambda \times \epsilon$ combinations for two values of $\rho$: $\rho = 0.5$ (incomplete sampling in the order of magnitude of the $\rho$-range for the bird trees) and $\rho = 1$ (complete sampling).

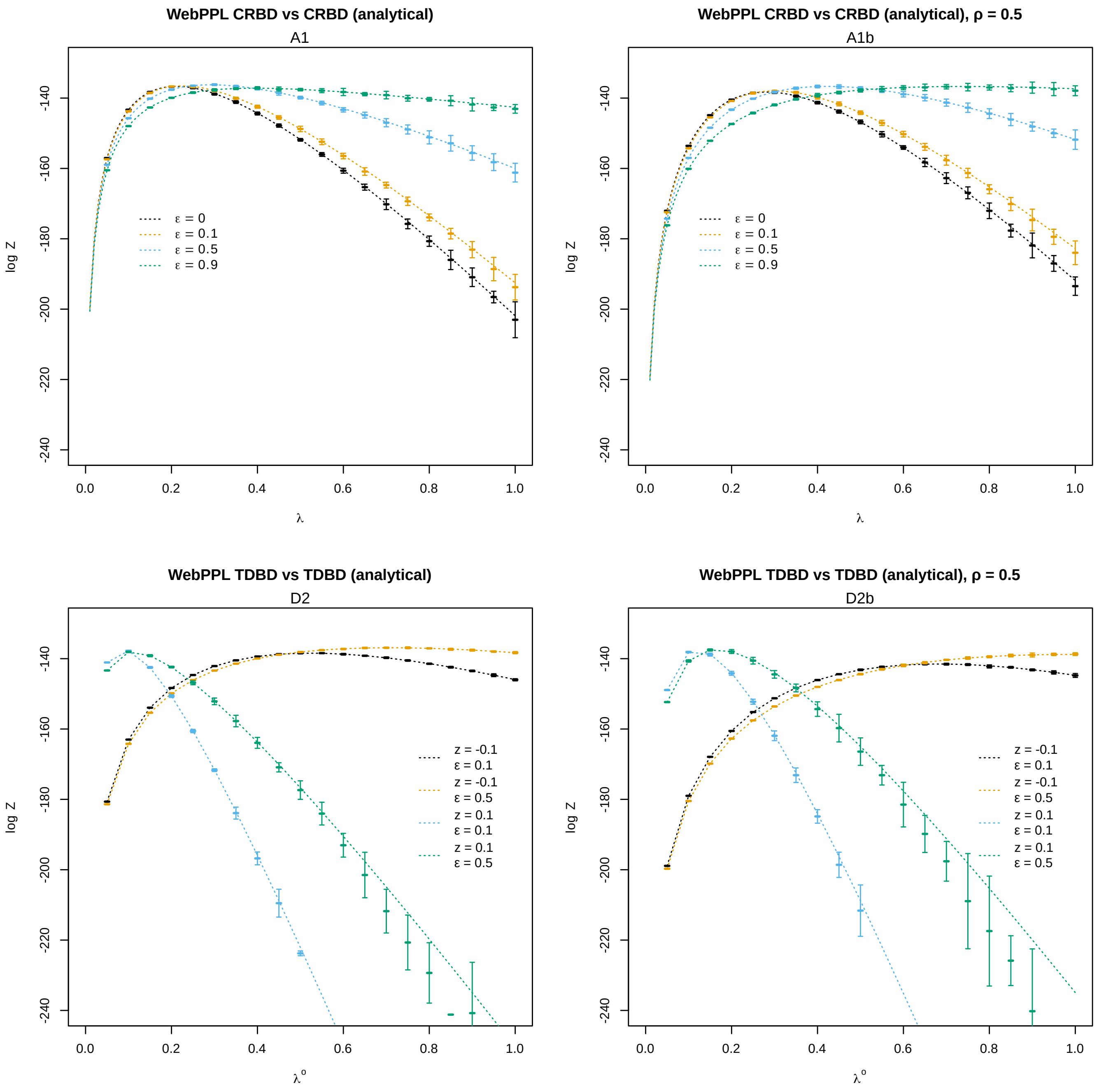


**Fig. 4** Verification of explicit simulation scripts (WebPPL) for simple diversification models: normalization constants match analytical likelihoods for select parameter values. Error bounds: ±2 standard deviations. Experiment codes in the GitHub repository indicated under the main title.

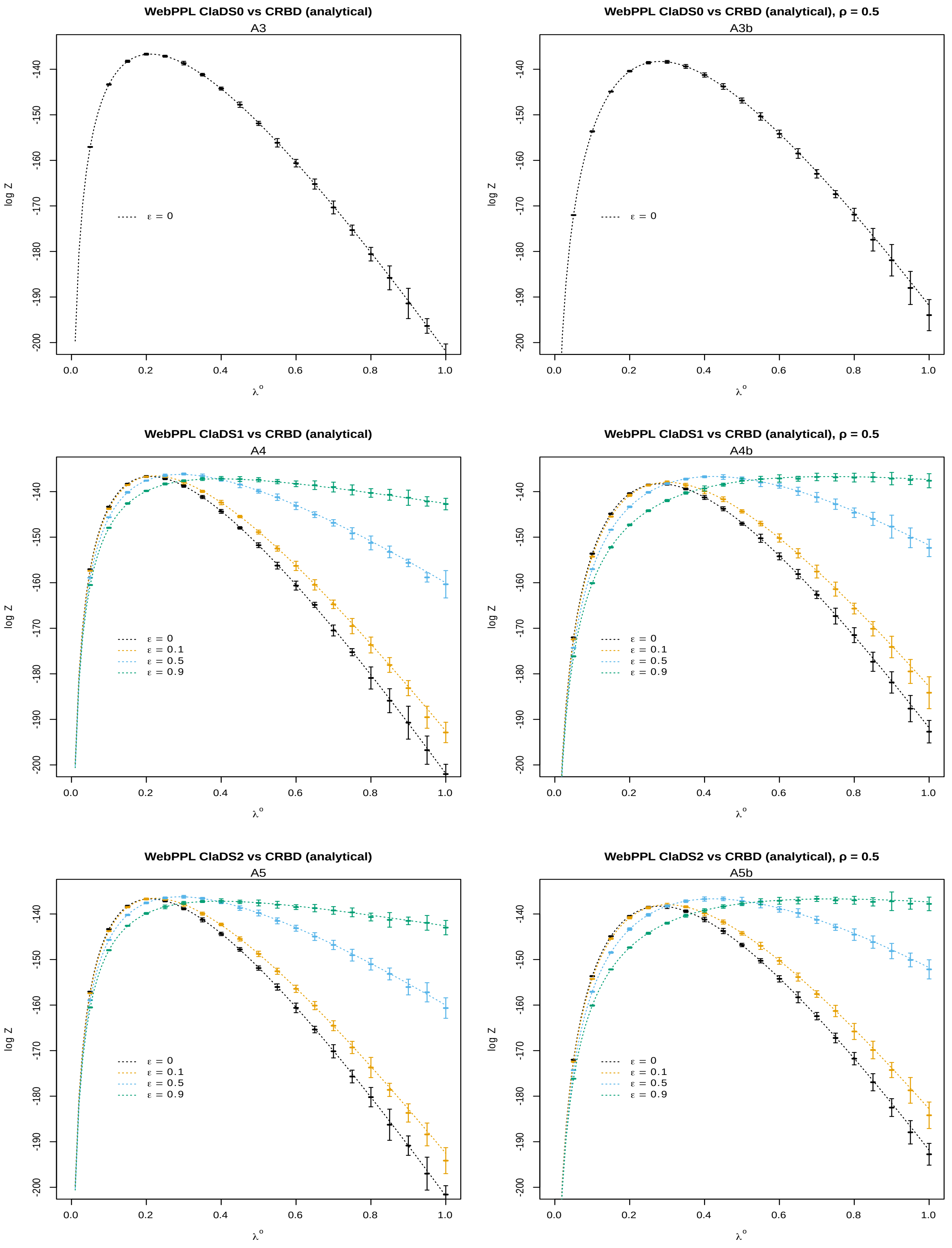


**Fig. 5** Verification of WebPPL simulation scripts for the ClaDS[0-2] models: normalization constants match analytical likelihoods for select parameter values. Error bounds: $\pm 2$ standard deviations. Experiment codes in the GitHub repository indicated under the main title.

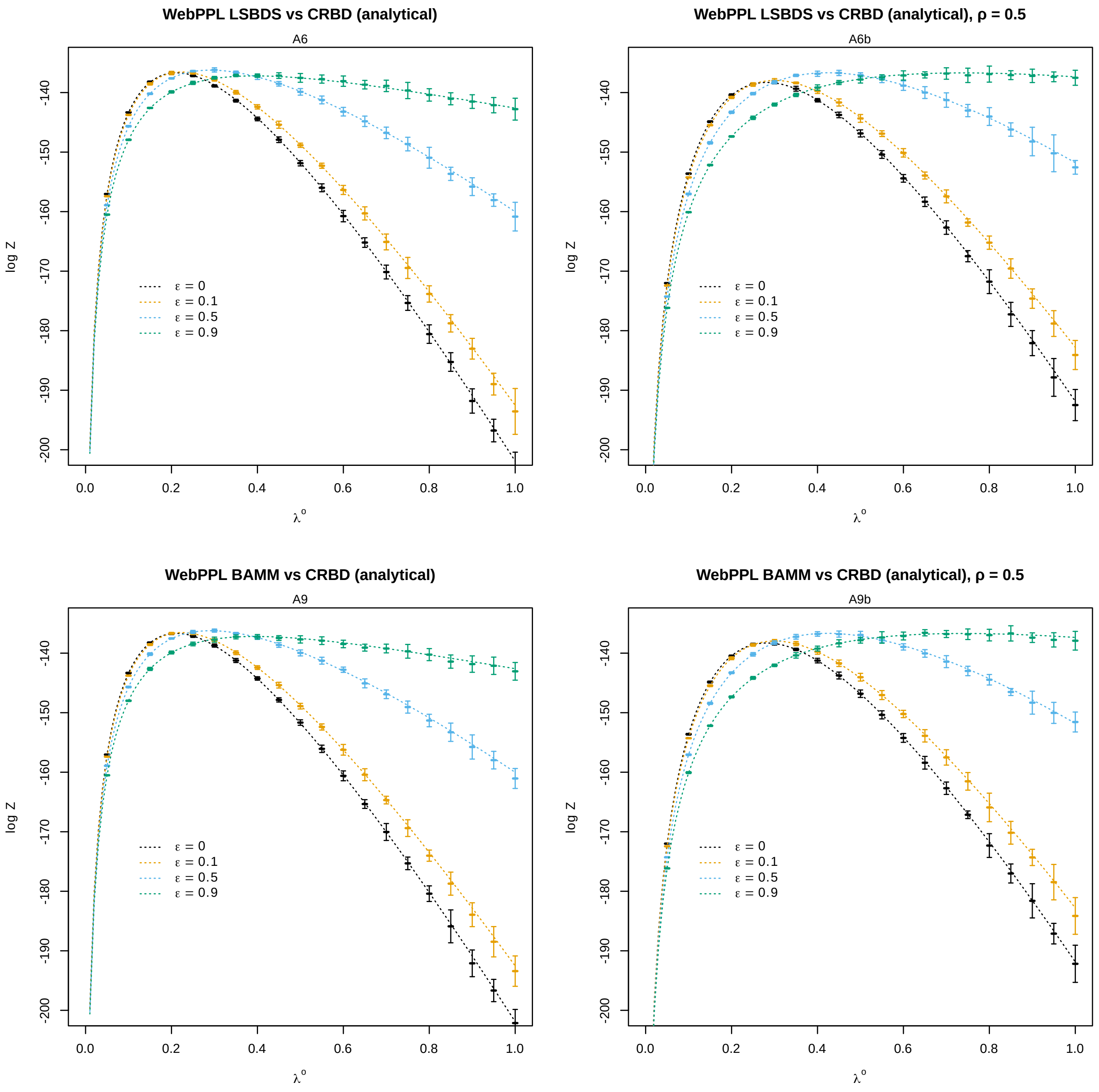


**Fig. 6** Verification of WebPPL simulation scripts for lineage-specific diversification models (complex models): normalization constants match analytical likelihoods for select parameter values. Error bounds: $\pm 2$ standard deviations. Experiment codes in the GitHub repository indicated under the main title.

**Table 5** Cross-verification of WebPPL and Birch for $\rho < 1$.

| | | WebPPL | | Birch | |
|---|---|---|---|---|---|
| Model | $\rho$ | mean log $Z$ | std. dev. log $Z$ | mean log $Z$ | std. dev. log $Z$ |
| BAMM | 0.1 | -146.839 | 0.623 | -146.608 | 0.279 |
| BAMM | 0.5 | -139.612 | 0.198 | -139.544 | 0.089 |
| ClaDS0 | 0.1 | -152.348 | 0.921 | -150.540 | 0.663 |
| ClaDS0 | 0.5 | -142.506 | 0.450 | -142.528 | 0.173 |
| ClaDS1 | 0.1 | -153.957 | 2.003 | -151.245 | 0.635 |
| ClaDS1 | 0.5 | -143.422 | 0.351 | -143.385 | 0.164 |
| ClaDS2 | 0.1 | -152.426 | 1.578 | -151.747 | 0.915 |
| ClaDS2 | 0.5 | -143.085 | 0.315 | -143.000 | 0.161 |
| CRBD | 0.1 | -172.826 | 0.093 | -172.795 | 0.089 |
| CRBD | 0.5 | -143.259 | 0.083 | -143.314 | 0.077 |
| LSBDS | 0.1 | -172.794 | 0.120 | -172.819 | 0.095 |
| LSBDS | 0.5 | -143.361 | 0.075 | -143.321 | 0.032 |
| TDBD | 0.1 | -145.567 | 0.989 | -145.042 | 0.063 |
| TDBD | 0.5 | -139.039 | 0.263 | -139.028 | 0.000 |

#### 7.3 Birch and WebPPL cross-verification

Under parameter and prior settings for which closed solutions do not exist, the independently developed Birch and WebPPL scripts for advanced diversification models resulted in matching normalization constant estimates. Birch estimates were slightly more precise than WebPPL estimates (Supplementary Figure 7) but this is expected given the more powerful inference algorithms used by Birch. In the tests illustrated in the figure, $\rho = 1$ is assumed. To verify that our code is correct also for $\rho < 1$, we ran several additional grid point experiments as summarized in Table 5. All experiments in the table were conducted on the 32-taxon tree using the standard priors, except for $\lambda = 0.2$, $\epsilon = 0.5$, and $\rho$, for which point values were used instead. For TDBD, the Birch implementation uses the analytical solution.

#### 7.4 Verification of ClaDS models against RPANDA

Verification of the probabilistic programs and PPL inference algorithms described in this paper against the reference RPANDA implementation of the ClaDS models is quite involved, and would not have been possible without extensive help from the author of the ClaDS code in RPANDA (Odile Maliet), as computation of the likelihoods with RPANDA is not part of its public API. RPANDA computes the likelihood for points in parameter space where all the initial $\lambda^i$ values for the branches in the reconstructed tree are known, as well as the $\lambda^o$ value, pertaining to the MRCA of the tree. The likelihood in RPANDA is also conditioned on specific values of the model parameters $\alpha$ and $\sigma$, as well as on $\mu$ (for ClaDS1) or $\epsilon$ (for ClaDS2). The ClaDS0 likelihood function in RPANDA is based on analytical equations, while the ClaDS1 and ClaDS2 functions are based on numerical approximations using a variety of techniques. In addition, the functions only give the density up to a proportionality constant, further complicating direct comparisons with our scripts.

The RPANDA setup means that the PPL scripts have to condition on specific values for all of the model parameters, including the initial $\lambda$ values for all branches, to emulate the RPANDA likelihood computations. To be able to conduct the verification experiments, we decided to use a fixed value $\lambda_f$ for $\lambda^o$ and all $\lambda^i$ parameters of the model in our WebPPL scripts; we did not attempt to perform these verification experiments in Birch. We then chose a range of $\lambda_f$ values, and explored these points in parameter space under some specific values of $\alpha$, $\sigma$ and $\mu$ (for ClaDS1) and $\epsilon$ (for ClaDS2). Likelihoods for the same points in parameter space were then computed in RPANDA with the analytical likelihood function (for ClaDS0) and the numerical solvers (for ClaDS1 and ClaDS2). In the git repository accompanying the paper, we provide both the WebPPL scripts emulating the RPANDA computations and the R scripts we used to compute likelihoods for the corresponding points in parameter space with RPANDA.

For ClaDS0, the initial experiments showed that the likelihood function in RPANDA computes densities that very closely match the densities expected for oriented and unlabeled trees. Thus, we concluded that the proportionality constant for the ClaDS0 likelihood function in RPANDA is the same as the conversion factor from densities on oriented and unlabeled trees to densities on labeled, unoriented trees. This factor is $L_p = \log(2^{(n-1)}/n!)$, where $n$ is the number of leaves in the tree (see Section 3.2).

When controlling for this, the likelihoods estimated by WebPPL for ClaDS0 are consistent with those computed by RPANDA (Supplementary Figure 8). For points in parameter space where ClaDS1 and ClaDS2 collapse to ClaDS0, that is, for points where $\mu = \epsilon = 0$, likelihoods estimated by WebPPL and RPANDA are also very similar. The same is true for small values of $\lambda$ and $\mu$) in ClaDS1, and for small values of $\lambda$ and $\epsilon$ in ClaDS2. For larger values, RPANDA apparently overestimates the likelihood for both models, and there are also some apparent discretization effects at very high values of $\lambda$. We tried to examine the effects of these inaccuracies in RPANDA on the posterior estimates of the ClaDS1 and ClaDS2 model parameters for the test tree, but were unable to get sufficiently good MCMC convergence in RPANDA to allow meaningful analysis of these results.

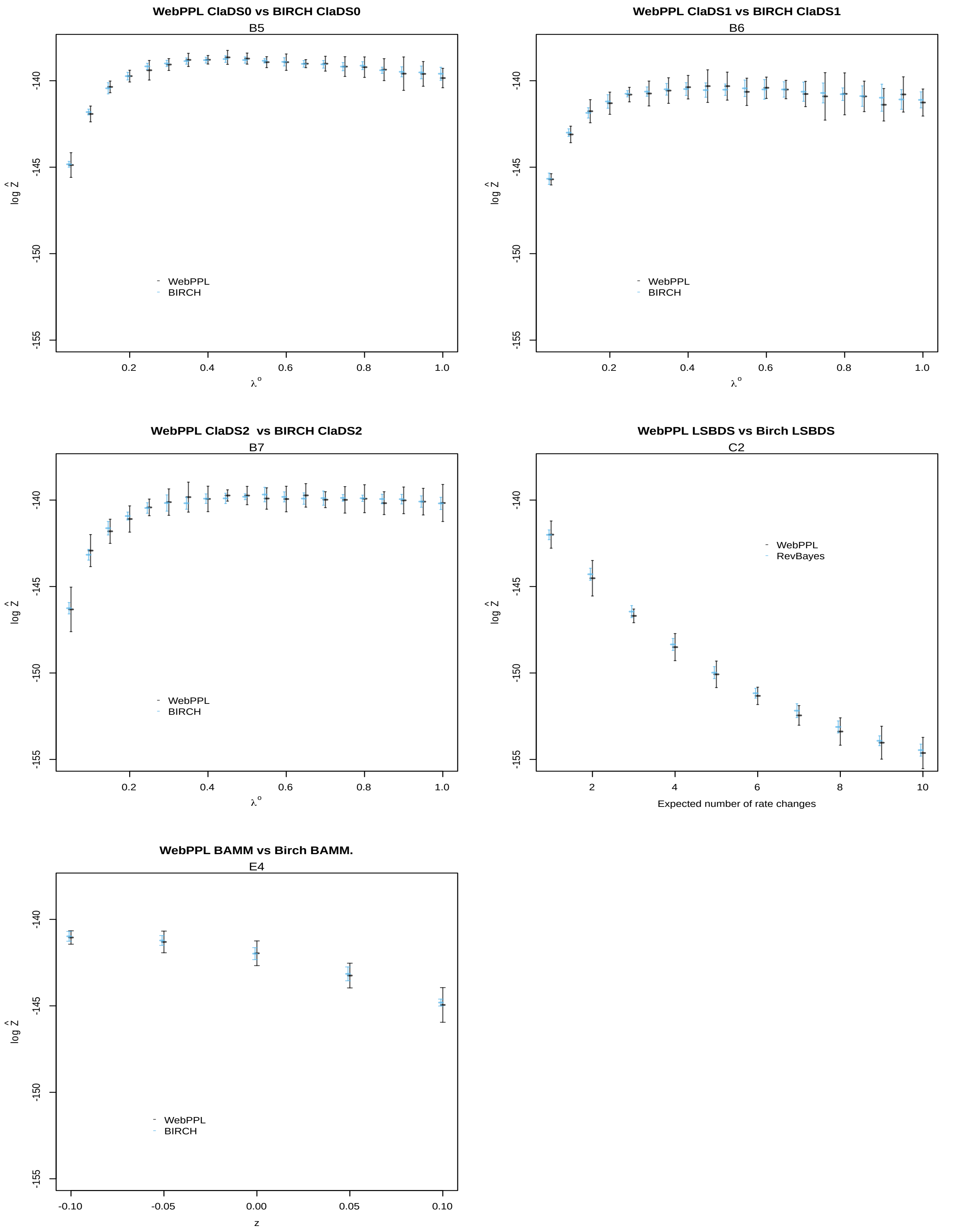


**Fig. 7** Verification that the WebPPL and Birch scripts for lineage-specific diversification models generate normalization constants that match each other for select parameter values.

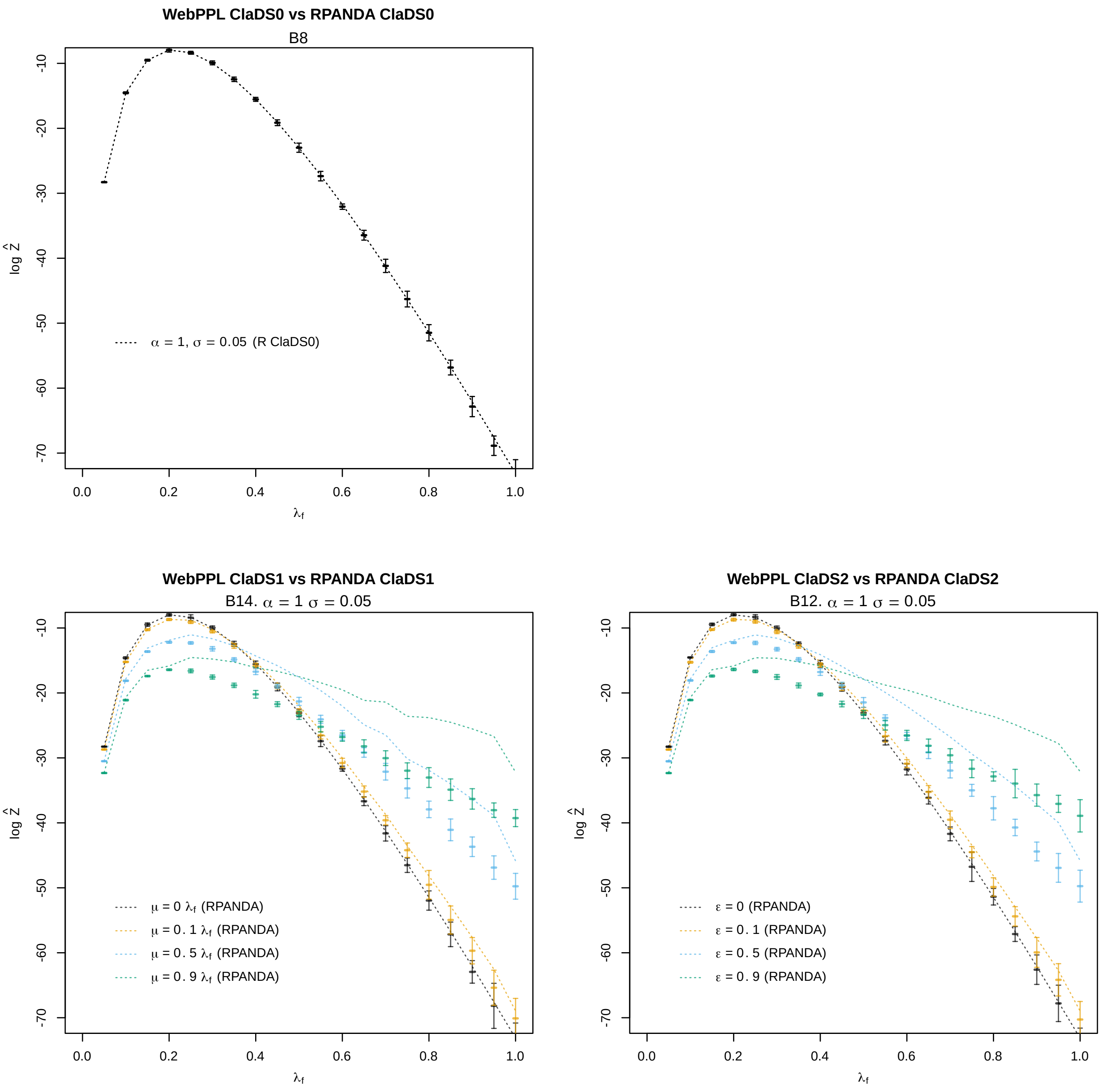


**Fig. 8** Verification of the likelihoods computed by WebPPL in programs emulating RPANDA against likelihoods computed by RPANDA for the ClaDS models.

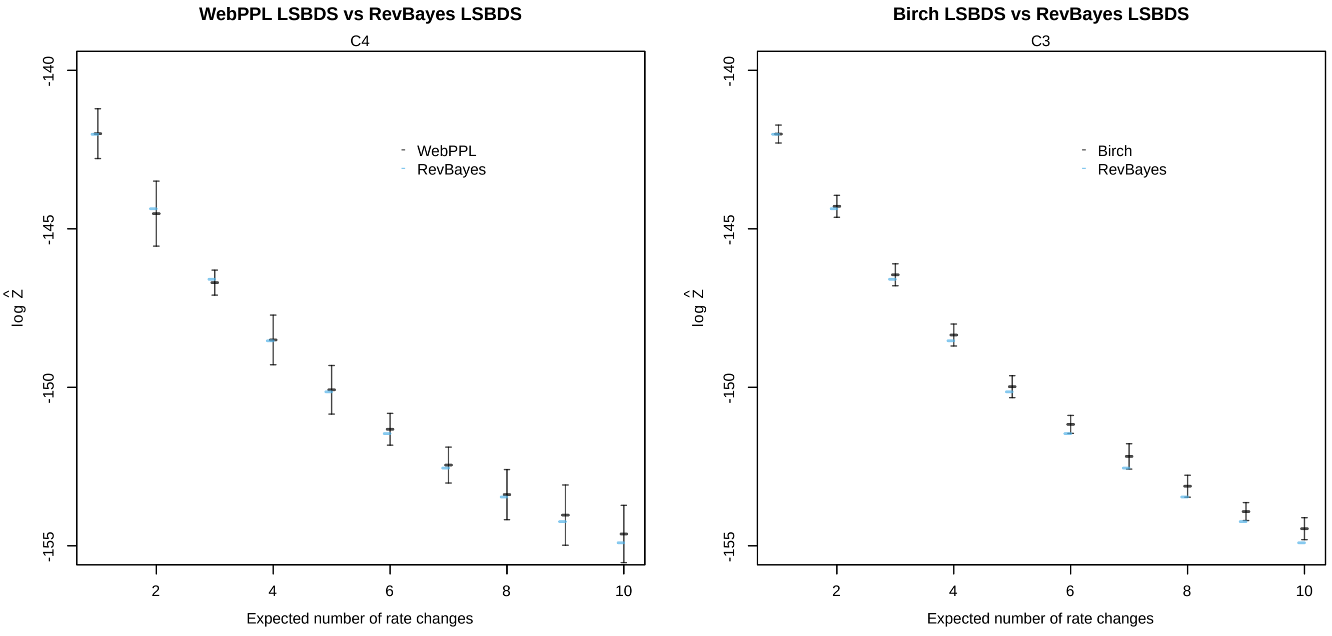


**Fig. 9** Verification that the WebPPL and Birch scripts for the LSBDS model generate normalization constant estimates that match the numerically estimated likelihood computed by RevBayes.

##### 7.5 Verification of LSBDS against RevBayes

In the current implementation of LSBDS in RevBayes (the SCM algorithm), the likelihoods are computed by discretizing the $\lambda$ and $\mu$ priors. Transitions happen by "jumping" from one pair of discrete values of $\lambda$ and $\mu$ to a different pair. We discovered that $\lambda$ and $\mu$ are coupled when these jumps are made: i.e., the discrete vectors representing the prior distributions $f_\lambda$ and $f_\mu$ have to be of the same length and, when a jump happens, a single new array index is chosen for both the $\lambda$ and the $\mu$ vector. Thus, usually, the RevBayes LSBDS examples[h] fix $\lambda$ but discretize $\mu$ (or vice-versa). However, it is possible to discretize both $\lambda$ and $\mu$ and then expand the two arrays so that all possible combinations of $\lambda$ and $\mu$ values appear when sweeping both vectors simultaneously with a single array index. This has to be done manually.

We verified our LSBDS scripts against RevBayes for specific values of $\eta$ and integrated out $\lambda \sim$ Exponential(1) and $\mu = \epsilon\lambda$, where $\epsilon \sim$ Uniform(0, 1) using $k = 10$ rate categories for both $\lambda$ and $\mu$. We implemented the appropriate vector of parameter values manually, as described above. The RevBayes scripts used in the verification experiments are provided in the git repository accompanying the paper.

Under these settings, both the WebPPL and Birch scripts for the LSBDS model generate normalization constant estimates that match the likelihoods computed by RevBayes (Supplementary Figure 9). As observed previously in several experiments, Birch provides slightly more precise estimates of the normalization constant than WebPPL.

##### 7.6 Verification of BAMM against LSBDS

There is no third-party software implementing BAMM that we can verify the WebPPL and Birch scripts against. However, we can use the fact that BAMM collapses to LSBDS when all $z^i$ values approach 0. Under these conditions, and when integrating out the other model parameters, both the WebPPL and Birch simulations scripts for BAMM produce the same normalization constants as the corresponding LSBDS scripts (Supplementary Figure 10),

#### 8 Empirical data

For the empirical analyses illustrating PPL inference for phylogenetic diversification models, we used the bird trees analyzed previously for the ClaDS2 model by Maliet et al.[30]. The trees originate from an earlier study inferring a global timed phylogeny of birds[44]. Specifically, clades with 50 or more leaves (excluding outgroups) from the earlier study were selected in the ClaDS2 study[30] and post-processed to remove outgroups and to rescale branch lengths to time units (myr). Also, only species for which there is molecular data have been included in the trees analyzed by Maliet et al.[30]; consequently the authors calculated a sampling fraction ($\rho$) by dividing the number of tips in the trees computed for species with genetic data by number of tips in the corresponding complete tree (private correspondence).

[h] https://github.com/hoehna/birth-death-shift-analyses

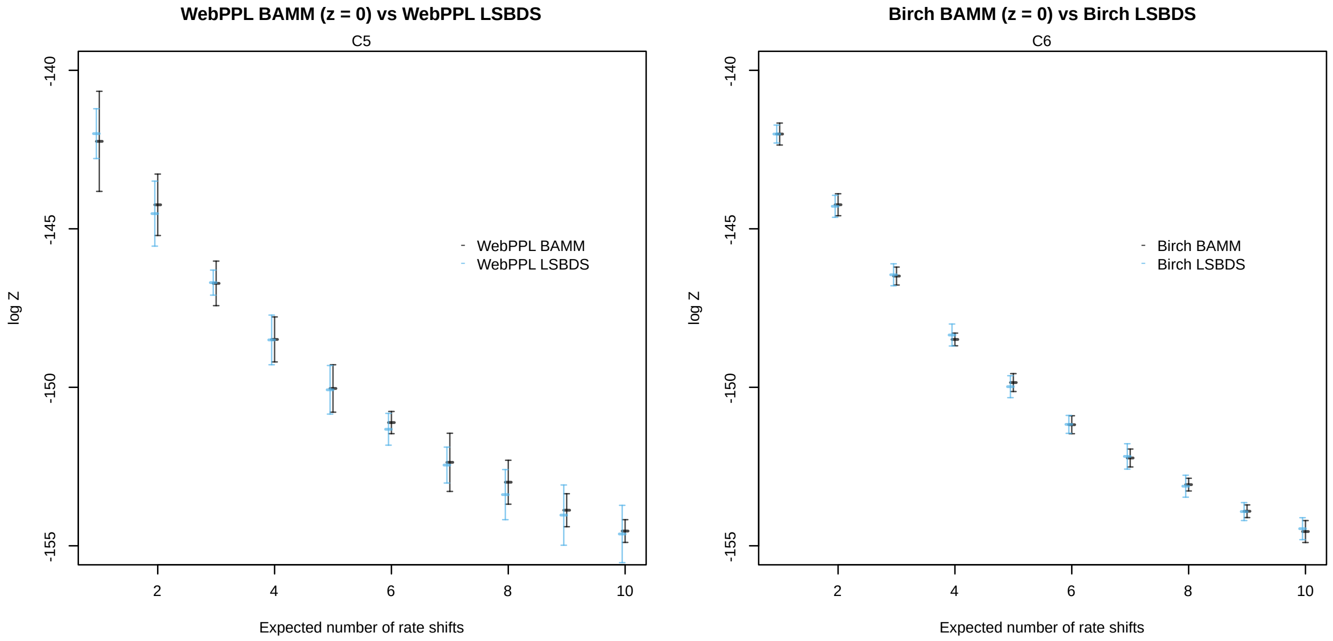


**Fig. 10** Verification that the WebPPL and Birch scripts for the BAMM model generate normalization constant estimates that match those of the corresponding scripts for the LSBDS model for some points in parameter space where the BAMM model collapses to LSBDS.

We downloaded these post-processed trees from the repository[i] accompanying the ClaDS2 paper and extracted the corresponding sampling fractions $\rho$.

The trees were converted from binary R data (RData) to text format (Nexus) with the `ape` package. The Nexus files were then converted to PhyJSON with the `nexus2phyjson` tool that we provide[18]. Next, the PhyJSON trees were reoriented to avoid any systematic left-right imbalances in the original trees that could have a negative effect on inference (see Section 6.4). The resulting PhyJSON trees were then used as input data for the WebPPL and Birch analyses.

There are 42 bird clades in the Jetz et al.[44] study with more than 50 species excluding outgroups. However, we discovered that two of the trees, P2 and Scolopaci, have negative branch lengths. Rather than introducing arbitrary corrections for the negative branch lengths, we excluded these trees from further analysis. The remaining 40 bird trees are summarized in Table 6. The original names of the bird clades[44] are rather cryptic. Here, we named the clades after the family (or other higher taxon) to which most members belong according to the taxonomic classification used in the original bird study[44]. If a family is split between two clades, the clades are numbered 1 and 2. A '-' sign after the family name indicates that some members of the family are not included in the clade; a '+' sign indicates that the clade includes some members of other families. Four of the trees in the repository accompanying the ClaDS paper[30] are mislabeled there: Caprimulgidae is incorrectly labeled CC7, CC4 is labeled Cathartidae, CC7 is labeled CC5CC6B, and CC8 is labeled CC5CC6C.

The size of the trees vary from 54 (Alcedinidae) to 316 leaves (Tyrannidae+), and the ages from 12.5 Ma (Thraupidae1+) to 66.6 Ma (Cuculidae). The fraction of species included in the trees, that is, the sampling fraction $\rho$, varies from 0.43 (Columbidae) to 0.91 (Hirundinidae). The tree shapes are depicted in Supplementary Figures 11, 12.

### 9 Efficiency of inference algorithms

In this section, we provide detailed information about the efficiency of the inference algorithms, taking into account both the quality of the samples of the posterior distribution, and the computational resources needed in obtaining those samples. Specifically, we take advantage of the most obvious approach to measuring the quality of a Monte Carlo inference procedure: we repeat the analysis many times (by running the corresponding program multiple times), and then assess the consistency of the estimates.

We focus on the normalization constant estimates across independent Monte Carlo analyses, as the normalization constant is influenced by all components in the model, including priors, latent variables and data. Thus, the consistency of the normalization constant estimates should provide a good overall estimate of the efficiency of the inference procedure.

Clearly, the more computational resources we invest in obtaining an estimate of the normalization constant, the better that estimate will be. When is the estimate good enough? This depends on how the estimate is to be used. Consider, for example, if the estimate is to be used within a pseudomarginal Metropolis-Hastings sampler. In this case, there is a trade-off between the quality of the normalization constant estimate and the overall efficiency of the MCMC procedure. The efficiency of the MCMC procedure (per time unit) will increase with the precision of the normalization constant estimate, but will decrease with the time required to obtain it. Given some reasonable assumptions, it turns out that the optimal choice is to target a standard deviation of

[i] `https://github.com/OdileMaliet/ClaDS/tree/master/birds_MCC_results`

**Table 6** Overview of the bird trees used for diversification analyses.

| Tree | Clade (Jetz et al.) | Leaves | $\rho$ | Age (Ma) | Notes |
|---|---|---|---|---|---|
| Accipitridae | Accipitridae | 175 | 0.71 | 59.6 | Hawks, eagles, kites and allies |
| Alcedinidae | Alcedinidae | 54 | 0.57 | 34.9 | Kingfishers |
| Anatinae | Anatinae | 108 | 0.87 | 20.3 | Dabbling ducks |
| Caprimulgidae | Caprimulgidae | 57 | 0.61 | 57.3 | Nightjars |
| Campephagidae- | CC4 | 70 | 0.85 | 30.1 | Cuckooshrikes and allies |
| Charadrii | Charadrii | 63 | 0.62 | 59.6 | Waders |
| Columbidae | Columbidae | 133 | 0.43 | 35.9 | Pigeons and doves |
| Corvidae+ | CC8 | 234 | 0.66 | 30.0 | Crows, magpies, monarchs and allies |
| Cuculidae | Cuculidae | 126 | 0.88 | 66.6 | Cuckoos |
| Emberizidae- | P20b | 125 | 0.77 | 14.8 | Buntings |
| Estrildidae | P7 | 101 | 0.62 | 19.5 | Estrildid finches |
| Fringillidae+ | P10 | 123 | 0.63 | 25.4 | True finches |
| Furnaridae | Furnaridae | 205 | 0.67 | 19.9 | Ovenbirds |
| Hirundinidae | S6 | 77 | 0.91 | 23.1 | Swallows, martins and allies |
| Icteridae | P21 | 92 | 0.88 | 14.0 | New World blackbirds, orioles and allies |
| Lari | Lari | 127 | 0.84 | 24.6 | Gulls |
| Malaconotidae+ | CC7 | 80 | 0.55 | 31.4 | Bushshrikes |
| Meliphagidae-+ | BC7 | 90 | 0.49 | 37.1 | Honeyeaters |
| Muscicapidae-+ | M6 | 231 | 0.77 | 20.2 | Old World flycatchers |
| Paridae+ | S2 | 55 | 0.72 | 40.9 | Tits |
| Parulidae+ | P20a | 111 | 0.89 | 17.2 | New World warblers |
| Phasianidae | Phasianidae | 131 | 0.73 | 27.2 | Pheasants, partridges and allies |
| Picidae | Picidae | 137 | 0.61 | 27.1 | Woodpeckers |
| Procellariidae | Procellariidae | 105 | 0.81 | 59.6 | Shearwaters, fulmarine petrels and allies |
| Psittacidae1 | Psittacidae1 | 111 | 0.65 | 33.2 | True parrots (part) |
| Psittacidae2 | Psittacidae2 | 118 | 0.72 | 34.9 | True parrots (part) |
| Pycnonotidae-+ | S9 | 95 | 0.73 | 29.4 | Bulbuls |
| Ramphastidae | Ramphastidae | 81 | 0.65 | 32.2 | Toucans |
| Strigidae | Strigidae | 101 | 0.52 | 45.7 | True owls |
| Sturnidae+ | M4 | 130 | 0.87 | 24.9 | Starlings, mockingbirds and allies |
| Syvliidae1+ | S11 | 79 | 0.65 | 28.1 | Warblers, parrotbills and allies (part) |
| Sylviidae2+ | S7S8 | 93 | 0.79 | 24.3 | Warblers, parrotbills and allies (part) |
| Thamnophilidae | Thamnophilidae | 165 | 0.74 | 22.4 | Antbirds |
| Thraupidae1+ | P13P14P16 | 158 | 0.71 | 12.5 | Tanagers (part) |
| Thraupidae2+ | P17P18 | 139 | 0.89 | 13.7 | Tanagers (part) |
| Timaliidae-+ | S13 | 180 | 0.49 | 21.0 | Old World babblers |
| Trochilidae | Trochilidae | 233 | 0.69 | 28.1 | Hummingbirds |
| Troglodytidae+ | M1 | 91 | 0.69 | 32.7 | Wrens |
| Turdidae-+ | M5 | 134 | 0.86 | 21.7 | Thrushes |
| Tyrannidae+ | Tittyranrest | 316 | 0.69 | 33.6 | Tyrant flycatchers |

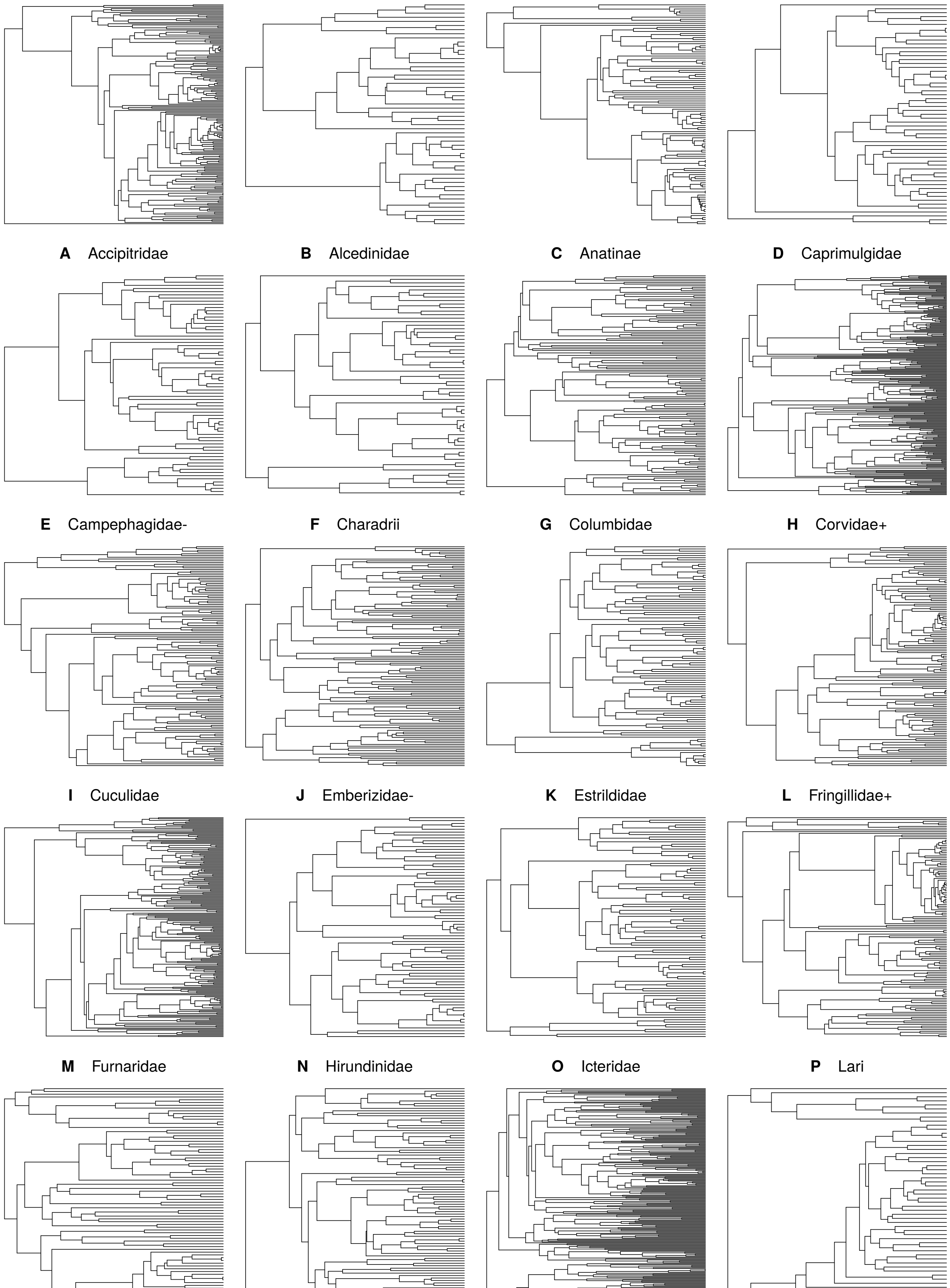


**Fig. 11** Shape of the bird trees, part 1

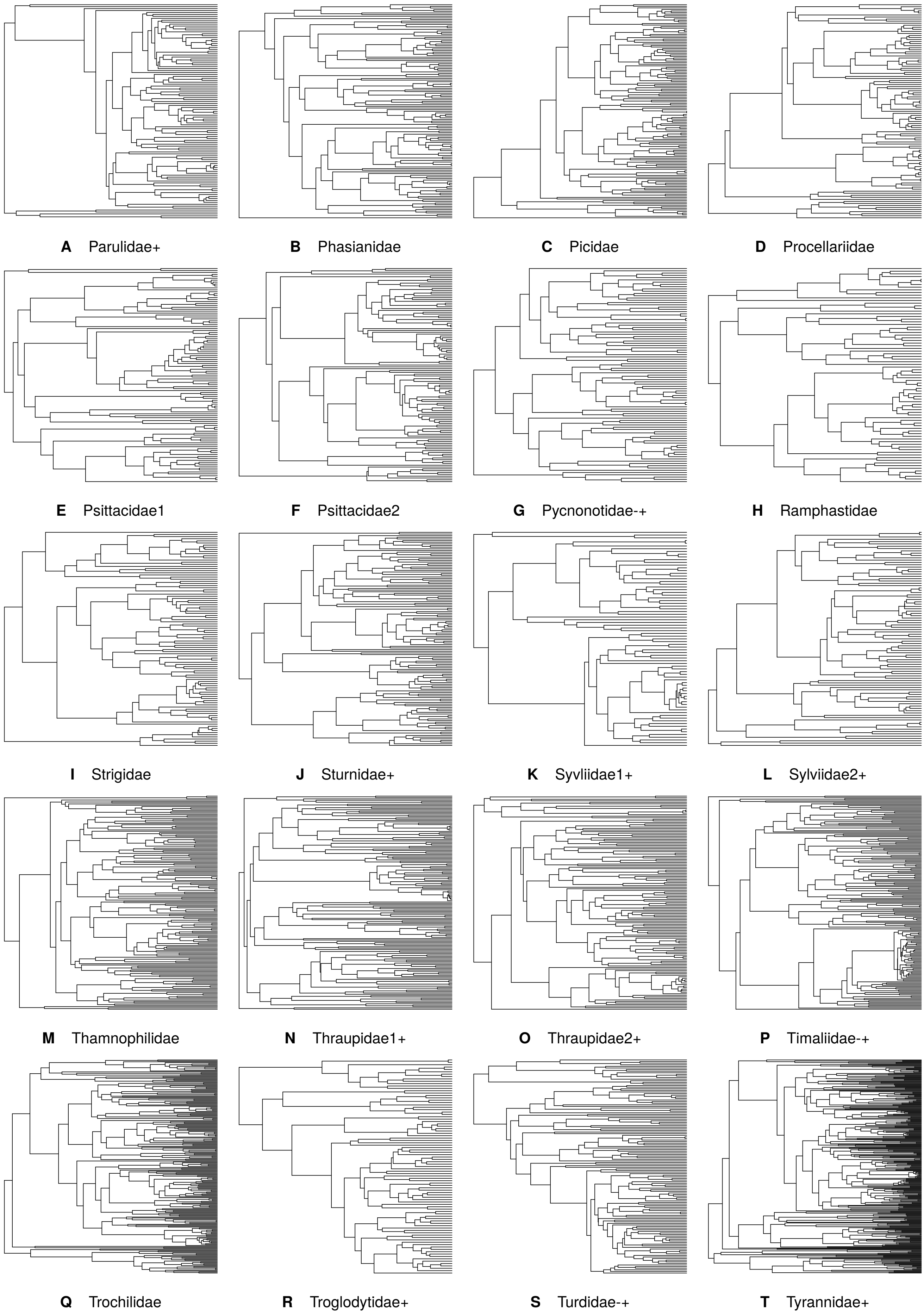


**Fig. 12** Shape of the bird trees, part 2

1.0 if the Metropolis-Hastings algorithm using the exact likelihood is efficient, and around 1.7 when it is not[45]. This suggests that a normalization constant estimate with a standard deviation close to 1.0 is more than satisfactory for this kind of demanding application.

With this in mind, we use two additional diagnostics to assess the efficiency of our inference procedure: the relative effective sample size (RESS) and the conditional acceptance rate (CAR). The former assesses the effect of using the normalizing constant estimates as the weights for an importance sampler, the latter of using them as unbiased estimates of the marginal likelihood for a pseudomarginal sampler.

Specifically, if we have $N$ particles with normalized weights $\{w_1, w_2, \ldots, w_N\}$, the effective sample size $N_{\text{eff}}$ is computed as

$$N_{\text{eff}} = \frac{(\sum_i w_i)^2}{\sum_i w_i^2}. \tag{14}$$

This is known as Kish's ESS[46]. We then compute the relative ESS (RESS) simply by dividing the ESS with the number of independent Monte Carlo estimates we had of the normalization constant. The RESS, which is a value on the interval $(0, 1]$, measures the efficiency of the Monte Carlo estimation procedure[40].

Like RESS, CAR is a value on the interval $(0, 1]$[47]. It measures the expected drop in acceptance rate of a Metropolis-Hastings proposal due to errors in the estimate of the normalization constant, that is, similar to the logic we described above in establishing a quality criterion for the standard deviation. This theoretical acceptance rate is measured for each sampled point, as though the proposal distribution were shrunk to a Dirac $\delta$ distribution. The CAR is related to the standard deviation of the normalization constant estimate, but, like RESS, is a more direct measure of the impacts of that estimate on an inference procedure.

In the initial analyses, we used 10,000 particles in the importance sampling procedure (for the CRB(D) and TDB(D) models) and 5,000 particles in the SMC procedure with the alive particle filter (for the remaining models). The standard deviation of the normalization constant was well below 1.0 in all sequential importance sampling runs (Table 7). In the SMC runs, the standard deviation was usually close to or below 1.0 for all models except BAMM. However, we noted standard deviations above 2.0 in 2 of 40 trees for ClaDS1 (Muscicapidae-+ and Thamnophilidae) and in 5 of 40 trees for LSBDS (Accipitridae, Muscicapidae-+, Timaliidae-+, Tyrannidae+ and Trochilidae). For BAMM, we increased the number of particles to 20,000, which brought the standard deviations down to more acceptable levels, even if we still noted values above 2.0 for 17 of 40 trees.

The RESS and CAR values largely reflect the standard deviation of the normalization constant estimates. However, we observed some SMC cases where the RESS and CAR values suggest that the sample is reasonably good, even though the standard deviation is high. Notable examples include the BAMM results for Corvidae+, Columbidae and Cuculidae, and the LSBDS results for Muscicapidae-+ and Trochilidae.

All analyses of empirical data were run on Tetralith, the largest high-performance computing cluster at the National Supercomputer Centre (NSC), Sweden, a part of the Swedish National Infrastructure for Computing (SNIC). The cluster comprises 1908 nodes, each with two Intel Xeon Gold 6130 CPUs (each with 16 cores). There are 1832 nodes with 96 GiB and 60 nodes with 384 GiB of RAM.[j] The operating system on the nodes is Linux (version 3.10.0) and the cluster uses SLURM (18.08.8) to schedule jobs. For each tree and model, we submitted an array of 10 single-core jobs with the memory limit set to 8 GiB (to avoid running out of memory for the largest trees), each running the respective Birch program 50 times. We used the latest development version of Birch[k] and its standard library (as of June 12, 2020).

The median time (among 500 replicates) required to complete an importance sampling analysis (10,000 particles) using this hardware and analysis setup ranged from a few seconds to around one minute (Table 8). The SMC analyses (5,000 or 20,000 particles) were more demanding, requiring from around a minute to more than 50 minutes in extreme cases. The longest run times were usually associated with the BAMM model, for which we used four times as many particles (20,000) as for the other models. For models other than BAMM, the median run times rarely exceeded ten minutes. As one might expect, the largest trees were generally associated with the longest run times.

### 10 Extended results

The main purpose of the empirical analyses is to demonstrate the power of probabilistic programming in addressing inference problems in phylogenetics, not to advance the field of diversification studies. Nevertheless, there are several interesting patterns in the results that deserve attention and that may inspire further study. In this section, we present model likelihoods and posterior estimates of model parameters for all bird clades and diversification models (Supplementary Figures 13–22). The plots of posterior distributions should be interpreted in relation to the prior distributions for the corresponding regions of parameter space (Supplementary Figures 23). We structure the discussion of the results around several cross-cutting themes.

#### 10.1 Conservative nature of Bayesian model tests

One of the most striking patterns across the bird trees, especially given the recent debate about the importance of accommodating lineage-specific diversification rates, is that simple birth-death models do so well in a Bayesian model comparison. In at least 16 of the 40 bird trees, there is no strong evidence against the simple CRB and CRBD models. In fact, in most of these cases, the simple

[j] https://www.nsc.liu.se/systems/tetralith/

[k] https://github.com/lawmurray/Birch

**Table 7** Diagnostics for the normalization constant estimates obtained across 500 runs for each tree and model. The first line in each cell shows the mean and standard deviation of the normalization constant estimates. We also give the relative effective sample size (RESS, the second line) and conditional acceptance ratio (CAR, the third line).

| Tree | CRB | CRBD | TDB | TDBD | ClaDS0 | ClaDS1 | ClaDS2 | LSBDS | BAMM |
|---|---|---|---|---|---|---|---|---|---|
| Accipitridae | −1219.1 ± 0.1<br>RESS: 0.995<br>CAR: 0.962 | −1218.9 ± 0.1<br>RESS: 0.994<br>CAR: 0.956 | −1220.6 ± 0.2<br>RESS: 0.970<br>CAR: 0.901 | −1220.1 ± 0.1<br>RESS: 0.987<br>CAR: 0.935 | −1196.7 ± 0.7<br>RESS: 0.656<br>CAR: 0.630 | −1196.9 ± 1.1<br>RESS: 0.042<br>CAR: 0.331 | −1196.7 ± 1.0<br>RESS: 0.442<br>CAR: 0.500 | −1214.1 ± 3.6<br>RESS: 0.034<br>CAR: 0.092 | −1212.3 ± 3.0<br>RESS: 0.055<br>CAR: 0.119 |
| Alcedinidae | −304.4 ± 0.0<br>RESS: 0.998<br>CAR: 0.974 | −305.5 ± 0.1<br>RESS: 0.997<br>CAR: 0.967 | −305.6 ± 0.1<br>RESS: 0.995<br>CAR: 0.960 | −306.0 ± 0.1<br>RESS: 0.996<br>CAR: 0.965 | −306.9 ± 0.2<br>RESS: 0.940<br>CAR: 0.861 | −308.9 ± 0.7<br>RESS: 0.112<br>CAR: 0.564 | −307.7 ± 0.6<br>RESS: 0.521<br>CAR: 0.666 | −307.5 ± 0.4<br>RESS: 0.857<br>CAR: 0.781 | −308.6 ± 0.6<br>RESS: 0.831<br>CAR: 0.784 |
| Anatinae | −587.8 ± 0.0<br>RESS: 0.999<br>CAR: 0.979 | −586.2 ± 0.0<br>RESS: 0.998<br>CAR: 0.977 | −587.8 ± 0.1<br>RESS: 0.995<br>CAR: 0.962 | −586.7 ± 0.0<br>RESS: 0.998<br>CAR: 0.972 | −576.1 ± 0.7<br>RESS: 0.547<br>CAR: 0.599 | −576.6 ± 1.0<br>RESS: 0.041<br>CAR: 0.365 | −575.9 ± 0.9<br>RESS: 0.376<br>CAR: 0.511 | −581.0 ± 1.2<br>RESS: 0.236<br>CAR: 0.393 | −579.8 ± 1.2<br>RESS: 0.053<br>CAR: 0.292 |
| Caprimulgidae | −347.8 ± 0.1<br>RESS: 0.997<br>CAR: 0.970 | −349.1 ± 0.1<br>RESS: 0.994<br>CAR: 0.955 | −349.4 ± 0.1<br>RESS: 0.990<br>CAR: 0.942 | −350.1 ± 0.1<br>RESS: 0.992<br>CAR: 0.949 | −348.1 ± 0.5<br>RESS: 0.775<br>CAR: 0.735 | −348.7 ± 0.7<br>RESS: 0.475<br>CAR: 0.580 | −348.5 ± 0.6<br>RESS: 0.617<br>CAR: 0.638 | −351.5 ± 1.7<br>RESS: 0.084<br>CAR: 0.399 | −352.3 ± 3.2<br>RESS: 0.004<br>CAR: 0.030 |
| Campephagidae- | −395.7 ± 0.0<br>RESS: 0.998<br>CAR: 0.975 | −397.0 ± 0.1<br>RESS: 0.996<br>CAR: 0.962 | −396.4 ± 0.1<br>RESS: 0.997<br>CAR: 0.967 | −396.9 ± 0.1<br>RESS: 0.997<br>CAR: 0.968 | −397.0 ± 0.2<br>RESS: 0.944<br>CAR: 0.863 | −398.7 ± 0.5<br>RESS: 0.219<br>CAR: 0.667 | −397.9 ± 0.4<br>RESS: 0.869<br>CAR: 0.794 | −399.0 ± 0.3<br>RESS: 0.903<br>CAR: 0.821 | −399.7 ± 0.3<br>RESS: 0.888<br>CAR: 0.815 |
| Charadrii | −400.3 ± 0.1<br>RESS: 0.996<br>CAR: 0.966 | −399.9 ± 0.1<br>RESS: 0.996<br>CAR: 0.966 | −401.9 ± 0.1<br>RESS: 0.983<br>CAR: 0.925 | −400.2 ± 0.1<br>RESS: 0.995<br>CAR: 0.961 | −404.3 ± 0.3<br>RESS: 0.895<br>CAR: 0.812 | −404.5 ± 0.7<br>RESS: 0.601<br>CAR: 0.632 | −402.4 ± 0.7<br>RESS: 0.541<br>CAR: 0.615 | −402.2 ± 0.4<br>RESS: 0.833<br>CAR: 0.770 | −403.9 ± 0.9<br>RESS: 0.353<br>CAR: 0.532 |
| Columbidae | −889.0 ± 0.1<br>RESS: 0.997<br>CAR: 0.969 | −890.7 ± 0.1<br>RESS: 0.987<br>CAR: 0.936 | −889.4 ± 0.1<br>RESS: 0.989<br>CAR: 0.940 | −888.9 ± 0.1<br>RESS: 0.992<br>CAR: 0.951 | −887.4 ± 0.7<br>RESS: 0.606<br>CAR: 0.603 | −890.4 ± 1.7<br>RESS: 0.186<br>CAR: 0.307 | −888.4 ± 1.2<br>RESS: 0.412<br>CAR: 0.459 | −894.1 ± 0.9<br>RESS: 0.589<br>CAR: 0.581 | −894.0 ± 4.1<br>RESS: 0.185<br>CAR: 0.389 |
| Corvidae+ | −1594.3 ± 0.1<br>RESS: 0.996<br>CAR: 0.966 | −1596.8 ± 0.1<br>RESS: 0.981<br>CAR: 0.922 | −1595.1 ± 0.1<br>RESS: 0.989<br>CAR: 0.939 | −1596.5 ± 0.1<br>RESS: 0.980<br>CAR: 0.919 | −1586.0 ± 0.8<br>RESS: 0.499<br>CAR: 0.550 | −1589.0 ± 1.9<br>RESS: 0.148<br>CAR: 0.340 | −1587.9 ± 1.3<br>RESS: 0.386<br>CAR: 0.431 | −1600.0 ± 1.0<br>RESS: 0.468<br>CAR: 0.507 | −1600.7 ± 3.2<br>RESS: 0.276<br>CAR: 0.408 |
| Cuculidae | −881.5 ± 0.1<br>RESS: 0.994<br>CAR: 0.957 | −883.2 ± 0.1<br>RESS: 0.982<br>CAR: 0.923 | −883.1 ± 0.2<br>RESS: 0.976<br>CAR: 0.912 | −883.2 ± 0.1<br>RESS: 0.987<br>CAR: 0.937 | −882.1 ± 0.4<br>RESS: 0.855<br>CAR: 0.776 | −884.0 ± 0.6<br>RESS: 0.698<br>CAR: 0.676 | −882.7 ± 0.5<br>RESS: 0.794<br>CAR: 0.729 | −886.1 ± 0.6<br>RESS: 0.705<br>CAR: 0.671 | −889.0 ± 2.8<br>RESS: 0.129<br>CAR: 0.277 |
| Emberizidae- | −737.9 ± 0.0<br>RESS: 0.998<br>CAR: 0.974 | −740.7 ± 0.1<br>RESS: 0.985<br>CAR: 0.930 | −725.7 ± 0.2<br>RESS: 0.969<br>CAR: 0.899 | −727.5 ± 0.4<br>RESS: 0.870<br>CAR: 0.783 | −731.2 ± 0.4<br>RESS: 0.831<br>CAR: 0.756 | −735.4 ± 1.2<br>RESS: 0.508<br>CAR: 0.556 | −732.7 ± 0.5<br>RESS: 0.715<br>CAR: 0.700 | −744.2 ± 1.2<br>RESS: 0.428<br>CAR: 0.472 | −733.7 ± 2.3<br>RESS: 0.080<br>CAR: 0.172 |
| Estrildidae | −567.4 ± 0.0<br>RESS: 0.998<br>CAR: 0.977 | −569.0 ± 0.1<br>RESS: 0.995<br>CAR: 0.961 | −567.6 ± 0.1<br>RESS: 0.998<br>CAR: 0.972 | −568.4 ± 0.1<br>RESS: 0.996<br>CAR: 0.966 | −569.5 ± 0.7<br>RESS: 0.639<br>CAR: 0.629 | −571.2 ± 1.1<br>RESS: 0.384<br>CAR: 0.452 | −570.0 ± 1.1<br>RESS: 0.394<br>CAR: 0.464 | −570.9 ± 1.0<br>RESS: 0.320<br>CAR: 0.471 | −569.3 ± 1.5<br>RESS: 0.064<br>CAR: 0.199 |
| Fringillidae+ | −727.7 ± 0.0<br>RESS: 0.998<br>CAR: 0.974 | −728.8 ± 0.1<br>RESS: 0.996<br>CAR: 0.962 | −728.8 ± 0.1<br>RESS: 0.995<br>CAR: 0.960 | −729.6 ± 0.1<br>RESS: 0.994<br>CAR: 0.958 | −718.2 ± 0.6<br>RESS: 0.706<br>CAR: 0.697 | −720.3 ± 0.6<br>RESS: 0.674<br>CAR: 0.657 | −719.3 ± 0.7<br>RESS: 0.659<br>CAR: 0.639 | −726.4 ± 0.9<br>RESS: 0.439<br>CAR: 0.533 | −726.3 ± 0.9<br>RESS: 0.377<br>CAR: 0.495 |
| Furnaridae | −1262.7 ± 0.0<br>RESS: 0.998<br>CAR: 0.974 | −1265.0 ± 0.1<br>RESS: 0.989<br>CAR: 0.941 | −1263.0 ± 0.1<br>RESS: 0.996<br>CAR: 0.965 | −1264.7 ± 0.1<br>RESS: 0.988<br>CAR: 0.938 | −1253.5 ± 0.8<br>RESS: 0.537<br>CAR: 0.594 | −1256.0 ± 1.0<br>RESS: 0.456<br>CAR: 0.520 | −1255.1 ± 0.9<br>RESS: 0.494<br>CAR: 0.547 | −1264.1 ± 1.0<br>RESS: 0.426<br>CAR: 0.509 | −1262.8 ± 0.9<br>RESS: 0.487<br>CAR: 0.527 |
| Hirundinidae | −433.1 ± 0.0<br>RESS: 0.998<br>CAR: 0.974 | −434.9 ± 0.1<br>RESS: 0.993<br>CAR: 0.952 | −432.1 ± 0.1<br>RESS: 0.997<br>CAR: 0.970 | −433.0 ± 0.1<br>RESS: 0.995<br>CAR: 0.961 | −433.5 ± 0.3<br>RESS: 0.909<br>CAR: 0.826 | −435.6 ± 0.6<br>RESS: 0.740<br>CAR: 0.684 | −434.2 ± 0.5<br>RESS: 0.735<br>CAR: 0.701 | −437.1 ± 0.5<br>RESS: 0.798<br>CAR: 0.732 | −436.1 ± 0.8<br>RESS: 0.078<br>CAR: 0.454 |
| Icteridae | −495.6 ± 0.0<br>RESS: 0.999<br>CAR: 0.979 | −497.9 ± 0.1<br>RESS: 0.992<br>CAR: 0.948 | −492.6 ± 0.1<br>RESS: 0.995<br>CAR: 0.962 | −494.2 ± 0.1<br>RESS: 0.987<br>CAR: 0.936 | −494.6 ± 0.4<br>RESS: 0.863<br>CAR: 0.787 | −497.4 ± 0.6<br>RESS: 0.714<br>CAR: 0.683 | −495.8 ± 0.5<br>RESS: 0.808<br>CAR: 0.744 | −500.2 ± 0.5<br>RESS: 0.816<br>CAR: 0.738 | −497.6 ± 0.6<br>RESS: 0.664<br>CAR: 0.652 |
| Lari | −743.9 ± 0.0<br>RESS: 0.998<br>CAR: 0.978 | −738.2 ± 0.0<br>RESS: 0.998<br>CAR: 0.972 | −742.2 ± 0.1<br>RESS: 0.989<br>CAR: 0.942 | −738.4 ± 0.1<br>RESS: 0.996<br>CAR: 0.964 | −710.9 ± 0.5<br>RESS: 0.783<br>CAR: 0.717 | −712.2 ± 1.0<br>RESS: 0.482<br>CAR: 0.538 | −711.9 ± 0.9<br>RESS: 0.558<br>CAR: 0.570 | −718.7 ± 1.1<br>RESS: 0.502<br>CAR: 0.520 | −718.6 ± 1.0<br>RESS: 0.536<br>CAR: 0.556 |
| Malaconotidae+ | −501.2 ± 0.1<br>RESS: 0.997<br>CAR: 0.968 | −503.3 ± 0.1<br>RESS: 0.987<br>CAR: 0.935 | −498.0 ± 0.1<br>RESS: 0.995<br>CAR: 0.959 | −497.8 ± 0.1<br>RESS: 0.994<br>CAR: 0.957 | −494.9 ± 0.5<br>RESS: 0.797<br>CAR: 0.736 | −498.2 ± 1.1<br>RESS: 0.438<br>CAR: 0.499 | −495.7 ± 0.7<br>RESS: 0.732<br>CAR: 0.685 | −506.4 ± 0.9<br>RESS: 0.458<br>CAR: 0.554 | −503.6 ± 2.6<br>RESS: 0.056<br>CAR: 0.213 |
| Meliphagidae-+ | −570.6 ± 0.1<br>RESS: 0.997<br>CAR: 0.968 | −572.8 ± 0.1<br>RESS: 0.986<br>CAR: 0.932 | −568.6 ± 0.1<br>RESS: 0.994<br>CAR: 0.955 | −569.1 ± 0.1<br>RESS: 0.993<br>CAR: 0.954 | −570.4 ± 0.5<br>RESS: 0.773<br>CAR: 0.723 | −573.0 ± 1.1<br>RESS: 0.249<br>CAR: 0.423 | −571.2 ± 0.9<br>RESS: 0.413<br>CAR: 0.535 | −575.9 ± 1.0<br>RESS: 0.530<br>CAR: 0.598 | −574.4 ± 1.5<br>RESS: 0.084<br>CAR: 0.296 |
| Muscicapidae-+ | −1576.7 ± 0.1<br>RESS: 0.996<br>CAR: 0.964 | −1580.3 ± 0.3<br>RESS: 0.942<br>CAR: 0.860 | −1541.9 ± 0.2<br>RESS: 0.944<br>CAR: 0.864 | −1543.4 ± 0.6<br>RESS: 0.823<br>CAR: 0.736 | −1548.3 ± 0.9<br>RESS: 0.390<br>CAR: 0.501 | −1553.9 ± 2.4<br>RESS: 0.051<br>CAR: 0.178 | −1549.7 ± 1.4<br>RESS: 0.267<br>CAR: 0.370 | −1586.3 ± 2.9<br>RESS: 0.147<br>CAR: 0.209 | −1557.3 ± 8.1<br>RESS: 0.016<br>CAR: 0.043 |
| Paridae+ | −327.2 ± 0.0<br>RESS: 0.998<br>CAR: 0.972 | −328.8 ± 0.1<br>RESS: 0.993<br>CAR: 0.953 | −327.8 ± 0.1<br>RESS: 0.994<br>CAR: 0.956 | −327.8 ± 0.1<br>RESS: 0.996<br>CAR: 0.964 | −319.1 ± 0.6<br>RESS: 0.536<br>CAR: 0.649 | −321.5 ± 0.8<br>RESS: 0.534<br>CAR: 0.579 | −320.0 ± 0.7<br>RESS: 0.626<br>CAR: 0.637 | −331.0 ± 1.0<br>RESS: 0.437<br>CAR: 0.608 | −327.0 ± 3.2<br>RESS: 0.038<br>CAR: 0.102 |

Table 7: (continued)

| Tree | | CRB | CRBD | TDB | TDBD | ClaDS0 | ClaDS1 | ClaDS2 | LSBDS | BAMM |
|---|---|---|---|---|---|---|---|---|---|---|
| Parulidae+ | | −620.5 ± 0.0 | −622.7 ± 0.1 | −619.7 ± 0.1 | −621.3 ± 0.1 | −600.3 ± 0.9 | −603.1 ± 1.0 | −601.2 ± 1.0 | −622.6 ± 1.6 | −615.1 ± 2.8 |
| | RESS: | 0.998 | 0.992 | 0.997 | 0.993 | 0.468 | 0.421 | 0.320 | 0.227 | 0.030 |
| | CAR: | 0.978 | 0.950 | 0.969 | 0.953 | 0.539 | 0.498 | 0.476 | 0.362 | 0.073 |
| Phasianidae | | −808.7 ± 0.0 | −809.5 ± 0.1 | −809.9 ± 0.1 | −809.5 ± 0.1 | −810.7 ± 0.5 | −811.0 ± 0.7 | −810.4 ± 0.8 | −812.0 ± 0.7 | −812.9 ± 1.1 |
| | RESS: | 0.998 | 0.996 | 0.993 | 0.996 | 0.813 | 0.558 | 0.460 | 0.573 | 0.125 |
| | CAR: | 0.972 | 0.963 | 0.954 | 0.964 | 0.746 | 0.611 | 0.564 | 0.613 | 0.400 |
| Picidae | | −830.7 ± 0.0 | −832.8 ± 0.1 | −831.8 ± 0.1 | −833.4 ± 0.1 | −828.5 ± 0.8 | −830.2 ± 1.1 | −830.6 ± 1.5 | −835.4 ± 1.5 | −835.7 ± 2.7 |
| | RESS: | 0.998 | 0.991 | 0.995 | 0.987 | 0.547 | 0.334 | 0.190 | 0.173 | 0.028 |
| | CAR: | 0.974 | 0.945 | 0.959 | 0.934 | 0.597 | 0.444 | 0.339 | 0.291 | 0.068 |
| Procellariidae | | −686.0 ± 0.1 | −684.0 ± 0.1 | −686.5 ± 0.2 | −684.8 ± 0.1 | −680.8 ± 0.7 | −681.9 ± 0.9 | −681.0 ± 0.9 | −685.4 ± 1.0 | −687.1 ± 1.4 |
| | RESS: | 0.997 | 0.996 | 0.972 | 0.993 | 0.638 | 0.444 | 0.486 | 0.324 | 0.058 |
| | CAR: | 0.967 | 0.965 | 0.905 | 0.954 | 0.628 | 0.559 | 0.535 | 0.447 | 0.299 |
| Psittacidae1 | | −690.7 ± 0.1 | −688.8 ± 0.1 | −690.9 ± 0.1 | −689.2 ± 0.1 | −691.9 ± 0.6 | −692.5 ± 1.0 | −691.5 ± 0.9 | −691.4 ± 0.7 | −693.0 ± 2.0 |
| | RESS: | 0.997 | 0.997 | 0.987 | 0.995 | 0.713 | 0.426 | 0.339 | 0.636 | 0.232 |
| | CAR: | 0.972 | 0.969 | 0.934 | 0.962 | 0.685 | 0.511 | 0.493 | 0.633 | 0.419 |
| Psittacidae2 | | −729.5 ± 0.0 | −730.3 ± 0.1 | −730.7 ± 0.1 | −731.3 ± 0.1 | −726.6 ± 0.5 | −727.8 ± 0.7 | −727.3 ± 0.7 | −731.5 ± 0.8 | −732.3 ± 0.7 |
| | RESS: | 0.998 | 0.995 | 0.990 | 0.992 | 0.772 | 0.194 | 0.529 | 0.436 | 0.595 |
| | CAR: | 0.972 | 0.962 | 0.944 | 0.950 | 0.715 | 0.561 | 0.585 | 0.540 | 0.643 |
| Pycnonotidae-+ | | −589.9 ± 0.1 | −592.3 ± 0.1 | −584.0 ± 0.1 | −584.9 ± 0.1 | −585.9 ± 0.5 | −588.9 ± 0.9 | −586.9 ± 0.7 | −595.1 ± 0.6 | −589.8 ± 1.9 |
| | RESS: | 0.997 | 0.983 | 0.995 | 0.991 | 0.803 | 0.457 | 0.638 | 0.772 | 0.482 |
| | CAR: | 0.970 | 0.926 | 0.958 | 0.947 | 0.740 | 0.545 | 0.626 | 0.706 | 0.542 |
| Ramphastidae | | −475.0 ± 0.0 | −475.0 ± 0.1 | −475.9 ± 0.1 | −475.7 ± 0.1 | −476.8 ± 0.5 | −478.3 ± 0.8 | −477.4 ± 0.7 | −477.0 ± 0.5 | −478.2 ± 0.6 |
| | RESS: | 0.998 | 0.998 | 0.993 | 0.996 | 0.795 | 0.480 | 0.493 | 0.811 | 0.483 |
| | CAR: | 0.974 | 0.972 | 0.952 | 0.966 | 0.736 | 0.584 | 0.606 | 0.750 | 0.633 |
| Strigidae | | −645.2 ± 0.1 | −646.4 ± 0.1 | −646.8 ± 0.1 | −647.7 ± 0.1 | −647.5 ± 0.6 | −649.8 ± 1.0 | −649.1 ± 0.9 | −649.1 ± 0.8 | −650.9 ± 1.0 |
| | RESS: | 0.997 | 0.993 | 0.987 | 0.984 | 0.689 | 0.501 | 0.468 | 0.560 | 0.464 |
| | CAR: | 0.968 | 0.954 | 0.935 | 0.928 | 0.677 | 0.531 | 0.517 | 0.589 | 0.558 |
| Sturnidae+ | | −794.2 ± 0.1 | −796.6 ± 0.1 | −792.9 ± 0.1 | −794.2 ± 0.1 | −793.2 ± 0.6 | −795.6 ± 1.0 | −794.5 ± 1.0 | −799.2 ± 0.7 | −797.9 ± 0.9 |
| | RESS: | 0.997 | 0.987 | 0.995 | 0.992 | 0.670 | 0.302 | 0.408 | 0.668 | 0.426 |
| | CAR: | 0.971 | 0.935 | 0.961 | 0.949 | 0.661 | 0.481 | 0.520 | 0.652 | 0.520 |
| Syvliidae1+ | | −453.3 ± 0.0 | −454.1 ± 0.1 | −454.4 ± 0.1 | −454.6 ± 0.1 | −442.9 ± 0.5 | −445.0 ± 0.7 | −444.2 ± 0.7 | −451.0 ± 0.7 | −451.5 ± 0.9 |
| | RESS: | 0.998 | 0.997 | 0.995 | 0.996 | 0.787 | 0.519 | 0.605 | 0.571 | 0.357 |
| | CAR: | 0.975 | 0.971 | 0.962 | 0.966 | 0.721 | 0.620 | 0.616 | 0.612 | 0.569 |
| Sylviidae2+ | | −540.6 ± 0.0 | −542.1 ± 0.1 | −541.3 ± 0.1 | −541.7 ± 0.1 | −537.4 ± 0.6 | −539.4 ± 0.7 | −538.4 ± 0.7 | −544.2 ± 0.6 | −544.2 ± 1.1 |
| | RESS: | 0.998 | 0.994 | 0.997 | 0.996 | 0.698 | 0.634 | 0.603 | 0.659 | 0.195 |
| | CAR: | 0.975 | 0.958 | 0.967 | 0.966 | 0.678 | 0.631 | 0.610 | 0.664 | 0.423 |
| Thamnophilidae | | −1061.2 ± 0.1 | −1064.2 ± 0.2 | −1049.0 ± 0.1 | −1050.5 ± 0.2 | −1046.9 ± 0.9 | −1050.2 ± 2.3 | −1048.6 ± 1.3 | −1067.7 ± 1.1 | −1056.6 ± 2.4 |
| | RESS: | 0.997 | 0.974 | 0.989 | 0.970 | 0.513 | 0.140 | 0.206 | 0.476 | 0.002 |
| | CAR: | 0.970 | 0.909 | 0.940 | 0.901 | 0.542 | 0.306 | 0.365 | 0.511 | 0.004 |
| Thraupidae1+ | | −935.9 ± 0.0 | −938.3 ± 0.1 | −931.3 ± 0.1 | −932.2 ± 0.2 | −919.5 ± 1.0 | −922.8 ± 1.2 | −920.1 ± 1.0 | −935.9 ± 0.8 | −926.0 ± 1.1 |
| | RESS: | 0.998 | 0.989 | 0.993 | 0.977 | 0.309 | 0.180 | 0.406 | 0.559 | 0.264 |
| | CAR: | 0.976 | 0.940 | 0.951 | 0.913 | 0.476 | 0.391 | 0.484 | 0.594 | 0.423 |
| Thraupidae2+ | | −807.6 ± 0.0 | −810.1 ± 0.1 | −801.6 ± 0.1 | −803.1 ± 0.2 | −791.0 ± 0.6 | −794.1 ± 1.0 | −791.7 ± 0.7 | −807.6 ± 1.8 | −798.7 ± 1.6 |
| | RESS: | 0.998 | 0.988 | 0.990 | 0.970 | 0.730 | 0.471 | 0.558 | 0.311 | 0.109 |
| | CAR: | 0.976 | 0.938 | 0.944 | 0.901 | 0.682 | 0.528 | 0.595 | 0.522 | 0.239 |
| Timaliidae-+ | | −1120.2 ± 0.0 | −1121.0 ± 0.1 | −1121.4 ± 0.1 | −1121.4 ± 0.1 | −1082.9 ± 1.5 | −1084.8 ± 1.6 | −1084.3 ± 1.8 | −1096.5 ± 2.2 | −1095.5 ± 3.2 |
| | RESS: | 0.998 | 0.996 | 0.995 | 0.995 | 0.205 | 0.161 | 0.076 | 0.131 | 0.013 |
| | CAR: | 0.974 | 0.963 | 0.960 | 0.958 | 0.332 | 0.295 | 0.206 | 0.207 | 0.030 |
| Trochilidae | | −1567.5 ± 0.1 | −1569.7 ± 0.1 | −1567.9 ± 0.1 | −1569.0 ± 0.1 | −1562.8 ± 0.9 | −1565.4 ± 1.3 | −1563.9 ± 1.2 | −1573.0 ± 3.8 | −1573.0 ± 3.7 |
| | RESS: | 0.997 | 0.983 | 0.992 | 0.986 | 0.483 | 0.363 | 0.366 | 0.273 | 0.097 |
| | CAR: | 0.968 | 0.926 | 0.948 | 0.933 | 0.534 | 0.435 | 0.439 | 0.378 | 0.190 |
| Troglodytidae+ | | −545.7 ± 0.0 | −547.8 ± 0.1 | −546.0 ± 0.1 | −547.4 ± 0.1 | −547.8 ± 0.4 | −550.5 ± 0.7 | −549.3 ± 0.6 | −550.3 ± 0.6 | −550.7 ± 1.0 |
| | RESS: | 0.998 | 0.989 | 0.996 | 0.991 | 0.846 | 0.643 | 0.742 | 0.736 | 0.346 |
| | CAR: | 0.973 | 0.941 | 0.963 | 0.948 | 0.773 | 0.621 | 0.693 | 0.681 | 0.515 |
| Turdidae-+ | | −830.1 ± 0.1 | −833.0 ± 0.2 | −821.3 ± 0.1 | −822.7 ± 0.2 | −813.0 ± 0.9 | −817.4 ± 1.5 | −814.4 ± 1.0 | −836.0 ± 0.8 | −826.4 ± 2.2 |
| | RESS: | 0.997 | 0.975 | 0.992 | 0.978 | 0.508 | 0.299 | 0.461 | 0.578 | 0.014 |
| | CAR: | 0.970 | 0.910 | 0.949 | 0.915 | 0.550 | 0.373 | 0.498 | 0.594 | 0.043 |
| Tyrannidae+ | | −2273.6 ± 0.1 | −2276.0 ± 0.2 | −2274.1 ± 0.1 | −2275.1 ± 0.2 | −2262.0 ± 1.0 | −2262.7 ± 1.6 | −2260.9 ± 1.3 | −2278.2 ± 8.8 | −2281.7 ± 11.4 |
| | RESS: | 0.995 | 0.971 | 0.985 | 0.978 | 0.334 | 0.176 | 0.275 | 0.096 | 0.004 |
| | CAR: | 0.961 | 0.902 | 0.931 | 0.915 | 0.489 | 0.300 | 0.387 | 0.171 | 0.006 |

**Table 8** Median execution time in Birch in seconds for each tree and model. See text for details.

| Tree | CRB | CRBD | TDB | TDBD | ClaDS0 | ClaDS1 | ClaDS2 | LSBDS | BAMM |
|---|---|---|---|---|---|---|---|---|---|
| Accipitridae | 14 | 27 | 30 | 35 | 400 | 460 | 606 | 860 | 2097 |
| Alcedinidae | 4 | 8 | 8 | 11 | 77 | 95 | 214 | 154 | 792 |
| Anatinae | 8 | 17 | 19 | 22 | 222 | 236 | 631 | 226 | 1719 |
| Caprimulgidae | 4 | 9 | 10 | 11 | 96 | 115 | 209 | 289 | 958 |
| Campephagidae- | 5 | 11 | 12 | 14 | 107 | 120 | 221 | 124 | 966 |
| Charadrii | 5 | 11 | 11 | 13 | 109 | 134 | 571 | 328 | 890 |
| Columbidae | 10 | 20 | 23 | 27 | 258 | 295 | 475 | 310 | 1918 |
| Corvidae+ | 18 | 36 | 41 | 47 | 570 | 636 | 820 | 415 | 2852 |
| Cuculidae | 10 | 19 | 22 | 25 | 241 | 271 | 380 | 441 | 1730 |
| Emberizidae- | 10 | 19 | 23 | 25 | 234 | 260 | 450 | 222 | 1669 |
| Estrildidae | 8 | 16 | 17 | 20 | 179 | 213 | 613 | 200 | 1344 |
| Fringillidae+ | 10 | 18 | 21 | 25 | 212 | 233 | 287 | 209 | 1329 |
| Furnaridae | 15 | 30 | 35 | 41 | 463 | 520 | 625 | 243 | 1941 |
| Hirundinidae | 6 | 12 | 13 | 15 | 111 | 129 | 277 | 180 | 942 |
| Icteridae | 7 | 15 | 17 | 20 | 152 | 178 | 334 | 159 | 1077 |
| Lari | 9 | 19 | 22 | 25 | 228 | 238 | 477 | 295 | 1295 |
| Malaconotidae+ | 6 | 12 | 14 | 16 | 131 | 155 | 512 | 813 | 1091 |
| Meliphagidae-+ | 7 | 14 | 16 | 18 | 150 | 173 | 326 | 257 | 1140 |
| Muscicapidae-+ | 18 | 36 | 40 | 47 | 576 | 640 | 909 | 414 | 2941 |
| Paridae+ | 4 | 8 | 9 | 11 | 84 | 98 | 216 | 429 | 677 |
| Parulidae+ | 8 | 18 | 20 | 24 | 210 | 234 | 366 | 257 | 1527 |
| Phasianidae | 10 | 21 | 25 | 26 | 245 | 267 | 412 | 230 | 1348 |
| Picidae | 10 | 21 | 24 | 28 | 269 | 279 | 456 | 295 | 1611 |
| Procellariidae | 8 | 16 | 18 | 20 | 193 | 231 | 386 | 267 | 1351 |
| Psittacidae1 | 8 | 17 | 19 | 22 | 205 | 219 | 477 | 316 | 1237 |
| Psittacidae2 | 9 | 18 | 21 | 22 | 216 | 237 | 360 | 220 | 1425 |
| Pycnonotidae-+ | 7 | 14 | 17 | 19 | 144 | 169 | 243 | 182 | 1261 |
| Ramphastidae | 6 | 13 | 15 | 15 | 122 | 138 | 299 | 233 | 851 |
| Strigidae | 8 | 15 | 18 | 20 | 170 | 204 | 392 | 232 | 1198 |
| Sturnidae+ | 9 | 19 | 21 | 27 | 247 | 251 | 352 | 237 | 1538 |
| Syvliidae1+ | 6 | 10 | 14 | 16 | 114 | 141 | 276 | 279 | 864 |
| Sylviidae2+ | 7 | 14 | 16 | 17 | 152 | 158 | 315 | 287 | 883 |
| Thamnophilidae | 12 | 24 | 28 | 33 | 327 | 418 | 524 | 415 | 1802 |
| Thraupidae1+ | 12 | 27 | 27 | 34 | 314 | 363 | 428 | 224 | 1673 |
| Thraupidae2+ | 11 | 23 | 24 | 30 | 327 | 323 | 723 | 277 | 1753 |
| Timaliidae-+ | 14 | 27 | 31 | 32 | 366 | 411 | 481 | 262 | 1864 |
| Trochilidae | 18 | 33 | 41 | 47 | 578 | 657 | 821 | 432 | 2699 |
| Troglodytidae+ | 7 | 13 | 16 | 18 | 149 | 171 | 289 | 182 | 1193 |
| Turdidae-+ | 10 | 21 | 23 | 27 | 251 | 284 | 426 | 223 | 1447 |
| Tyrannidae+ | 23 | 48 | 55 | 63 | 963 | 1022 | 2073 | 2064 | 3250 |

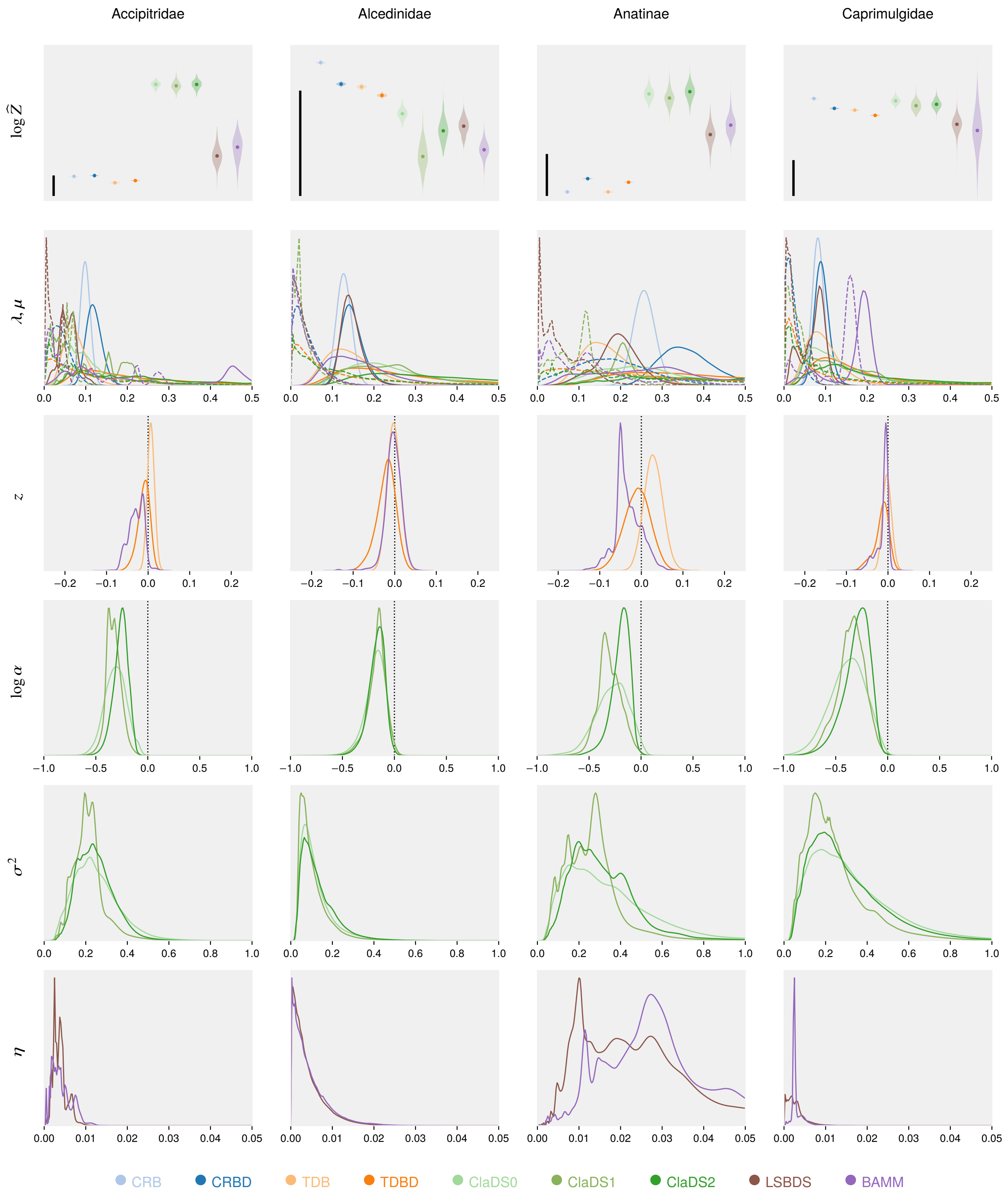


**Fig. 13** Normalization constants and parameter estimates for Accipitridae, Alcedinidae, Anatinae, Caprimulgidae.

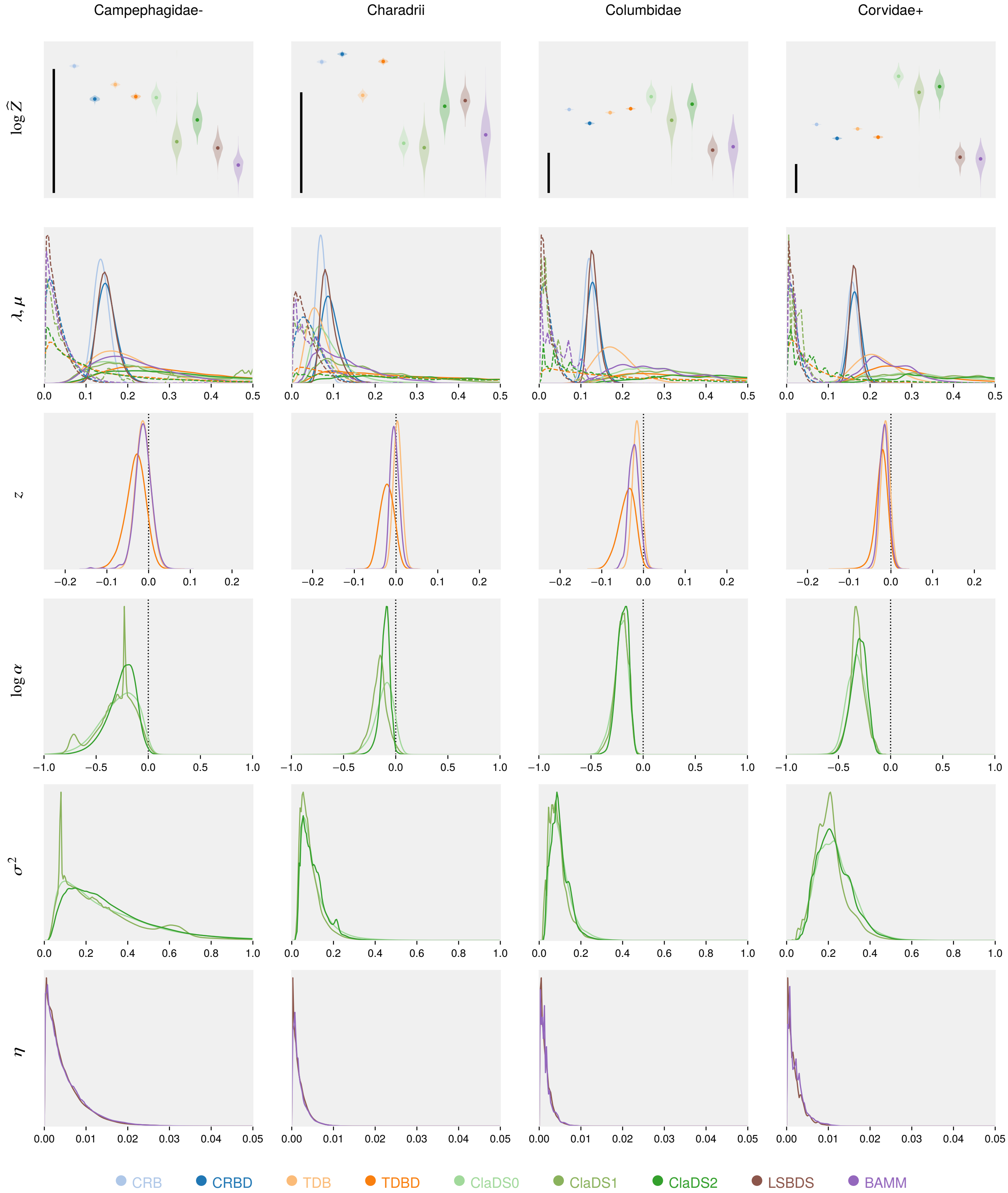


**Fig. 14** Normalization constants and parameter estimates for Campephagidae-, Charadrii, Columbidae, Corvidae+.

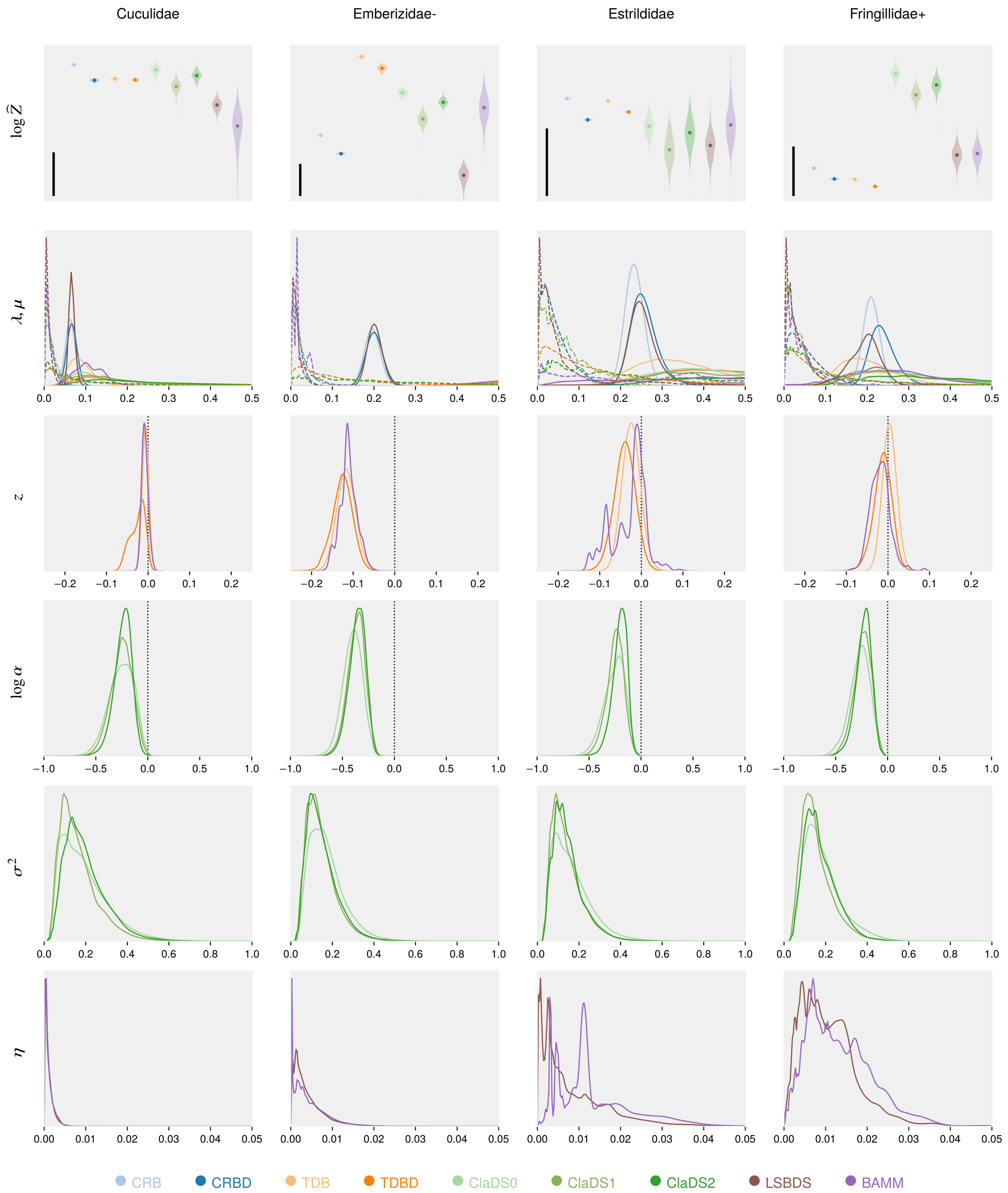


**Fig. 15** Normalization constants and parameter estimates for Cuculidae, Emberizidae-, Estrildidae, Fringillidae+.

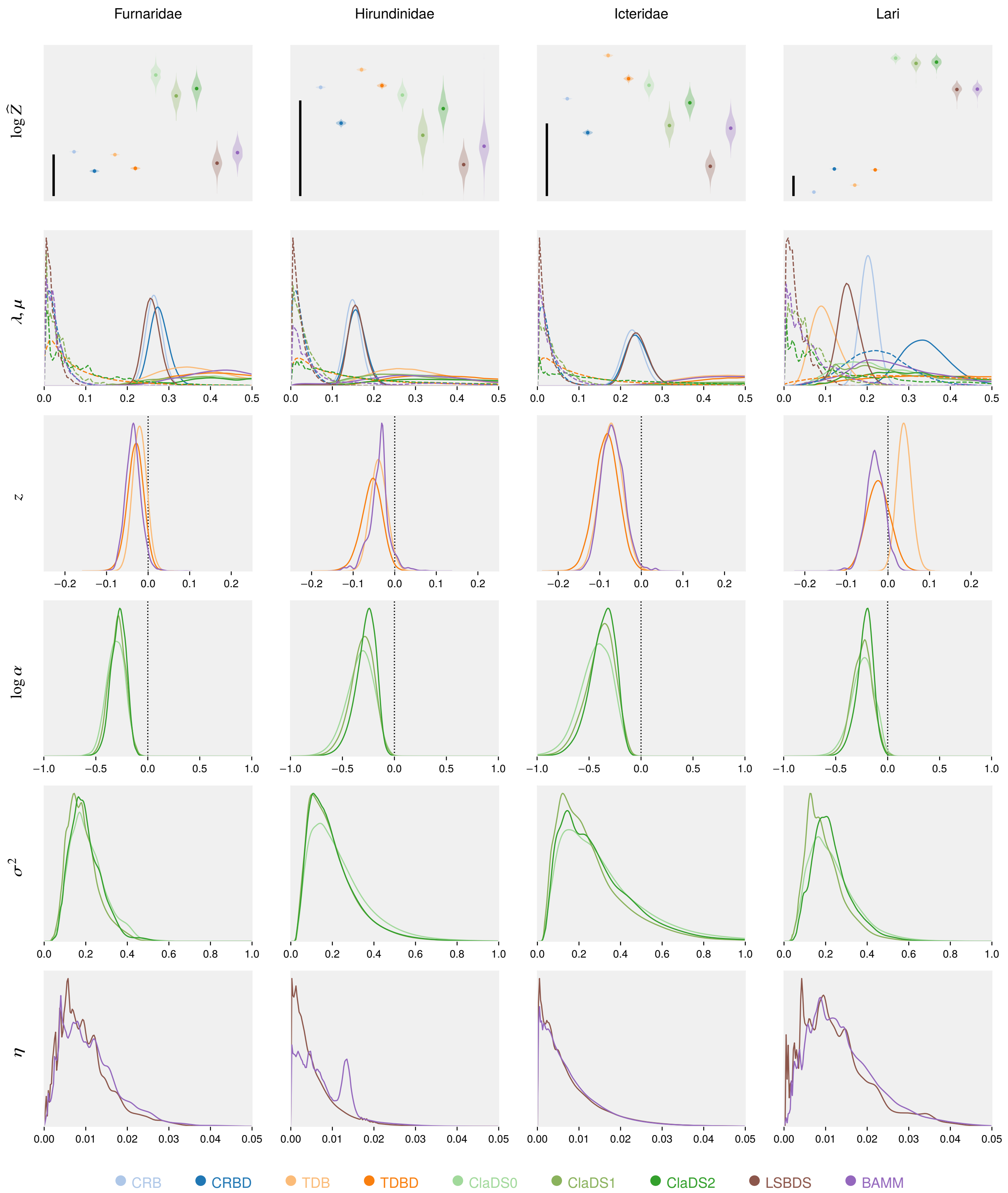


**Fig. 16** Normalization constants and parameter estimates for Furnaridae, Hirundinidae, Icteridae, Lari.

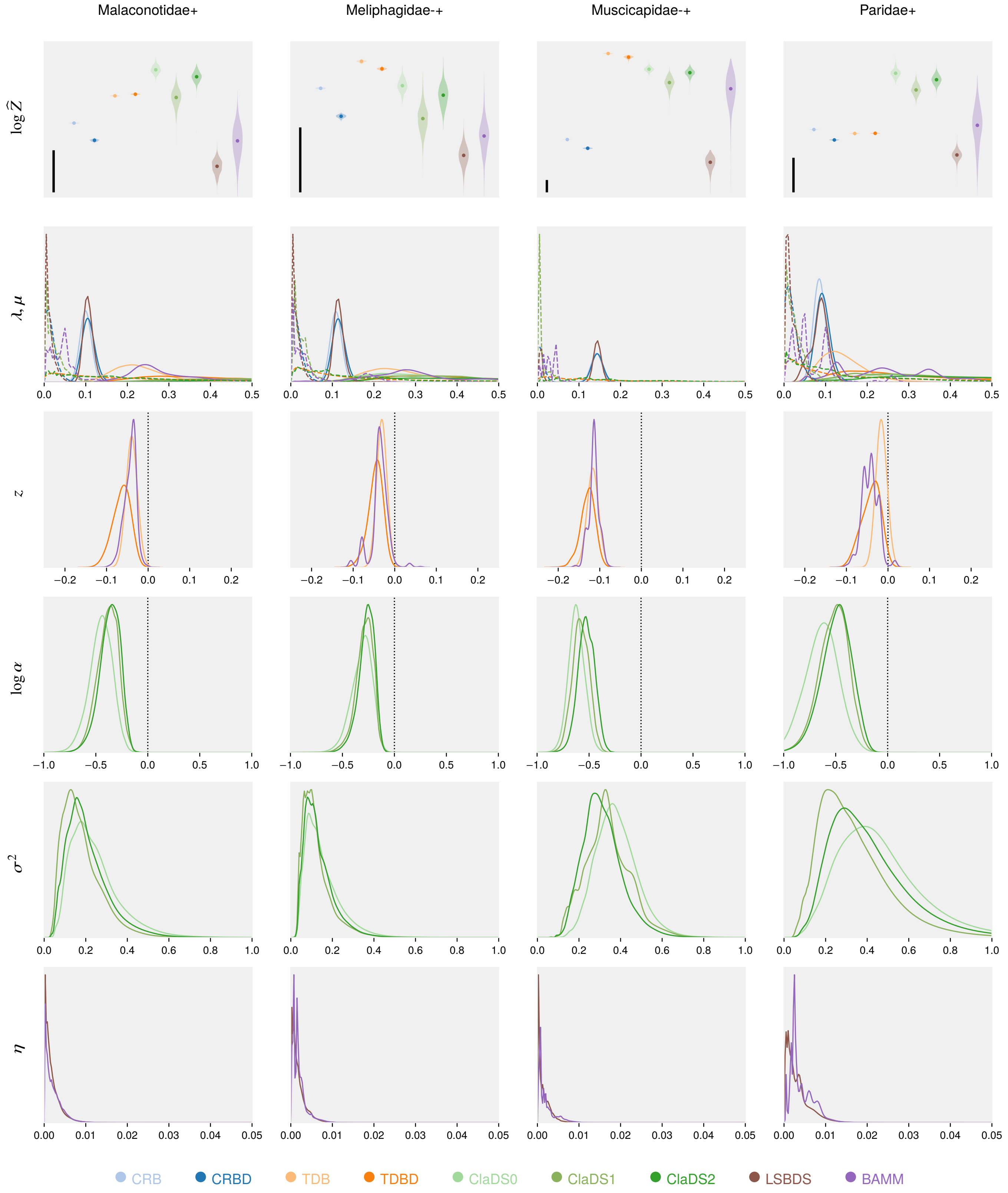


**Fig. 17** Normalization constants and parameter estimates for Malaconotidae+, Meliphagidae-+, Muscicapidae-+, Paridae+.

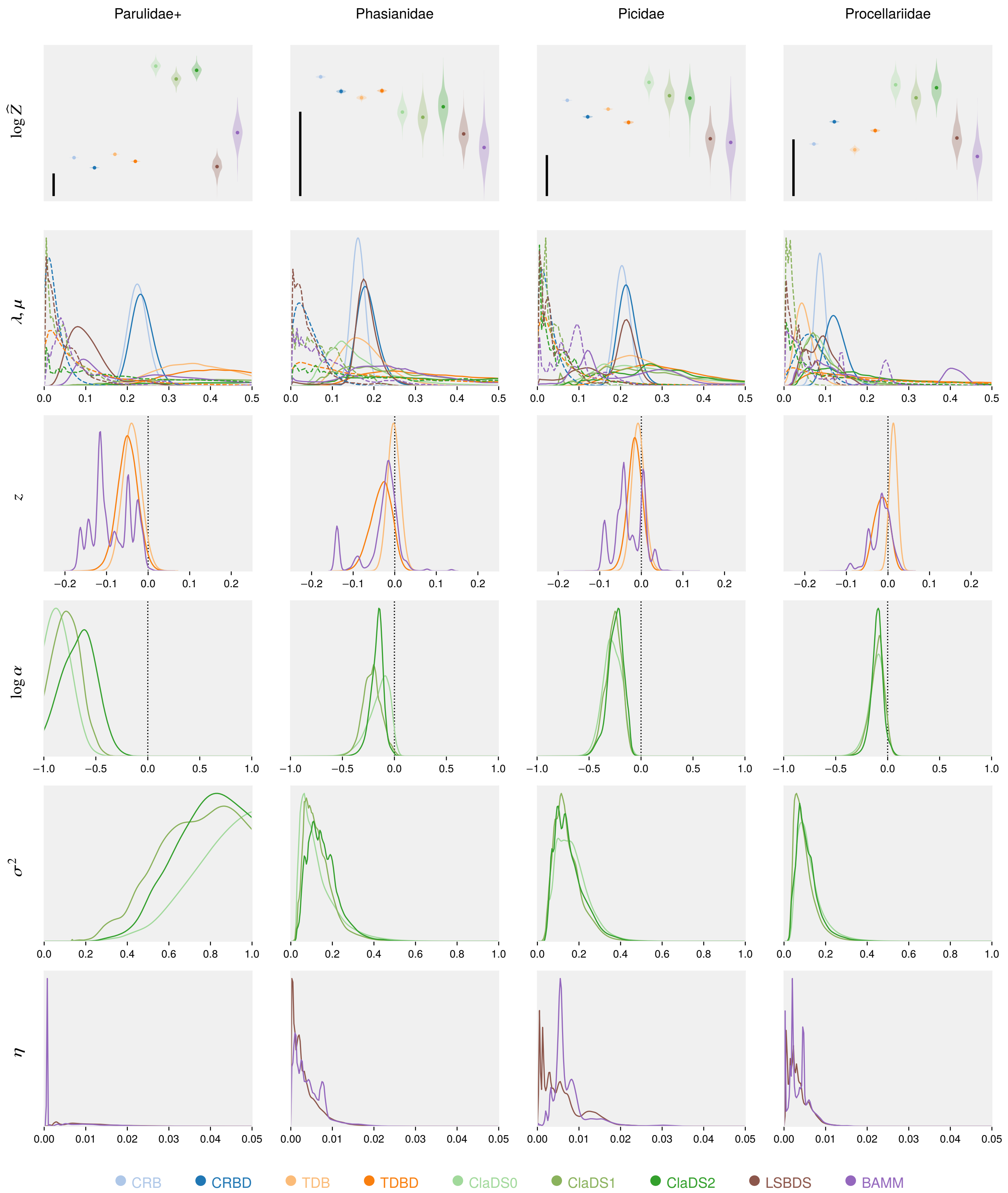


**Fig. 18** Normalization constants and parameter estimates for Parulidae+, Phasianidae, Picidae, Procellariidae.

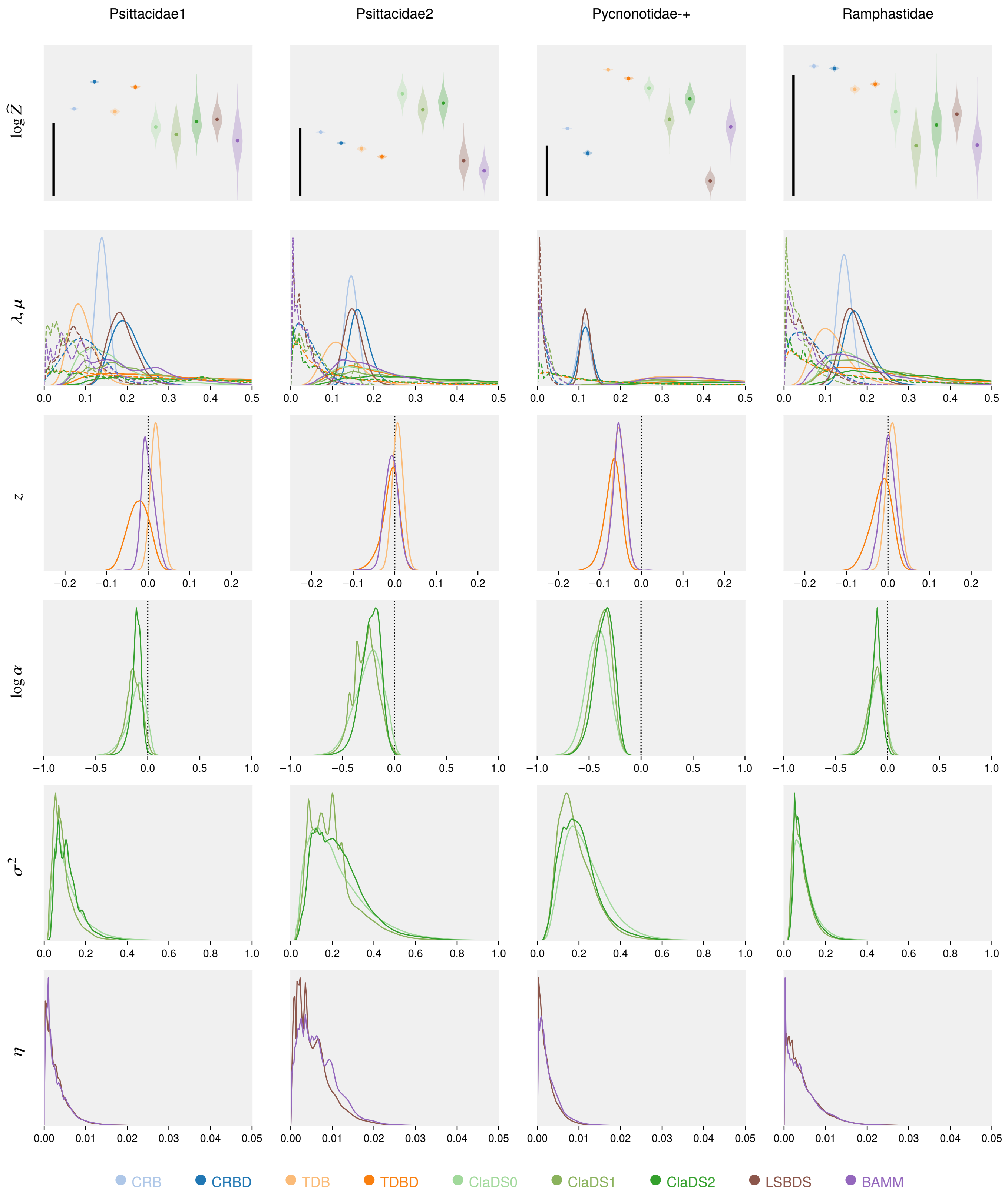


**Fig. 19** Normalization constants and parameter estimates for Psittacidae1, Psittacidae2, Pycnonotidae-+, Ramphastidae.

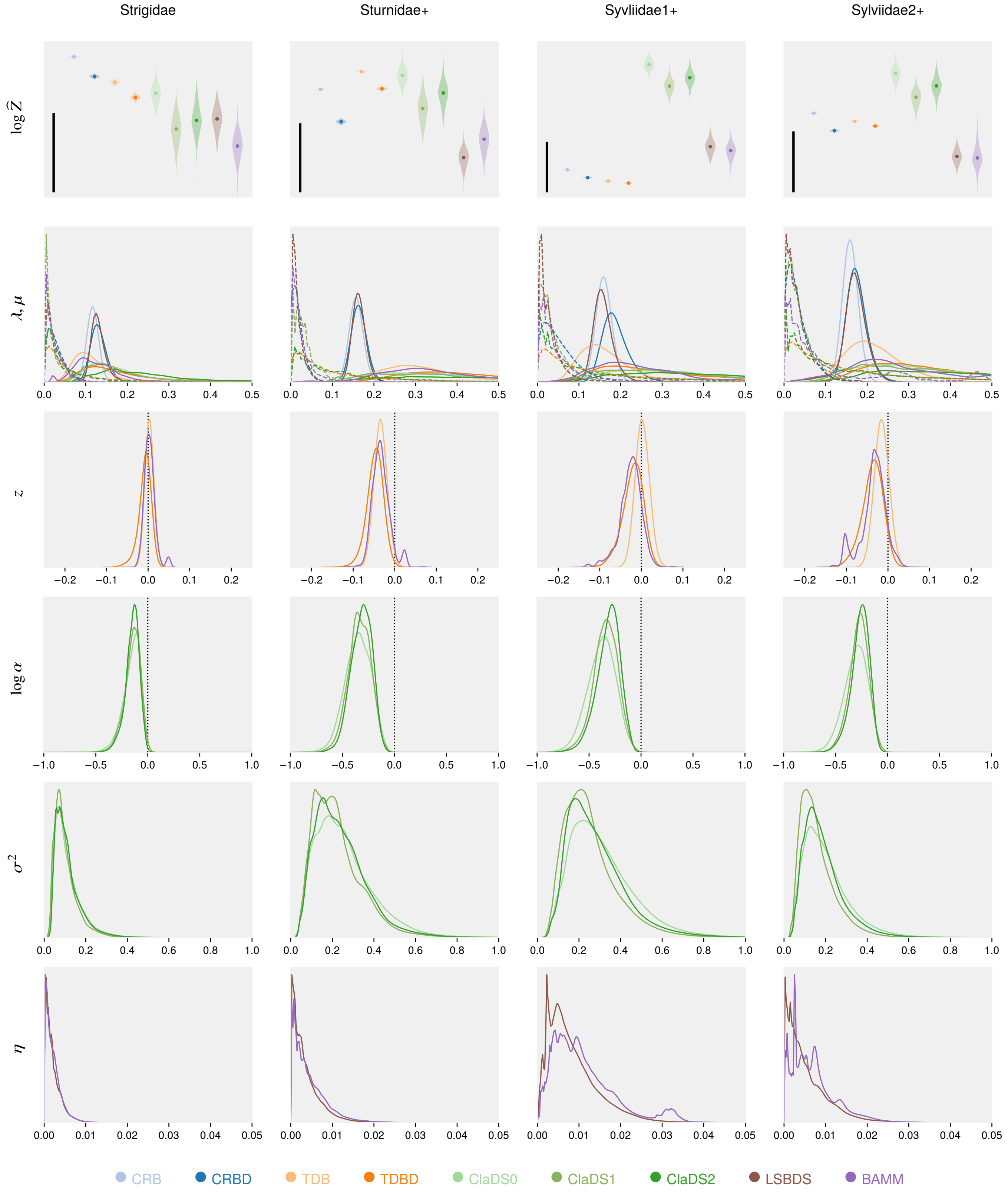


**Fig. 20** Normalization constants and parameter estimates for Strigidae, Sturnidae+, Syvliidae1+, Sylviidae2+.

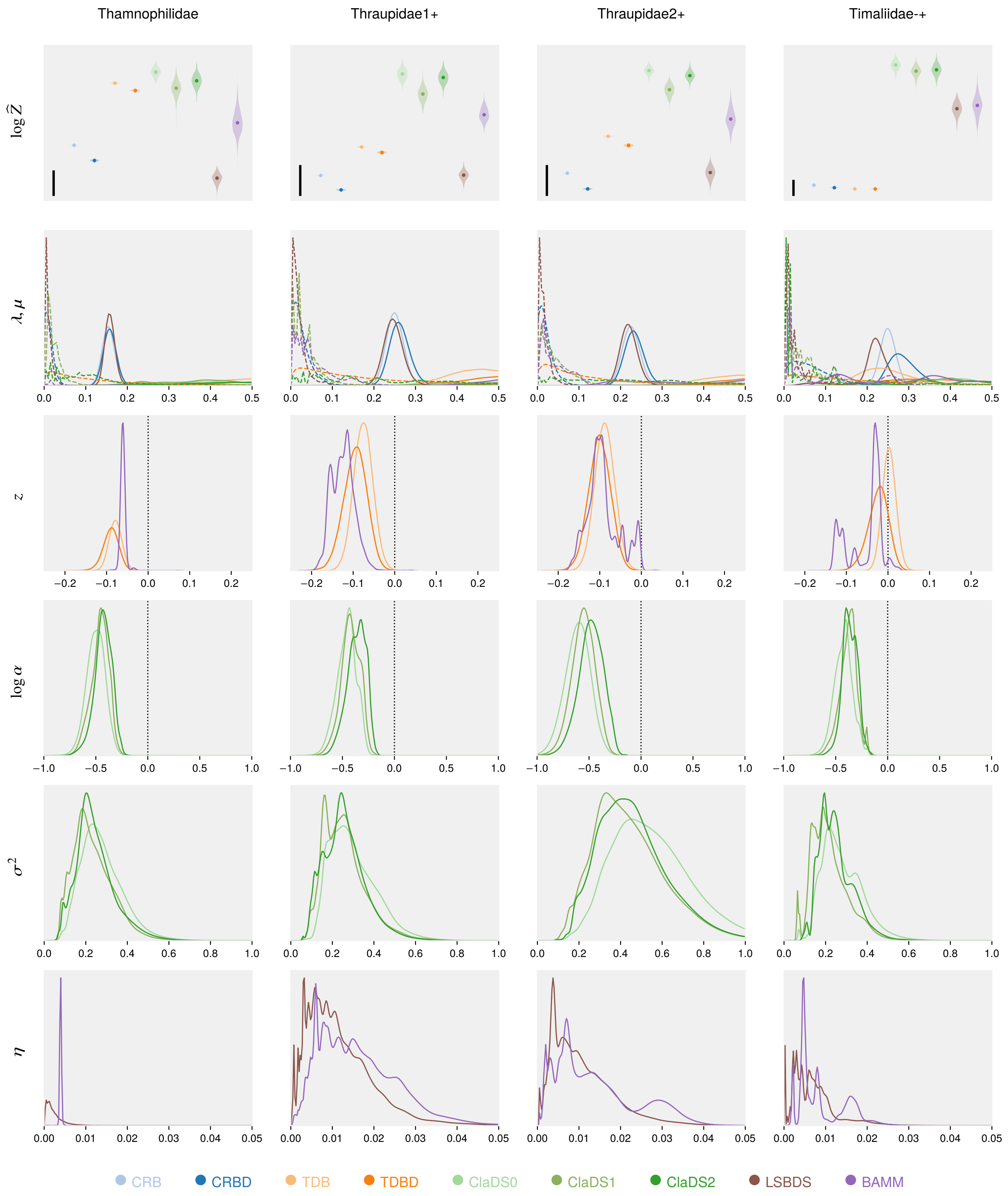


**Fig. 21** Normalization constants and parameter estimates for Thamnophilidae, Thraupidae1+, Thraupidae2+, Timaliidae-+.

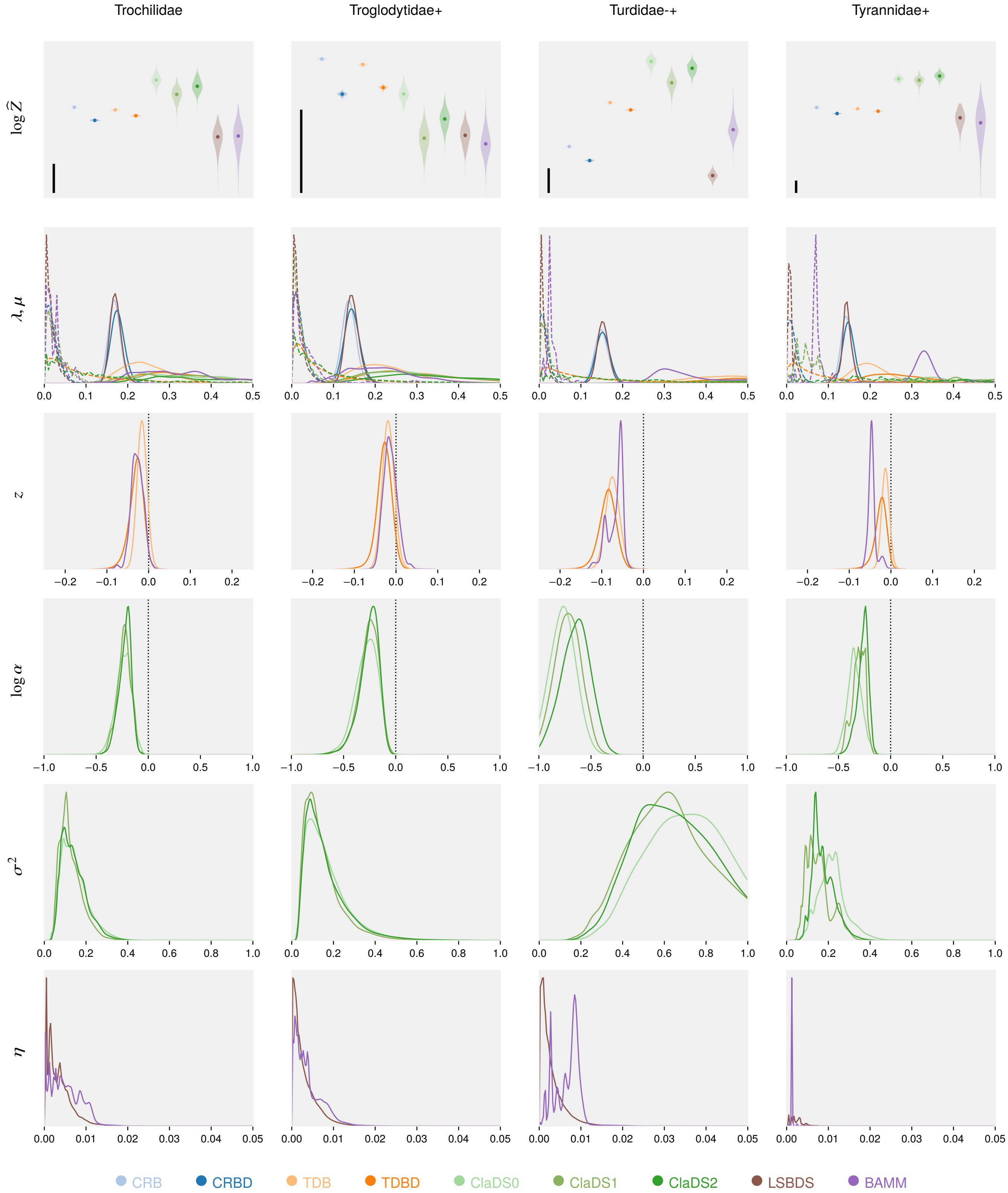


**Fig. 22** Normalization constants and parameter estimates for Trochilidae, Troglodytidae+, Turdidae-+, Tyrannidae+.

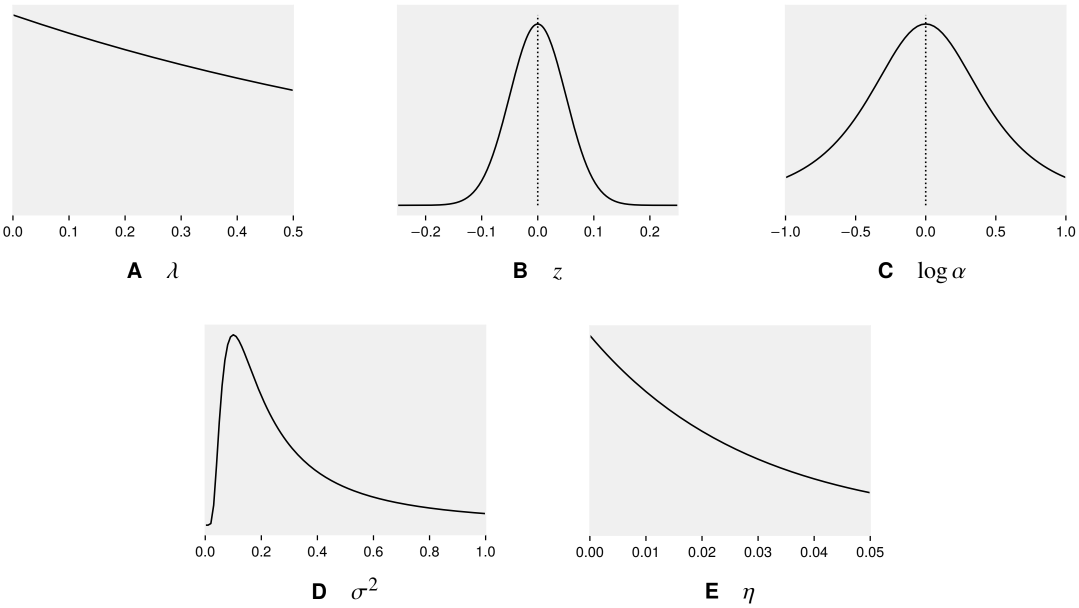


**Fig. 23** Prior distributions plotted for the same region of parameter space used for the posterior distributions in Supplementary Figures 13–22.

models (either CRB(D) or TDB(D)) have the best normalizing constants. There are also some cases where the TDB(D) models do clearly better than the other models, significantly so in a couple of cases (Emberizidae- and Muscicapidae-+, Supplementary Figures 15 and 17, respectively).

There is a clear correlation between the size of the tree and the outcome of the model comparison. Of the trees with less than 100 leaves, the CRB(D) models adequately describe the diversification process in a majority of cases, as indicated by Bayes factors. The largest trees lacking strong evidence of lineage-specific or slowing diversification have around 130 leaves (Columbidae, Cuculidae, Phasianidae and Sturnidae+; Supplementary Figures 14, 15, 18 and 20, respectively). Above that size, all trees bear a clear mark of lineage-specific or, at least, slowing (TDB(D)) diversification. The age of the tree appears to be positively related to the adequacy of simple diversification models. Of the ten youngest trees, only two (Estrildidae+ and Icteridae; Supplementary Figures 15 and 16, respectively) lack strong support for lineage-specific or slowing diversification, while this is fairly common among the oldest trees.

These patterns appear to be best explained by the density of branching events in the reconstructed tree. The more branching events there are per unit time, the more likely it is that the evolutionary process has left signs of density-dependent or lineage-specific diversification. These fluctuations in diversification rates may tend to even out over longer time scales, as old and species-poor trees are often adequately explained by simple models. However, there are clear exceptions. For instance, the Paridae+ (Supplementary Figure 17) shows clear evidence of lineage-specific diversification, despite being an old group (40.9 Ma) with relatively few species (55).

Overall, our results clearly illustrate that Bayes factors are inherently conservative, preferring simpler models unless the signal in the data is sufficiently strong to decisively reject them. While this could be considered a reasonable feature, some caution is nevertheless needed when interpreting the outcome of the model comparison experiment. In particular, the fact that very simple models seem adequate for so many of the bird clades appears to largely reflect the lack of (sufficiently strong) evidence and should not be interpreted as evidence of absence. Many of the trees analysed here (and elsewhere) are too small or not informative enough to allow for a non-trivial outcome.

The degree to which Bayes factors are conservative is dependent on the prior distributions used for the additional parameters of the more complex models. More diffuse priors automatically result in higher penalties in the model comparison—and this even if the posterior distribution itself is not impacted or only marginally impacted. Often, this is not a major problem, for instance when comparing models of sequence evolution, where the signal contributed by the sequence data easily overwhelms the penalty induced by diffuse priors. Here, in contrast, the empirical signal contributed by phylogenetic trees of surviving lineages about the underlying diversification process is somewhat weaker, making the relative impact of the prior on the outcome of Bayesian model tests more substantial.

Whether our priors strike a reasonable balance between simple and complex models is, of course, open to discussion. We note, however, that our priors for the ClaDS models are less conservative than the ones proposed originally for these models[30]. Thus, our priors penalize the ClaDS models less than would otherwise have been the case. We also want to re-emphasize that, to allow fair model comparisons, we chose priors on analogous model parameters that were similar, if not identical, across models.

An alternative to Bayesian model comparison is to focus on model adequacy, that is, the extent to which the models are consistent with the data. A popular approach to assess model adequacy is to use posterior predictive checks, but this requires the specification of an appropriate discrepancy measure[48]. A general criterion of model adequacy that avoids this difficulty is the recently introduced data consistency criterion[49]. However, we refrain from pursuing this topic further here.

#### 10.2 Robustness of complex models

An important result that emerges from our analyses, and that we want to emphasize, is the robustness of complex diversification models. Even when Bayes factors indicate that simple models are adequate, the more sophisticated models often give consistent estimates for the additional model parameters. Good examples are provided by the posterior estimates for $\sigma^2$, describing the rate of gradual, lineage-specific change in diversification rates in the ClaDS models, and $\eta$, denoting the rate of punctuated change in the LSBDS and BAMM models; both of these parameters are usually estimated to be close to 0 when simple models appear adequate. Similarly, the parameters related to potential density-dependent effects ($z$ for TDB(D) and BAMM, and $\log \alpha$ for ClaDS) are often close to 0 when the CRB(D) models have the best marginal likelihoods. Furthermore, no-extinction models , such as ClaDS0, often have higher marginal likelihoods than their counterparts that accommodate extinction. However, in these cases, the more complex models almost always estimate extinction or turnover rates that are close to 0. This usually occurs with very little impact on the estimation of other parameters, as is well illustrated by the very similar posterior distributions obtained across the ClaDS model series, despite the fact that ClaDS0 often has (slightly) better marginal likelihood than the more complex variants.

If the results are scrutinized, one discovers that the advanced diversification models actually appear to pick up weak but consistent signal for more complex patterns even when they are not favored by the model tests. For instance, when posterior estimates of $\log \alpha$ or $z$ are significantly different from 0 in these cases, the estimates always suggest slowing diversification rates, and the models that accommodate such variation over time tend to be the ones with the best model likelihoods, even if they are only marginally better than the constant-rate models. Taken together, these observations suggest that the more complex models might in fact be generally more adequate than the simpler ones. The risk of obtaining erroneous or misleading inference under more complex models appears to be low, at least in comparisons among nested models with similar dimensionality.

#### 10.3 Slowing diversification rates

The strongest signal across bird clades in our analyses is undoubtedly the support for slowing diversification rates. This is seen already in the model comparisons but perhaps more clearly in the posterior estimates of $\log \alpha$ in the ClaDS models, and $z$ in the TDB(D) and BAMM models (Supplementary Figures 13–22). The estimates are almost universally below 0, indicating decelerating rates, and usually significantly so (more than 95% of the credible interval on negative values). Nowhere is the signal more evident than in the four bird clades where the models that only account for changing diversification rates over time—the TDB(D) models—come out distinctly ahead of all others in the model comparison (Emberizidae-, Meliphagidae-+, Muscicapidae-+ and Pycnonotidae-+). In two of those cases (Emberizidae- and Muscicapidae-+; Supplementary Figures 15 and 17, respectively), the Bayes factors even provide strong evidence in favor of TDB(D) over all other models.

Diversification rates that slow down over time are usually attributed to competition for limited resources or niches[50,51,52]. Alternative explanations that have been proposed include: (1) subdivision of geographic ranges at speciation; (2) speciation bursts driven by environmental or geological change; (3) failure to keep pace with environmental change; and (4) protracted speciation (related to the diversified sampling bias, see below)[53]. It might be possible to tease apart some of these factors by developing more sophisticated diversification models within the PPL framework, but this is outside the scope of the current paper. Regardless of the causes, it is clear that there is a strong signature of slowing diversification rates in the bird clades, and that it is important to account for this in diversification models.

#### 10.4 Gradual change, punctuated change or both?

Unsurprisingly, there is also clear evidence of variation across lineages in diversification rates. Of the 40 bird trees, Bayes factors strongly favor models accommodating lineage-specific effects over simpler ones in 15 cases. Even in the remaining cases, there is often some support for lineage-specific variation in diversification rates, as indicated by posterior estimates of model parameters.

The ClaDS models consistently explain this variation in diversification rates better than the LSBDS and BAMM models. In fact, there are only three groups for which the LSBDS and BAMM models are strongly favored over the corresponding simple models: Anatinae, Lari, and Timaliidae-+ (Supplementary Figures 13, 16 and 21, respectively), The BAMM model also does comparatively well on the Thraupidae1+ tree (Supplementary Figure 21). As expected, these trees are also associated with posterior estimates of $\eta$ that differ substantially from 0. However, even for these trees, where BAMM and LSBDS detect major shifts in diversification rates, the ClaDS models provide a better fit to the data.

We may conclude that lineage-specific differences in diversification rates are better explained by slow, gradual changes, which accumulate over time, than by a few events that drastically alter the rates. One possible explanation for this is that the punctuated models (BAMM and LSBDS) draw the new $\lambda$ and $\mu$ (and $z$ for BAMM) values from diffuse priors at process switching events. This means that they carry heavy penalties in Bayes factor comparisons; the more switches there are, the heavier the penalty against these models. An interesting difference between the punctuated models and the gradual models is that the former allow both $\lambda$ and $\mu$ to vary over the tree, while only $\lambda$ is modulated over the tree in the latter. Could this be the explanation for the gradual models outperforming the punctuated models? We tested this by modifying LSBDS and BAMM such that they assumed a constant turnover rate ($\epsilon = \mu/\lambda$), as in ClaDS2, and only varied $\lambda$ (and $z$ for BAMM) at switching points. We then re-computed the normalization constants for Lari, one of the clades with the strongest evidence for major shifts in diversification rates. The normalization constants of the punctuated models did not improve noticeably due to this modification (results not shown). This finding suggests that the strong evidence in favor of gradual over punctuated change is not due simply to the punctuated models postulating changes in extinction rates that are not supported by the data.

A fascinating question is whether there remains any evidence for occasional major shifts in diversification rates if one first adequately accounts for the strong underlying signal of slow and gradual change. This can now be examined by extending the PPL framework we present here to diversification models that combine ClaDS-like and BAMM-like features. Given the general lack of support for radical shifts in diversification rates across the bird trees, it seems likely that such shifts are rare, if they occur at all. Therefore, identifying them would presumably require analyses of larger trees than the ones examined here. However, it also seems likely that the two processes interact, such that it becomes more difficult to detect major shifts when the gradual changes are not accounted for. Thus, it is possible that there are major shifts in the bird trees that our analyses failed to detect because of shortcomings in the models. We will have to await future analyses using more sophisticated models before we know whether this is the case.

#### 10.5 Discretizing punctuated-change models

Computing likelihoods for punctuated models of diversification by integrating out the rate priors using discrete approximations is potentially a very powerful approach[29]. It allows for robust and computationally efficient MCMC inference, as long as a small number of rate categories yield sufficiently accurate likelihood estimates. This decidedly appears to be the case, especially if only changes in speciation rate are modeled; empirical analyses suggest that ten categories is quite sufficient for most problems[29].

Unfortunately, it is difficult to see how this approach can be extended to accommodate slowing (or increasing) diversification rates over times as in BAMM, because then it would be necessary to integrate out an infinite number of rate acceleration or deceleration processes with different starting points. This appears to be an important limitation from an empirical perspective, at least judging by the bird trees we analyzed. The LSBDS model does not fit many of the reconstructed trees well; this is undoubtedly linked to the substantial support for slowing diversification rates in most bird clades. If we restrict our attention to models of punctuated change, we find 8 trees with strong evidence favoring the BAMM model over the LSBDS model. There is not a single tree for which the evidence goes strongly in the other direction.

#### 10.6 Sampling biases

Some of the results that emerge from our analyses are probably due, at least in part, to sampling biases. The lack of evidence for extinction rates above zero is an obvious case. Models without extinction (CRB, TDB, ClaDS0) almost always do better than models with extinction (Supplementary Figures 13–22). The most notable exception is the Lari (gulls), where the CRBD and TDBD models significantly outperform their non-extinction counterparts (Supplementary Figure 16). A similar but much weaker signal is seen in a few other groups: the Anatinae (Supplementary Figure 13), Charadrii (Supplementary Figure 14), Procellariidae (Supplementary Figure 18) and Psittacidae1 (Supplementary Figure 19). The outcome of the model comparison is generally consistent with posterior estimates of $\mu$ under the models that do accommodate extinction (CRBD and TDBD). That is, estimated extinction rates are usually low except for Lari, and to a lesser extent for the other groups that weakly favor models that accommodate extinction. Interestingly, Lari is also unusual in that there is evidence for accelerating speciation rates ($z > 0$). However, this occurs only in the TDB model, and is probably an artefact of not accounting for extinction, as extinction rates noticeably above zero are expected to lead to an apparent acceleration of speciation rates close to the present in reconstructed trees[25].

The lack of support for extinction in our analyses is consistent with results from previous diversification studies[52]. Given the overwhelming evidence for frequent extinction in the fossil record, these results are not plausible. Clearly, current phylogenetic diversification analyses tend to underestimate extinction rates. An important factor that may contribute toward such underestimation of extinction rates is the *diversified sampling* bias[54]. This is the tendency of biologists to systematically select leaves of phylogenetic trees in such a way that the diversity represented in the sampled tree is maximized, instead of choosing leaves randomly as assumed by standard birth-death models. The diversified sampling bias will lead to pruning of the most recent splits from the complete reconstructed tree. The more incomplete the sampling is, the deeper the period that is devoid of splits will extend into the past. In the most extreme cases, the sampled tree will look like a bush: most of the splits will be close to the root of the tree, and all the leaves will sit on long terminal branches. If one analyzes a tree sampled to maximize diversity under a model assuming random sampling, extinction rates can be substantially underestimated[54]. Failing to account for diversified sampling bias can also result in severe biases in divergence time estimates[55,56].

The bird trees we examined here represent fairly complete samples of extant species (from about half of the species and up), with no obvious biases (Table 6). Nevertheless, it is easy to show that they are characterized to some extent by diversified sampling. In total, the bird data include 9,993 species, 6,670 of which were sampled for sequencing[44]. Only the latter were included in the trees we examined here. If the sequenced species were chosen randomly, then they would include approximately the same number of genera as any random sample of 6,670 species. However, the sequenced species cover as many as 1,880 genera, while 10,000 random samples of 6,670 species included only 1,759 genera on average (range 1,703 to 1,808). Thus, the sequenced set has significantly fewer species per genus than expected by chance, the type of bias we would predict from diversified sampling.

A similar bias that may also affect our results is that incipient species, sibling species and subspecies are likely to be underrepresented in the data. Some of those lineages will eventually develop into true species, and should be included in estimates of speciation and extinction rates at the species level. Unacknowledged diversified sampling around or just above the species level is linked to the phenomenon of *protracted speciation*[57].

Interestingly, diversified sampling bias that is not accounted for correctly could also contribute to the strong support for slowing diversification rates, even though there are also plausible biological explanations for such patterns[53]. To understand why diversified sampling may give the impression of slowing diversification, consider that the extinction rate estimates are largely based on the apparent acceleration of speciation towards the recent seen in surviving trees because there has not been time enough for extinction of the side lineages that will eventually disappear[25]. Diversified sampling would systematically remove the evidence for this apparent acceleration in speciation rates.

As one might expect, there is also a link between the posterior estimates of extinction and the evidence for slowing diversification rates. Specifically, models that accommodate slowing diversification rates (TDBD, ClaDS models and BAMM) are also associated with distinctly higher estimates of initial extinction rates than models that do not (LSBDS, CRBD) (Supplementary Figures 13–22). How this link might be affected by diversified sampling biases is currently unclear. Pursuing this topic further would be outside of the scope of the current paper. However, we do note that it is relatively straightforward to modify our script to account for diversified sampling according to the model suggested by Höhna et al.[54], or potentially even more realistic models.

Some recent papers have suggested that unrealistically low extinction rate estimates may be the result of applying models that do not account for the heterogeneity across lineages in diversification rates that characterize most phylogenetic trees[58,31]. If this were true, then one would expect extinction rate estimates in our analyses to be higher for models that accommodate lineage-specific variation than for comparable models that do not. However, this is not the case. For instance, extinction rate estimates are similar or lower for LSBDS than for the comparable CRBD model (Supplementary Figures 13–22). Similarly, the estimates are similar or lower for ClaDS1 than for CRBD. The initial extinction rate estimates of the ClaDS2 and BAMM models are best compared to the analogous estimates for the TDBD model, which does not accommodate variation across lineages in diversification rates. These estimates are similar for most bird trees (less clear for BAMM than for ClaDS2), further supporting the conclusion that unaccommodated heterogeneity in diversification rates is not responsible for the low extinction rate estimates. Of course, we cannot exclude the possibility that even more sophisticated models than the ones examined here could show that extinction rates are underestimated because of shortcomings in the modeling of across-lineage variation in diversification rates.

#### 10.7 Statistical power

Reconstructed trees carry only a limited amount of information about absolute speciation and extinction rates. This is illustrated well by the underestimation of extinction rates discussed above. For really powerful analyses of diversification processes, we need trees that include data from the fossil record, that is, observations of both extinct and extant lineages[59]. In most cases, however, observation of extinct lineages is not possible, or the information about extinct lineages is bound to be very incomplete, so we need to extract as much information as possible from trees that only (or mainly) comprise surviving lineages. There are several ways in which analyses of such trees can be improved. Addressing sampling biases appropriately would be an important step in the right direction. Making the model of the diversification process itself more realistic, for instance by combining gradual and punctuated change as suggested above, would be another. However, the most obvious way to improve the analyses would be to increase the amount of data.

A natural way of extending the present work in this direction would be to opt for a hierarchical model-averaging approach, in which all trees in a set, such as the bird clades, would be analyzed simultaneously. Specifically, each tree would randomly choose from a mixture of all available models, while the mixture proportions and the hyper-parameters tuning the priors over model-specific parameters would themselves be estimated across trees, using a hierarchical modeling design. Global estimation of hyper-priors and mixture weights based on the whole collection of trees is an efficient way to fit the priors to the true prevalence of alternative modes of diversification and the true variation in parameter values present in the data and therefore should result in well-calibrated model selection. Joint analysis of all trees would also make it possible to collect the weak signals disseminated across the many small trees of the analysis. Such developments are exactly what the probabilistic programming framework introduced here is meant to facilitate.

A completely different approach would be to analyze larger portions of the tree of life. For instance, our analyses of 40 bird clades could have been replaced with a single analysis of the entire bird tree, doubling the coverage of bird species (from 5,000 to 10,000). Such an analysis would not only include more of the variation seen across terminal clades, it would also add data on the deeper splits in the tree. These splits could potentially inform the model about long-term macroevolutionary patterns that could not be detected in analyses of only terminal clades, regardless of how sophisticated. For instance, it has been suggested that the bird radiation as a whole is characterized by rare but major boosts in diversification rates, presumably linked to key innovations opening up new ecological niches[44]. If these upward jumps in diversification rates are substantial enough, it could explain why there is overwhelming support for slowing diversification rates in individual bird clades, even though the rates appear to be accelerating over the bird tree as a whole[44]. Such a mega-analysis would have to be based on a model that is more sophisticated than the ones explored here. Minimally, it would have to account for both gradual and punctuated change in diversification rates. Ideally, it would also account for variation across lineages in the strength of the slowing forces on diversification, and in the rate of gradual change in speciation and extinction rates. Again, such developments are well supported by the probabilistic programming framework, although it is still an open question whether current inference strategies are efficient enough or whether further refinement is needed.
